# Nociceptive-evoked electroencephalography can predict inter-individual differences in high frequency induced sensitisation of central nociceptive pathways: A test-retest reliability study

**DOI:** 10.64898/2026.10.31.763300

**Authors:** Samuel Mugglestone, Sam Hughes, Giorgio Ganis

**Author notes:** Corresponding author and address: Dr Giorgio Ganis, School of Psychology, Faculty of Health, University of Plymouth, Plymouth, Exeter, Devon, United Kingdom, PL4 8AA. co-senior authors (Sam Hughes, Giorgio Ganis). Data availability statement: Data are available upon request.

## Abstract

Evoked-EEG can capture changes in cortical responses during experimentally induced sensitised pain states. However, the test-retest reliability of such methods requires validation. The present study applied HFS across two identical sessions in the same individuals to assess the test-retest reliability of EEG dynamics evoked by a 128mN pin-prick stimulator and single-pulse electrical stimuli before and after HFS conditioning. The value of these features for reflecting individual differences in the development of hyperalgesia was also assessed. Results demonstrate poor-to-moderate test-retest reliability and relatively wide agreement at baseline for all features, with lateralised differences often present. This was further reduced when post-HFS changes were considered. α-ERD assessed from contralateral central electrodes was significantly strengthened post-HFS for both electrical and pinprick stimuli, consistent with increased sensorimotor engagement in response to hyperalgesic and non-hyperalgesic stimuli. Notably, baseline α-ERD was related to the subsequent time course of SHA development. Furthermore, whilst VP magnitude displayed a significant relationship with the general intensity of perceived pain, no specificity for SHA development was evident. Alternatively, θ-ERS captured a general habituation effect relative to variations in reported pain intensity between sessions for both stimuli modalities. These findings indicate that evoked EEG responses may be useful for investigating individual differences in the cortical processing of acute pain and the development time course of HFS-induced sensitivity yet may not possess sufficient test-retest reliability for interventional testing.

## Introduction

Evoked electroencephalography (EEG) provides high temporal resolution for characterising the cortical processing of nociceptive information (Lenoir et al., 2020; Nierula et al., 2024). Combined with experimental pain models such as high-frequency stimulation (HFS), EEG can investigate mechanisms of maladaptive pain processing (Bannister & Hughes, 2023; Leone et al., 2025; Ǫuesada et al., 2021). However, whilst previous research has investigated established sensitised pain states, the development time course of HFS-induced changes to evoked EEG remains poorly characterised (Gousset et al., 2023; Leone et al., 2025; van den Broeke et al., 2020; van den Broeke et al., 2017; Van den Broeke et al., 2016; van den Broeke et al., 2015). Furthermore, although the test-retest reliability (Triccas et al., 2022) and individual variability (Hu & Iannetti, 2019; Tiemann et al., 2026) of nociceptive-evoked EEG have been assessed, no such investigation has been conducted within an experimentally sensitised pain state (Leone et al., 2025).

HFS induces dorsal horn plasticity (Klein et al., 2004; Klein et al., 2008; Pfau et al., 2011; van den Broeke et al., 2015). Heterotopic sensitivity reflects secondary hyperalgesia (SHA) which can emerge as early as 15 minutes post-HFS and persist for several hours, with good test-retest reliability at later time points (e.g., > 30 minutes post-HFS) (Fawsitt-Jones et al., 2024; Klein et al., 2004; Klein et al., 2008; Lebrun et al., 2025; Leone et al., 2025; Pfau et al., 2011; Ǫuesada et al., 2021). However, we have previously demonstrated that the time course of SHA development varies across repeated test sessions and is not directly attributable to typically considered psychophysical factors, such as the state of the descending pain modulatory system or pain during HFS-conditioning (Mugglestone et al., 2026). Here, we extend these findings by investigating time-locked EEG dynamics between repeated test sessions to better understand cortical mechanisms underlying variability in SHA development.

Nociceptive stimuli evoke multiple event-related spectral perturbations (ERSPs) across frequency bands (Diers et al., 2020; Gousset et al., 2023; Hauck et al., 2015; L Hu et al., 2014; Hu & Iannetti, 2019; Hu et al., 2013; Jin et al., 2018; Liberati et al., 2016; Linde et al., 2023; Nickel et al., 2017; Ploner et al., 2017; Schulz et al., 2015; Tiemann et al., 2026; van den Broeke et al., 2017; van den Broeke et al., 2019; Wang et al., 2023; Yue et al., 2025; Zhang et al., 2021; Zidda et al., 2024). Of primary interest are α-band event related desynchronisations (α-ERD) which comprise at least two components (Hu et al., 2013): reflecting somatosensory activity at contralateral central electrodes (Hu et al., 2013; Ploner et al., 2017), and endogenous task-related processes at parietal occipital electrodes (Hu et al., 2013; Wang et al., 2023). These components reveal the feedback processing of sensory input rather than pain intensity (Ploner et al., 2017), although somatosensory α-ERD can reflect intra-individual variations in pain perception (Tiemann et al., 2026).

Similarly, θ-ERS is not pain-specific and reflects early-latency neural activity from multiple cortical generators (Hu & Iannetti, 2019; Michail et al., 2016; van den Broeke et al., 2017; Zhang et al., 2021), including lemniscal and extralemniscal responses (Somervail et al., 2025). Thirdly, γ-ERS has been found to preferentially encode pain intensity, reflecting the spiking of somatosensory interneurons (Hu & Iannetti, 2019; Yue et al., 2025). However, this is predominantly exhibited within thermonociceptive modalities, and recent investigations of MPS have failed to identify significant γ-band fluctuations post-HFS (Gousset et al., 2023).

Additionally, sudden and isolated sensory stimuli (SISS) can evoke a biphasic vertex potential (VP) (Gousset et al., 2023; Hu & Iannetti, 2019; Perovic et al., 2025; Somervail et al., 2025; Somervail et al., 2021; Tiemann et al., 2026; E. van den Broeke et al., 2021; van den Broeke et al., 2020; van den Broeke et al., 2017; Van den Broeke et al., 2016; van den Broeke et al., 2015; Yue et al., 2025; Zhang et al., 2021), potentially indicating extralemniscal thalamocortical activity (Perovic et al., 2025; Somervail et al., 2025; Somervail et al., 2021). Within experimentally sensitised pain states, increased VP amplitudes have been consistently reported (Gousset et al., 2023; E. van den Broeke et al., 2021; van den Broeke et al., 2020; van den Broeke et al., 2017; Van den Broeke et al., 2016; van den Broeke et al., 2015). Although VP amplitude is modulated by stimulus intensity (Iannetti et al., 2005; van den Broeke et al., 2015), modality (Linde et al., 2023; Zhang et al., 2021), and laterality (Zhang et al., 2021), marked increases are most evident when perception changes categorically from unperceived to perceived (Lee et al., 2009) or sub-nociceptive to nociceptive (van den Broeke et al., 2015), consistent with allodynia-like sensitivity rather than hyperalgesia (Leone et al., 2025). Because VP amplitude reflects stimulus salience relative to the pre-stimulus state (L Hu et al., 2014; Hu & Iannetti, 2019; Hu et al., 2010; Iannetti et al., 2005; Iannetti et al., 2008; Jin et al., 2018; Lee et al., 2009; Somervail et al., 2025; Somervail et al., 2021; Squintani et al., 2021; Valentini et al., 2012; Wang et al., 2023; Zhang et al., 2021; Zidda et al., 2024), it may index the facilitation of ascending nociceptive processing post-HFS.

The present study examined the test-retest reliability of nociceptive-evoked EEG for characterising hyperalgesia development following HFS. We hypothesised that hyperalgesia would be associated with increased VP amplitude, reflecting greater extraleminscal activation. We further expected greater α-ERD over somatosensory electrodes and enhanced θ-ERS, consistent with increased spinocortical afferent drive over the HFS time course. It was hypothesised that baseline individual differences in these characteristics would predict variability in the early temporal dynamics of hyperalgesia development.

## Methods

### Pre-registration

This study is part of a wider research program investigating the reliability of psychophysical and neurophysiological methods for assessing temporal changes to central nociceptive function induced by HFS conditioning. The overall study protocol was preregistered on the Open Science Framework (OSF) and is available at: https://doi.org/10.17605/OSF.IO/S5A9Y.

One preregistered hypothesis aimed at supplementing the limited evidence of a prolonged positive deflection evident at Pz from 300ms onwards arising after the establishment of SHA, which may reflect the sustained synchronisation of neural generators as the product of an altered sensitivity state (Van den Broeke et al., 2016; van den Broeke et al., 2015). However, no such component was evident within the present data.

### Participants

Recruitment was conducted via the University of Plymouth’s Paid Participation Pool alongside word of mouth. Participants were instructed to abstain from alcohol and drugs prior to each visit, and to specifically avoid taking painkillers and to limit caffeine intake. At the start of each visit compliance was confirmed. All 35 recruited participants (final sample size = 27) provided written informed consent and were informed of their continual right to withdraw. The study was approved by the University of Plymouth Ethics Committee (ID:3777) and in compliance with the Declaration of Helsinki (Association, 2013).

Exclusion criteria resulted in the removal of one participant: no history of neurological conditions, long-term health conditions, chronic pain, psychiatric conditions, steroid use, inflammatory disease, or immunosuppression; no allergies or severe skin conditions that may affect the methods or areas of testing; not currently pregnant.

### Experimental design

Two functionally identical sessions were conducted at least a week apart with the aim of assessing the between-session reliability of the temporal changes in EEG features following HFS-conditioning. Various behavioural data were collected at the beginning of each session which are not relevant to the present publication and are discussed separately (Mugglestone et al., 2026). Prior to beginning the main experimental procedures, an EEG electrode array was set up and tested for impedance, before participants were familiarised with the sensory testing stimuli and electrical detection threshold (EDT) was determined over the volar surface of the forearm. Baseline mechanical pinprick sensitivity (MPS) and electrical pain perception (EPP) responses were assessed on the ipsilateral and contralateral forearms, followed by HFS conditioning. MPS and EPP responses were then reassessed at 4 time points post-HFS with 15-minute intervals between the beginning of each block. During stimulus delivery, view of the testing sites was occluded via a screen-box which limited the perceptual field to a computer monitor on which a fixation cross (white) was displayed throughout. Participants were instructed to focus their visual attention onto this fixation cross and to refrain from blinking or moving as much as possible. A chinrest was also provided to reduce muscle-strain artifacts.

Within each block, 20 stimuli were delivered per condition (arm x modality) with an interstimulus interval (ISI) of ∼2 seconds resulting in 40 stimuli per arm, and a total of 80 stimuli within each time point as both mechanical pinprick and single-pulse electrical stimulation were assessed, albeit in separate zones. This was considered the minimum number of trials to effectively characterise event-related EEG features based upon comparable studies (Biurrun Manresa et al., 2018; Gousset et al., 2023; van den Broeke et al., 2020; van den Broeke et al., 2017; Van den Broeke et al., 2016; van den Broeke et al., 2015). Furthermore, although this relatively short ISI may affect the appearance of longer lasting components, the primary analyses conducted within this chapter focused solely upon transient nociceptive-evoked responses.

### High-Frequency Stimulation

Cutaneous electrical stimuli were applied to the right volar forearm (5cm distal to the cubital fossa) using an EPS-P10 electrode (MRC Systems GmbH, Heidelberg, Germany) via a constant current stimulator (2ms pulse width; DS7; Digitimer Ltd; Welwyn Garden City, UK). Prior to attaching the HFS electrode, the skin was cleaned with a non-alcoholic aqueous solution and allowed to dry. The EPS-P10 electrode contains a circular array of 10 cathodal pins (diameter: 0.25mm) and a larger surface area anode (area: 410 mm2). Such concentric surface electrodes have been shown to selectively activate A-delta and & fibre free nerve endings located in the superficial epidermal layers without the excitation of A-beta fibre mechanoreceptors located in the dermis (Medina & Hughes, 2025; Poulsen et al., 2022). Sufficiently high intensity electrical stimuli can bypass peripheral transduction mechanisms directly activating voltage-gated ion channels (Leone et al., 2025). Notably, this specificity may falter depending upon the intensity of stimulation, resulting in the activation of deeper A-beta fibres (Legrain & Mouraux, 2013). However, characterising the incidence of this phenomenon is beyond the scope of the current study.

Electrical detection threshold was determined using 2ms pulses starting at 0.05 mA in 0.05 mA increments until participants reported a clear sensation, then decreased by 0.01 mA until the sensation was lost. The intensity was increased in 0.01 mA increments until the sensation returned, repeating this process three times. The average of the six recorded intensities represented each participant’s EDT. Trains of HFS (100 Hz; NL304 Period Generator; Digitimer Ltd) were applied 5 times to the test (right arm) for 1 second each (NL405 Width Delay; Digitimer Ltd) at 15x EDT every 10 seconds (i.e. over a 50 second period). The participants were asked to rate the intensity of the HFS after each train using a numerical rating scale (NRS; 0 = no pain, 100 = worst pain imaginable).

### Heterotopic mechanical stimulation

Mechanical pain sensitivity (MPS) was assessed across both forearms using a 128 mN pin prick stimulator within a typical heterotopic area laterally adjacent to the conditioning electrode yet within 2 cm as depicted in Lebrun et al. (2025) and Medina and Hughes (2025). However, the area of SHA was not systematically mapped. Although mechanical pinprick stimuli activate both high- and low-threshold mechanoreceptors, low-intensity pinpricks can be perceived as non-painful (Leone et al., 2021). Therefore, the medium range intensity of 128mN was used throughout the study and baseline pain ratings are reported.

Following every five successive stimuli, participants rated their average perceived pain on a numerical rating scale (NRS; 0 - 100). These were further averaged to provide a single value for each time point. Therefore, within each time point, participants received a total of 20 pinprick stimuli to each of the ipsilateral and contralateral arms.

### Homotopic electrical stimulation

Electrical pain perception (EPP) was assessed across both forearms using the same epi-cutaneous pin electrode as used to deliver HFS, with the same intensity of 15x EDT as outlined above. However, each stimulus consisted of a single square wave pulse via a constant current stimulator (2ms pulse width; DS7; Digitimer Ltd; Welwyn Garden City, UK) which were delivered directly to the area of HFS conditioning. Both test- and control-arm electrodes were positioned prior to baseline assessments and remained in place throughout the main experimental procedures 5cm distal to the cubital fossa. Such stimulation can selectively activate A-delta and & fibres through bypassing peripheral transduction mechanisms and directly activating voltage-gated ion channels within superficial afferent nociceptors (Leone et al., 2025).

The rating procedure was identical to that described for mechanical stimulation. Participants provided NRS ratings (0–100) after every five stimuli; ratings were averaged within block and collapsed to a single value per time point. Thus, 20 single-pulse electrical stimuli were delivered per arm at each time point.

### EEG data recording

EEG data were recorded continuously throughout the main experimental procedures using an ActiCAP Slim 64-channel electrode array positioned according to the extended 10-20 system, referenced to FCz and grounded at AFz (band-pass filter = 0.01 to 100 Hz; sampling rate = 500 Hz) using an ActiCHamp Plus amplifier and BrainVision Recorder software (Brain Products GmbH, Glitching-Sȕd, Germany).

### EEG data preprocessing

Preprocessing was conducted using EEGLAB (version 2025.1.0 (Delorme & Makeig, 2004)) and custom functions within MATLAB (version 2024b) as follows:

- Continuous EEG data were band-pass filtered (0.1-100 Hz) using a zero-phase, non-causal, Hamming-windowed FIR filter (transition bandwidth = 0.1 Hz; -6 dB cutoff definition).
- Aberrant channels were identified and removed automatically using the Clean_rawdata plugin (version 2.10) without ASR or window rejection (flatline >= 5 seconds; channel correlation = 0.8; line noise = 4 std; high-pass filter for channel correlation = 0.25 Hz; 0.75 Hz). Channels were also manually inspected and removed when deemed appropriate (Delorme, 2023).
- UK mains interference (50 Hz and its first harmonic at 100 Hz) was removed using the CleanLine plugin (v2.1).
- Aberrant data drifts were identified automatically using the Clean_rawdata plugin without channel or window rejection and artifact subspace reconstruction was applied (maximum acceptable 0.5 second, window deviation = 20 std) (Chang et al., 2018; Delorme, 2023).
- Data were re-referenced to average with care to avoid rank-deficiency issues (Delorme & Makeig, 2023; Kim et al., 2023).
- Independent component analysis (ICA) was conducted using AMICA (version 17 (Hsu et al., 2018)), using a single model with 3000 iterations and all other parameters at default.
- Artifact components classified as muscle, heart, or channel noise were rejected with a threshold of ≥ 0.9 using the ICLabel plugin. Eye-related components were handled separately: components were first labelled “ocular” when the eye class had the highest ICLabel probability (i.e., was the dominant class). Among these ocular-labelled components, only those with a brain-class probability < 5% were removed. This approach targeted ocular artefacts whilst reducing the risk of discarding components with a non-trivial neural contribution, as done in previous work (Hsu et al., 2024).
- Event-related data were segmented into epochs between -950 ms to 1450 ms relative to stimulus onset to allow for an appropriate time frequency resolution [-700 ms to 1200 ms]. Baseline correction between -950 ms to -100 ms was applied.
- Spherical interpolation of removed electrodes was applied to ensure comparability across datasets.

Critically, epoching and the interpolation of removed channels were conducted post-ICA to ensure that a continuous and valid data context was preserved, maximising AMICA performance. In total, 92% of channels were retained with an average of 4.6 (sd = 3.5) removed per dataset. Alternatively, 93% of components were retained with an average of 5.6 (sd = 2.5) removed per dataset.

### Event-related potential (ERP) analyses

ERP analyses were conducted using ERPLAB (v12.10 (Lopez-Calderon & Luck, 2014)) to acquire individual-level average waveforms and measurements for each time point, stimulated arm, and nociceptive modality, within each session:

- Prior to averaging, EEG data were band-pass filtered using a zero-phase, non-causal, Hamming-windowed FIR filter (transition width 0.3 Hz; passband edges [0.3 Hz 30 Hz]; cutoff frequencies at -6 dB [0.15 Hz 30.15 Hz]) (van den Broeke et al., 2020).
- ERP magnitudes and latencies were quantified by extracting peak amplitudes (Lopez-Calderon & Luck, 2014).

The vertex potential (VP) was quantified at Cz within a 50–500 ms post-stimulus window. Amplitude was taken as the maximum positive peak within this interval. Whilst the latency of neural responses evoked by different modalities are highly dependent on the conduction velocities of afferent fibres (Zhang et al., 2021), assessing VP peak latency provides a measure of when extralemniscal cortical activity was greatest.

### Event-related spectral perturbation (ERSP) analyses

The time-frequency decomposition of EEG data typically demonstrates a distinctive power distribution characterised by periodic narrowband oscillatory peaks combined with aperiodic 1/f-like activity (Donoghue et al., 2020; Gao et al., 2017). This distinction is critical for the analysis of ERSP as typically employed baselining methods such as division and log-transformation to decibel (dB) are dependent upon aperiodic activity (Gyurkovics et al., 2021) which can bias the estimation of evoked periodic oscillations (Samaha & Cohen, 2022). Therefore, as the present study aimed to investigate the reliability of oscillatory neural activity within fixed spectral frequency bands, Spectral Parameterisation Resolved in Time (SPRiNT) (version 1.0.3 (Wilson et al., 2022)) was used to estimate and remove the aperiodic component at each time. Furthermore, divisive baselining methods have been shown to bias the estimation of ERS over ERD, particularly when responses to acute nociceptive stimuli are assessed (Li Hu et al., 2014). Therefore, assuming an additive model of EEG (Gyurkovics et al., 2021), baseline correction was applied using subtraction (Zhang et al., 2021) following removal of the aperiodic slope:

- Short-time Fast Fourier Transforms (FFT) were used to compute time-frequency representations, providing uniform spectral resolution across frequencies. Spectra were estimated using 500 ms (250 sample) Hanning-window tapers with 80% overlap (Delorme & Makeig, 2004), yielding a native frequency resolution of 2 Hz. Spectra were zero-padded to interpolate estimates in 1 Hz steps, however, the true spectral resolution remained 2 Hz. The 500 ms window was selected to prioritise temporal tracking (100 ms step size) whilst maintaining adequate spectral resolution. Consequently, estimates at very low frequencies (< ∼4 Hz) were interpreted cautiously. Time-frequency estimates ranged from 1-30 Hz (1 Hz increments) and -700 to +1200 ms (100 ms steps).
- SPRiNT was applied to the averaged time-frequency decompositions for each participant at each time point, stimulated arm, and nociceptive modality, within sessions. Default spectral parameterisation parameters were used (peak detection threshold = 2.0 SD; peak widths = [0.5 12]; aperiodic mode = fixed, no minimum number of peaks; no maximum peak height) (Donoghue et al., 2020), alongside the majority of default SPRiNT parameters (Wilson et al., 2022). Notably, SPRiNT typically clusters oscillatory peaks over three-second windows, however, this is detrimental for temporal precision (Rawls & Sponheim, 2025) and therefore was not conducted.
- Baseline correction was applied using a -700 to -100 ms window and subtracting the mean power values at each frequency.
- ERSP magnitude was quantified as the mean of the lowest (ERD) or highest (ERS) 20% time-frequency points as applicable post-stimulus and within each relevant frequency band (Hu & Iannetti, 2019; Mouraux & Iannetti, 2008; Zhang et al., 2021).

Based upon previous research into acute pain perception, three regions of interest were defined for ERSP investigation:

- Theta-band (4 to 7 Hz) event-related synchronisation (θ-ERS) at the vertex electrode (Cz) (Iannetti et al., 2013; Michail et al., 2016; van den Broeke et al., 2017; Zhang et al., 2021).
- Alpha-band (8 to 13 Hz) desynchronisation (α-ERD) averaged across the respective contralateral central electrodes for the test-arm (FC3, FC5, C3, C5, CP3, CP5) and control-arm (FC4, FC6, C4, C6, CP4, CP6) separately (Hu et al., 2013; Ploner et al., 2017; Triccas et al., 2022; Zhang et al., 2021).
- Gamma-band (30 to 40 Hz) synchronisation (γ-ERS) at the vertex electrode (Cz) (Gousset et al., 2023; Linde et al., 2023; Schulz et al., 2015; Yue et al., 2025).

To preserve the flow of text time frequency figures are displayed within Supplementary Materials.

### Statistical Analyses

All statistical analyses were conducted using R (version 4.3.1) (R Core Team, 2023).

### Intraclass correlation coefficients

Test-retest reliability was assessed through intraclass correlation coefficients (ICC 3,1), using a two-way mixed-effects model with absolute agreement for single rater measurements (Koo & Li, 2016) implemented in the psych package (Revelle, 2018). ICCs were interpreted as follows:

- Values less than 0.5 were indicative of poor reliability.
- Values between 0.5 and 0.75 were indicative of moderate reliability.
- Values between 0.75 and 0.9 were indicative of good reliability. Values greater than 0.9 were indicative of excellent reliability.

### Bland Altman limits of agreement

Agreement between sessions was assessed using Bland-Altman analyses (Bland & Altman, 1986; Chhapola et al., 2015; De Vet et al., 2006; Koo & Li, 2016), quantifying bias and limits of agreement (mean differences ±1.96 SD) relative to the observed range of values across sessions. This was implemented in the blandr package (Datta, 2017).

### Linear mixed-effects modelling

To investigate HFS-induced changes in nociceptive-evoked EEG responses, statistical models were fit and interrogated for each EEG feature and stimulus modality separately. Furthermore, nociceptive-evoked EEG is not laterally homogenous, rendering direct comparisons between test and control-arm conditions unfeasible for assessing induced changes to magnitude or latency (Jin et al., 2018; Zhang et al., 2021). Therefore, further model separation was conducted for the test and control-arms of each feature and modality. Critically, this decision focused statistical investigation to ascertaining the presence and reliability of HFS-induced features, rather than potentially superfluous or confounding direct comparisons between the test and control-arms.

Data were analysed using linear mixed-effects models implemented in the lme4 package (Bates et al., 2015). Model assumptions were evaluated using the performance package (Lüdecke et al., 2021), including visual inspection of normality, linearity, collinearity, and homoscedasticity. Where violations were detected, optimal transformations were identified using the bestNormalize package (Peterson & Peterson, 2020). This resulted in log-plus-one transformations for pain rating data and ordered normalisation transformations for VP latency and ERSP data.

Data (*Y*) were subsequently modelled with fixed effects for time (*β1*), session (*β2*), age (*β3*), sex (*β4*), and their interactions (*β5*). Critically, interactions were separated for sex and age to avoid overfitting four-way comparisons. Random intercepts for each individual (*ui*) were included to account for a within-individuals repeated-measures design:

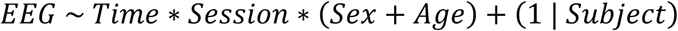

In addition, to investigate associations between nociceptive-evoked EEG and pain perception, EEG features were separately added as covariates to linear mixed-effect models as previously (Mugglestone et al., 2026) to assess their influence on reported pain intensity for each modality. Given the time-varying nature of ERP and ERSP data alongside pain ratings, it is critical to separate the between- and within-individuals variance so that valid inferences can be made (Hamaker & Muthén, 2020). Therefore, within each session, variance between-individuals was modelled as the baseline ERP/ERSP value, whilst post-HFS changes relative to these baselines were considered within-individual values. No other predictors were included to minimise collinearity and focus interpretations:

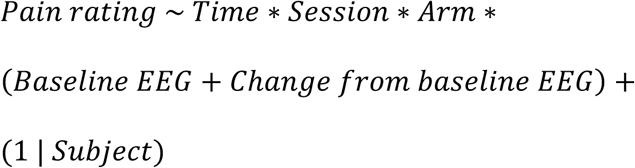

For each model, conditional and marginal estimates of variance and effect size were calculated using mixed model R^2^ and partial Eta^2^ respectively. Fixed effects were evaluated via three-way within-individuals Type III ANOVA with Satterthwaite’s method for adjusting degrees of freedom implemented in the lmerTest package (Kuznetsova et al., 2015). Pairwise comparisons were conducted using estimated marginal means with Dunnett’s adjustment for multiple comparisons, implemented in the emmeans package (Lenth et al., 2025). Standardised effect sizes are also reported. When applicable, Johnson–Neyman intervals were computed and visualised using the interactions package (Long & Long, 2021), applying a false discovery rate (FDR) at alpha = .05 to account for multiple comparisons (Esarey & Sumner, 2018). When necessary for visualisation, contrast-coding (-1; +1) was applied to the main predictor. However, this did not alter statistical significance or interpretation and was only implemented to facilitate visualisation.

## Results

### Demographic characteristics

A total of 35 participants were recruited to the study, however, following an interim analysis of *n* = 8 participants, discrepancies with the accuracy of pinprick EEG registration was noticed. Data collection was paused, and a problem was identified with the pinprick device. The device was returned to the manufacturer for repairs. Due to this equipment failure, 6 datasets that failed to retain valid pinprick trial markers were excluded. Subsequently, all raw data was visually assessed to identify for issues with pinprick-event generation. 1 additional individual was excluded due to extreme artifacts within the EEG recording resulting in a final sample of 27 participants (female = 19) comparable to similar previous work investigating nociceptive-evoked EEG within sensitised pain states (Gousset et al., 2023; van den Broeke et al., 2017; van den Broeke et al., 2015).

### Nociceptive-evoked EEG responses

To confirm the validity of EEG features according to well-established patterns of nociceptive-evoked responses, grand averaging across time points was conducted for each EEG feature, stimulus modality, and arm separately (Diers et al., 2020; Gousset et al., 2023; Hauck et al., 2015; L Hu et al., 2014; Hu & Iannetti, 2019; Hu et al., 2013; Jin et al., 2018; Liberati et al., 2016; Linde et al., 2023; Nickel et al., 2017; Perovic et al., 2025; Ploner et al., 2017; Schulz et al., 2015; Somervail et al., 2025; Somervail et al., 2021; Tiemann et al., 2026; E. van den Broeke et al., 2021; van den Broeke et al., 2020; van den Broeke et al., 2017; van den Broeke et al., 2019; Van den Broeke et al., 2016; van den Broeke et al., 2015; Wang et al., 2023; Yue et al., 2025; Zhang et al., 2021; Zidda et al., 2024). These responses are displayed within Figure 1 and Figure 2, alongside visualisations of pain intensity rating data as reported previously (Mugglestone et al., 2026). Only MPS demonstrated significant enhancement following HFS (Figure 1a).

**Figure 1.**
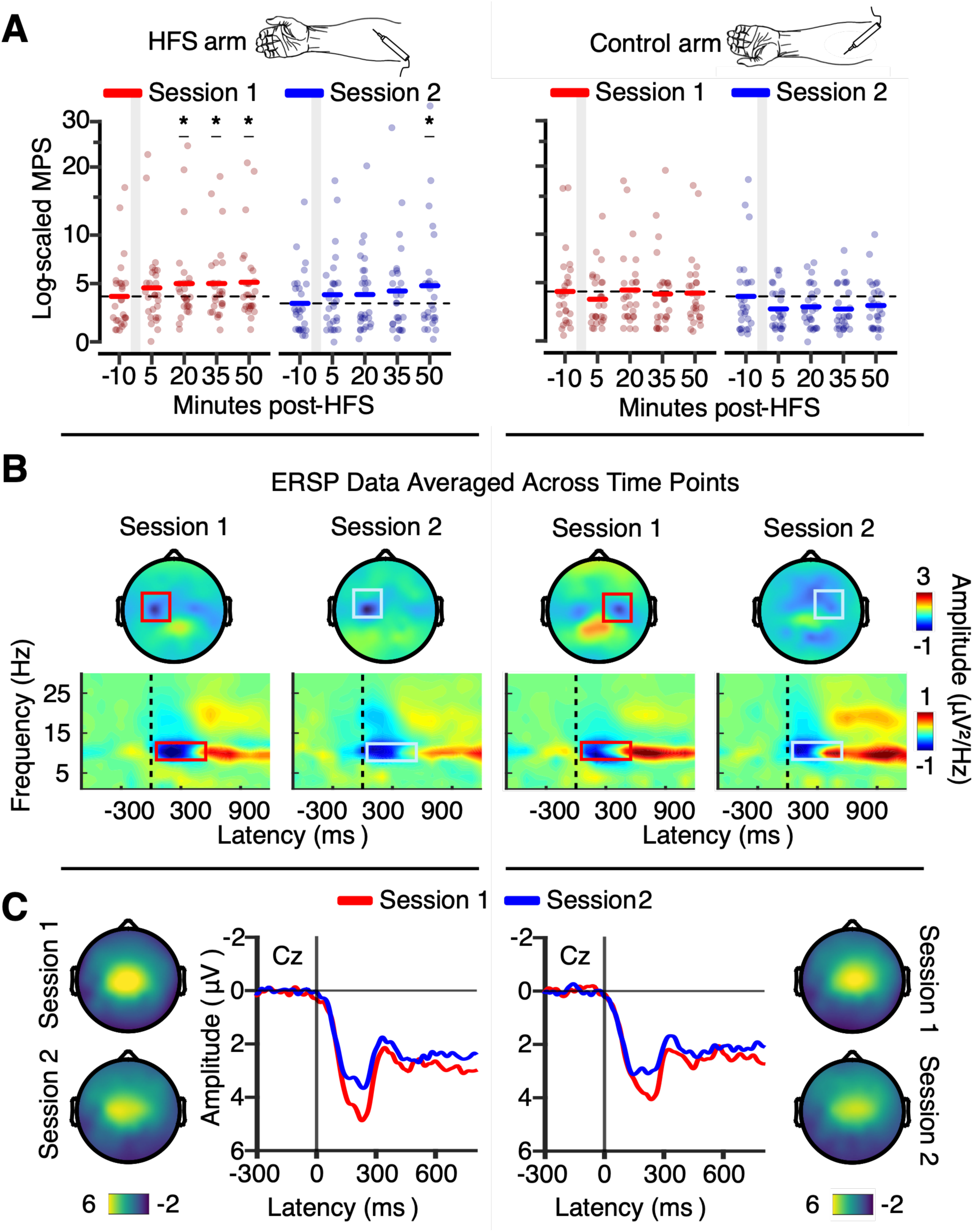
Mechanical pinprick responses grand averaged across time points. a) Changes in reported pain intensity over time corresponding to pinprick stimuli for HFS (left) and control (right) arms. Dot plots represent individual reported pain intensities displayed as raw data points alongside group mean bars for each time point, session, and arm separately. b) ERSP data and α-ERD topographies grand-averaged over all time points for each session and arm separately for lower frequency responses at contralateral central electrodes. c) ERP data and topographies grand averaged over all time points for each session arm separately. HFS = High-frequency stimulation. α-ERD = Alpha-band desynchronisation. VP = Vertex potential. MPS = Mechanical pinprick sensitivity. ERSP = Event-related spectral perturbation. ERP = Event-related potential.

**Figure 2.**
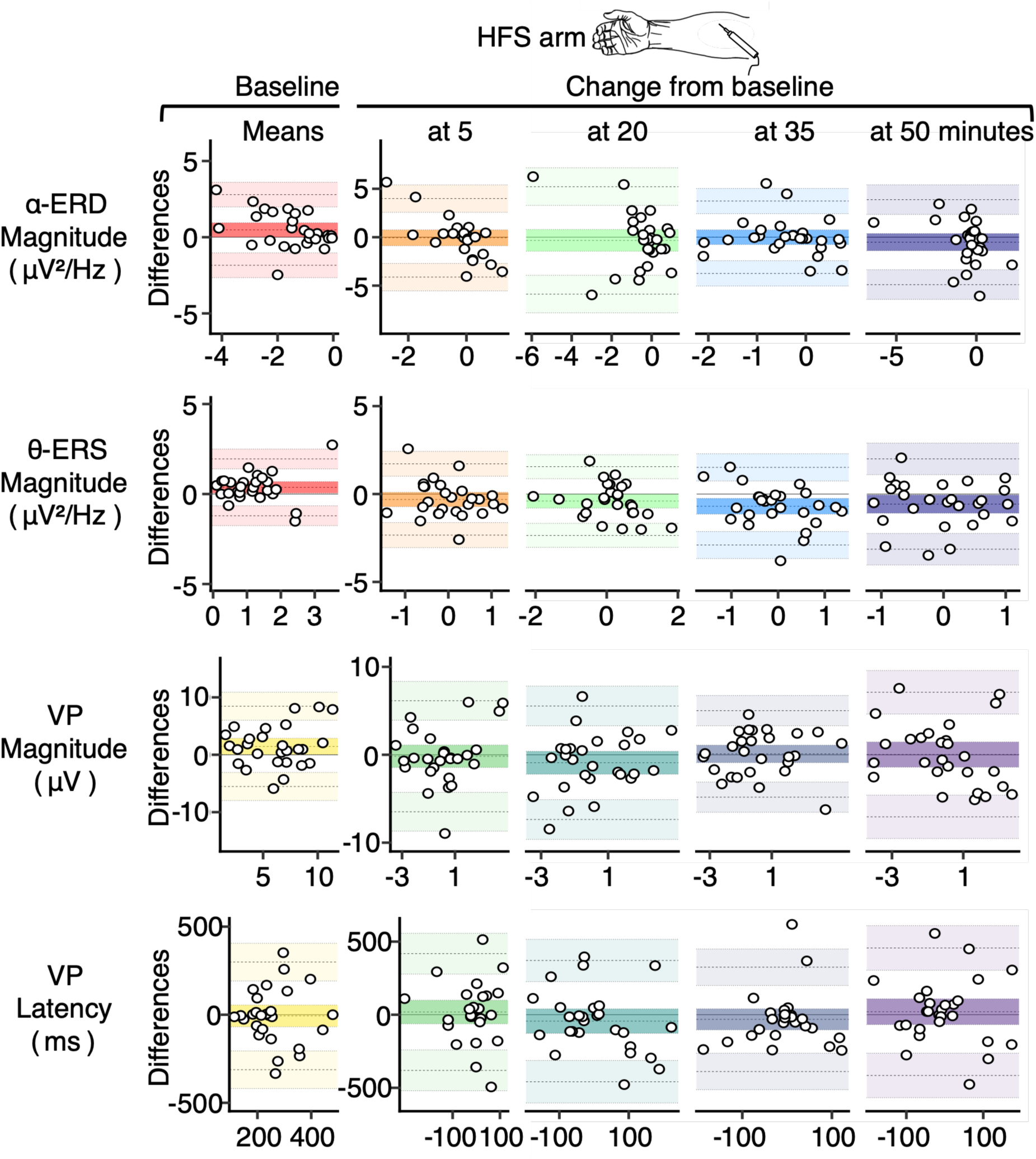
Electrical stimulation responses grand averaged across time points. a) Changes in reported pain intensity over time corresponding to electrical stimuli for HFS (left) and control (right) arms. Dot plots represent individual reported pain intensities displayed as raw data points alongside group mean bars for each time point, session, and arm separately. b) ERSP data and α-ERD topographies grand-averaged over all time points for each session and arm separately for lower frequency responses at contralateral central electrodes. c) ERP data and topographies grand averaged over all time points for each session arm separately. HFS = High-frequency stimulation. α-ERD = Alpha-band desynchronisation. VP = Vertex potential. EPP = Electrical pain perception. ERSP = Event-related spectral perturbation. ERP = Event-related potential

Consistent with previous research, robust ERSP α-ERD responses were evident at contralateral central electrodes across stimulus modalities (Figure 1b) (Hauck et al., 2015; Hu & Iannetti, 2019; Hu et al., 2013; Jin et al., 2018; Liberati et al., 2016; Nickel et al., 2017; Ploner et al., 2017; Tiemann et al., 2026; van den Broeke et al., 2020; van den Broeke et al., 2017; Wang et al., 2023; Yue et al., 2025; Zhang et al., 2021; Zidda et al., 2024). However, when assessed at the vertex electrode, whilst θ-ERS was present across modalities, γ-ERS was only evident at a group level for electrical stimuli. This finding is unsurprising as γ-ERS is predominantly exhibited within thermonociceptive modalities (Hu & Iannetti, 2019) and recent investigations into pinprick-evoked γ-ERS have failed to identify significant gamma fluctuations post-HFS (Gousset et al., 2023; Linde et al., 2023; Schulz et al., 2015; Yue et al., 2025). Therefore, whilst γ-ERS ERSP data are visualised within Supplementary Materials for both pinprick and electrical stimuli, statistical investigations were only conducted for electrical-evoked γ-ERS. Furthermore, whilst robust electrical-evoked VP waveforms (4c) were evident (Perovic et al., 2025; Somervail et al., 2025; Somervail et al., 2021; Tiemann et al., 2026; Yue et al., 2025), pinprick-evoked VP were comparatively weaker, which may have been due to the use of a handheld mechanical pinprick device resulting in variations to force profiles and stimuli durations (Triccas et al., 2022; van den Broeke et al., 2020; van den Broeke et al., 2017). These limitations are discussed further below.

### Test-retest reliability of pinprick-evoked responses

To investigate the test-retest reliability of pinprick-evoked EEG features at baseline and across the HFS development time course, ICCs were calculated alongside Bland-Altman mean biases and LOA, which are displayed in Figure 3 and Figure 4. This test-retest reliability data is also reported within Supplementary Table 1 alongside descriptive statistics concerning means, standard deviations, and ranges, of baseline and change from baseline ERSP and ERP data.

**Figure 3.**
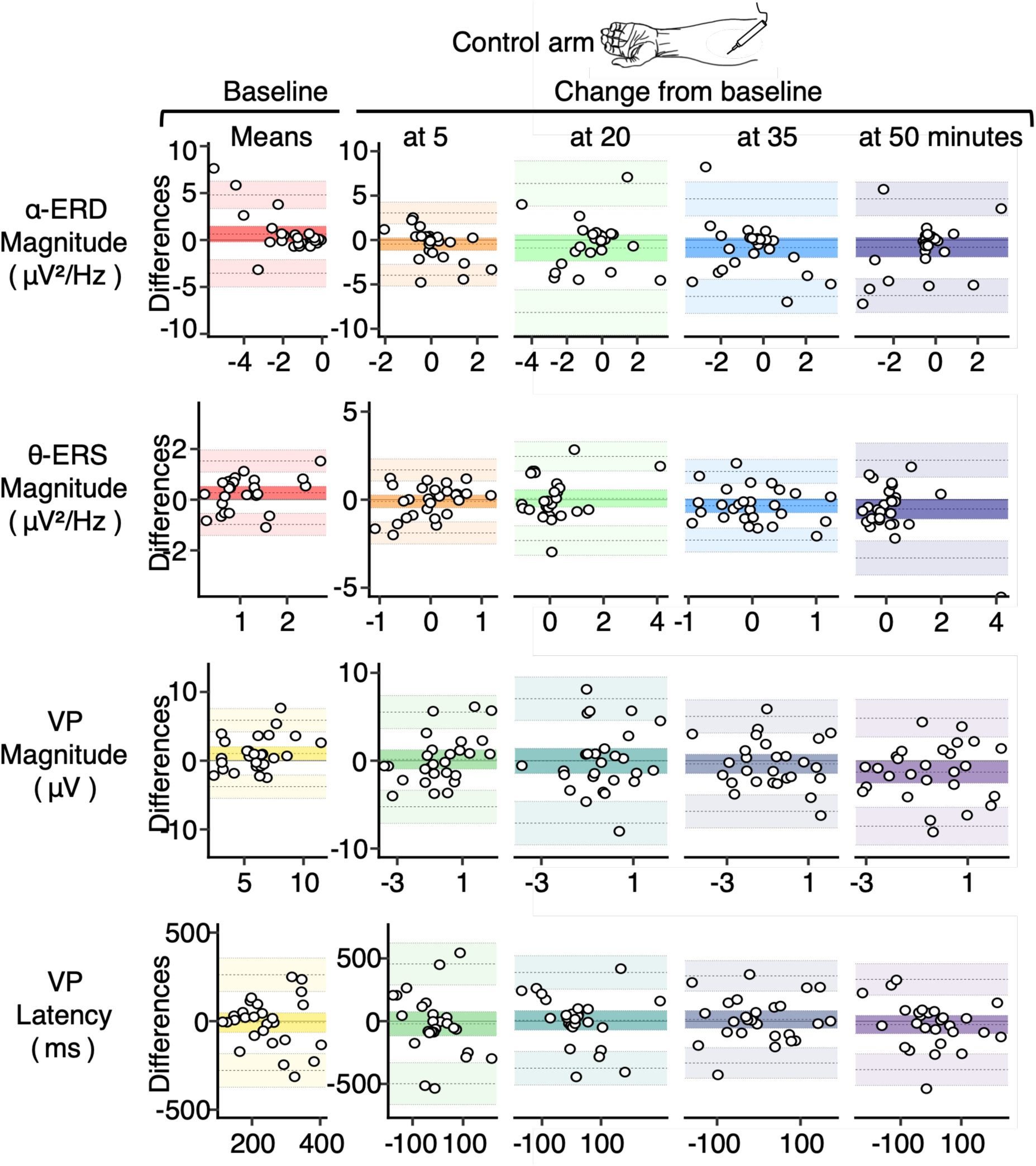
Bland-Altman assessments of agreement for HFS arm pinprick-evoked EEG. For each measure, Bland-Altman plots represent the mean bias (middle line), lower (bottom line) and upper (top line) limits of agreement, alongside 95% confidence intervals. Differences between measurements (y-axis) and the mean of paired measurements (x-axis) are displayed for each individual. α-ERD = alpha-band event-related desynchronisation; θ-ERS = theta-band event-related synchronisation; VP = vertex potential.

**Figure 4.**
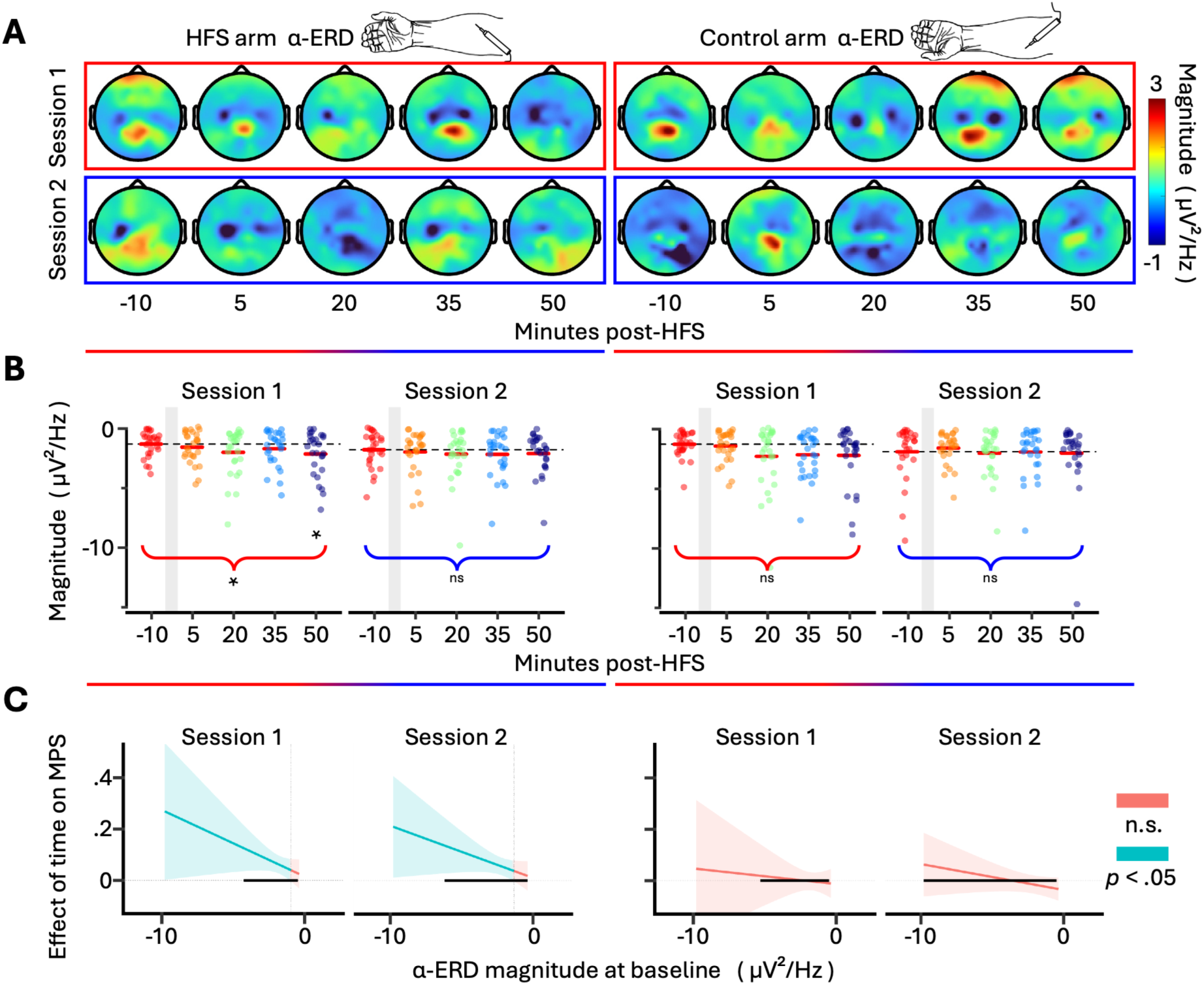
Bland-Altman assessments of agreement for control arm pinprick-evoked EEG. For each measure, Bland-Altman plots represent the mean bias (middle line), lower (bottom line) and upper (top line) limits of agreement, alongside 95% confidence intervals. Differences between measurements (y-axis) and the mean of paired measurements (x-axis) are displayed for each individual. α-ERD = alpha-band event-related desynchronisation; θ-ERS = theta-band event-related synchronisation; VP = vertex potential.

At baseline, ICCs were generally higher for ERSP measures than ERP measures. θ-ERS demonstrated moderate reliability across arms (HFS: ICC = .59; control: ICC = .57) whilst α-ERD demonstrated moderate reliability for the HFS arm but poor reliability for the control arm (HFS: ICC = .58; control: ICC = .28). In contrast, VP magnitude demonstrated poor-to-moderate reliability across arms (HFS: ICC = .41; control: ICC = .50), whilst VP latency was poor (HFS: ICC = .16; control: ICC = .29). Bland-Altman analyses indicated relatively small mean biases across all measures relative to the observed ranges across sessions, although LOA were substantially wider for ERP responses in comparison to ERSP responses.

Following HFS, change from baseline ICCs ranged from poor to extremely poor, with many values approaching zero indicating that within-subject variability exceeded between-subject variability, likely due to a lack of robust directional changes post-HFS resulting in near zero mean biases. This was accompanied by considerably wider LOA when change from baseline measures were assessed. α-ERD change from baseline ICCs in ascending time order were 0, 0, 0, .26 for the HFS arm and .03, 0, .04, 0 for the control arm. θ-ERS change from baseline ICCs were .22, .41, .25, .06 for the HFS arm and .19, .46, .15, .33 for the control arm. VP magnitude change from baseline ICCs were .26, .23, .55, .25 for the HFS arm and .25, 0, .23, .01 for the control arm. VP latency change from baseline ICCs were 0, .1, 0, 0 for the HFS arm and 0, 0, .1, .15 for the control arm.

Overall, pinprick-evoked EEG responses demonstrated the strongest test-retest reliability at baseline for ERSP measures. However, no feature demonstrated good reliability (Koo & Li, 2016). Furthermore, reliability deteriorated substantially when assessing HFS-induced changes over time although mean biases remained low, indicating that across repeated test sessions, intra-individual variations from baseline were greater than inter-individual changes.

### Post-HFS changes in pinprick-evoked EEG responses

To investigate HFS-induced changes in pinprick-evoked EEG responses, mixed-effects modelling was used to assess change over time, between sessions, and the influence of age and sex for each EEG feature and arm separately. Across pinprick-evoked EEG responses, α-ERD and VP magnitude demonstrated the strongest evidence for change over time and across sessions, whilst no other features demonstrated significant or consistent directional effects.

For the α-ERD (Supplementary Table 2), a significant negative effect of time was evident for the HFS-arm (Figure 5a-b), with decreasing ERSP values indicating stronger α-ERD relative to baseline (*F* (4, 243) = 4.7, *p* = .001). Although this main effect of time was consistent across time points, it was driven by a significant pairwise comparison to baseline at 50 minutes post-HFS within session 1 (*Std. effect* = -.82 [-1.5, -.2], *p* = .02). However, a significant session effect was also evident indicating lower α-ERD within session 2 (*F* (1, 243) = 8.17, *p* = .005), although the interaction between time and session was not significant (*F* (4, 243) = .23, *p* = .92). Alternatively, no significant main effects were present for the α-ERD control arm, although the interaction between session and age (*F* (1, 243) = 5.73, *p* = .02) indicated that individual fluctuations in α-ERD between repeated test sessions were greater for older participants (> 60). In contrast, θ-ERS (Supplementary Table 3) demonstrated no main effects for either arm (Figure 6a-b), although a significant time-sex interaction was present for the HFS arm (*F* (4, 243) = 2.63, *p* = .04), as male participants (*F* (4, 92) = 1.63, *p* = .17) demonstrated greater fluctuations in θ-ERS compared to female participants (*F* (4, 92) = .99, *p* = .42), driving a significant pairwise comparison at 50 minutes post-HFS during session 1 (*Std. effect* = -.81 [-1.44, - .18], *p* = .04).

**Figure 5.**
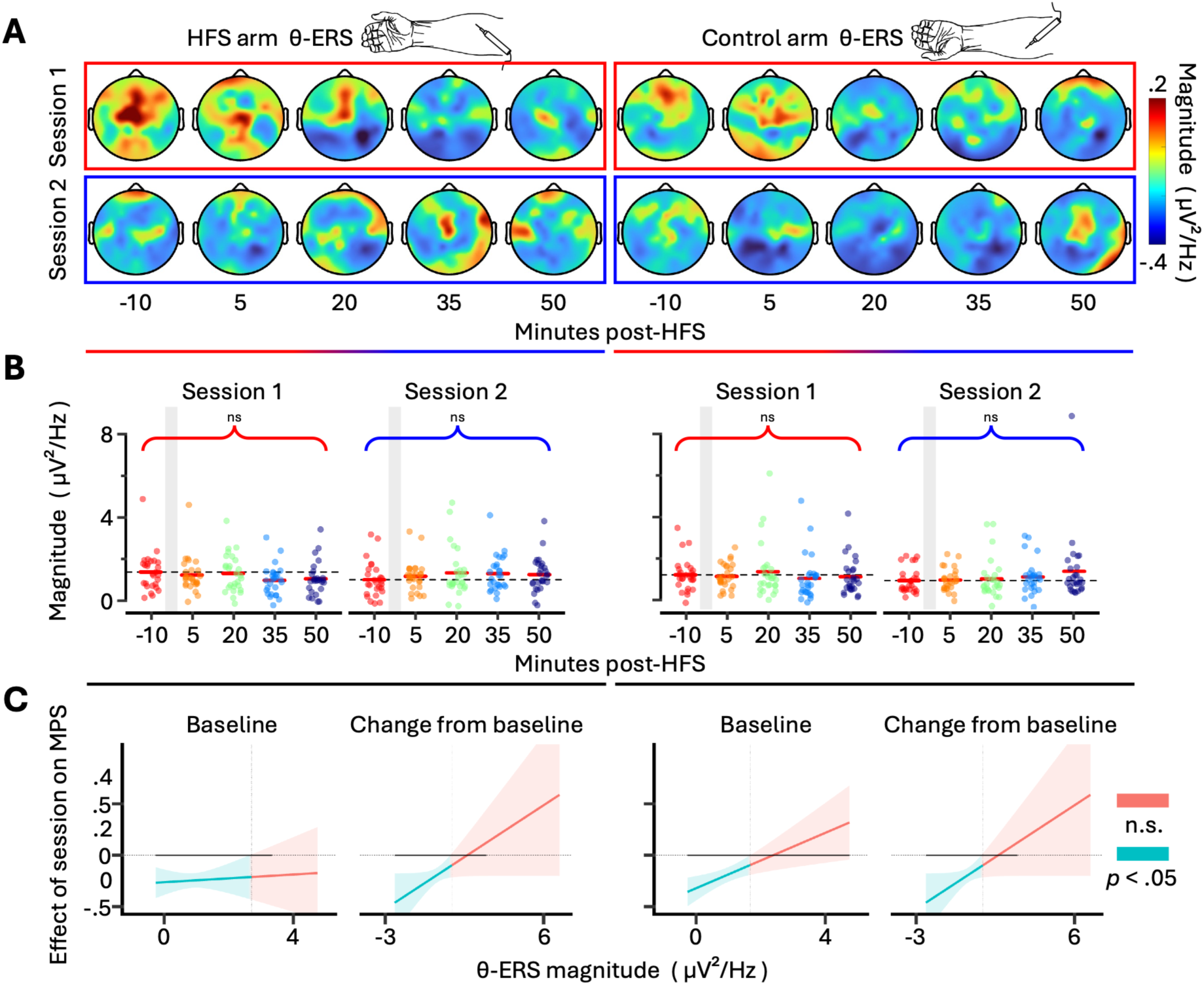
Pinprick-evoked α-ERD results. ERSP topography plots (top) represent the grand-averaged α-ERD across all participants for the HFS (left) and control (right) arms. Scalp topographies were averaged between 0ms to 500ms to capture the main α-ERD response between 8 Hz to 13 Hz. Dot plots (middle) represent individual α-ERD magnitudes displayed as raw data points alongside group means (red bars), for each time point, session, and arm separately. Statistics were derived from linear mixed-effects modelling and reflect change from baseline for each arm and session respectively. Regression plots (bottom) represent the relationships between baseline α-ERD magnitude and MPS change over time (bottom) for each session and arm separately, with regression lines derived from linear mixed-effects modelling demonstrating the estimated slope of change over time. ^ns^ indicates no statistical significance (p > .05); * indicates significance at alpha = 0.05; HFS = High-frequency stimulation; MPS = Mechanical pinprick sensitivity; α-ERD = alpha-band event-related desynchronisation.

**Figure 6.**
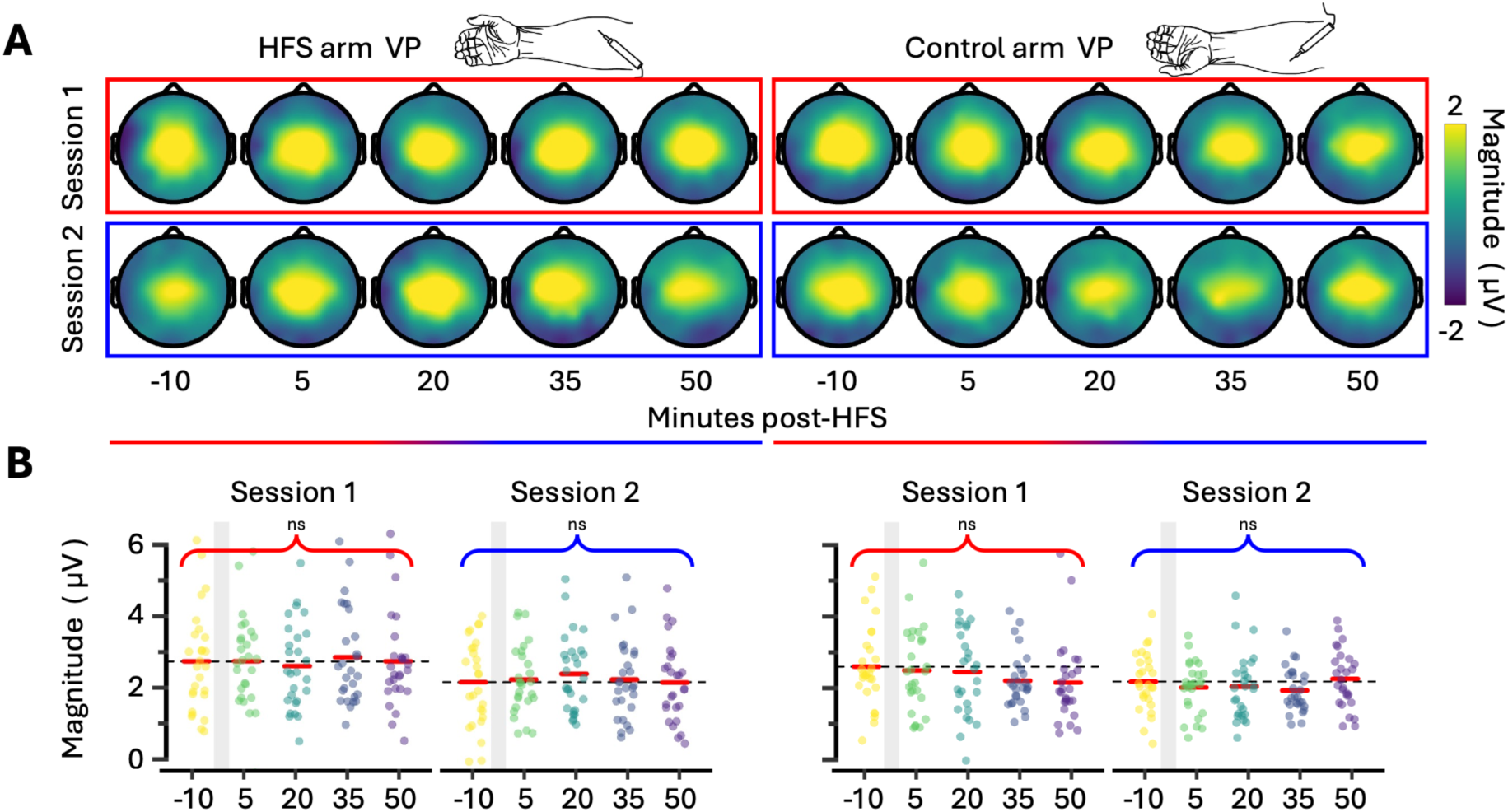
Pinprick-evoked θ-ERS results. ERSP topography plots (top) represent the grand-averaged θ-ERS across all participants for the HFS (left) and control (right) arms. Scalp topographies were averaged between 0ms to 500ms to capture the main θ-ERS response between 4 Hz to 7 Hz.Dot plots (middle) represent individual θ-ERS magnitudes displayed as raw data points alongside group means (red bars), for each time point, session, and arm separately. Statistics were derived from linear mixed-effects modelling and reflect change from baseline for each arm and session respectively.Regression plots (bottom) represent the relationships between MPS change between sessions and θ-ERS magnitude both at baseline and as change from baseline for each arm separately, with regression lines derived from linear mixed-effects modelling demonstrating the estimated slope of change between sessions. ^ns^ indicates no statistical significance (p > .05); * indicates significance at alpha = 0.05; HFS = High-frequency stimulation; MPS = Mechanical pinprick sensitivity; θ-ERS = theta-band event-related synchronisation

Alternatively, VP magnitude (Supplementary Table 4) exhibited a significant negative session effect for both the HFS (*F* (1, 243) = 25.29, *p* < .001) and control (*F* (1, 243) = 14.96, *p* < .001) arms (Figure 7a-b and Figure 8), demonstrating that systematic differences between test sessions were evident at baseline and persisted throughout the HFS development time course. For the HFS arm this was driven by multiple significant between-sessions pairwise comparisons at baseline (*Std. effect* = -.79 [-1.43, -.16], *p* = .014), and at 5 (*Std. effect* = -.85, [-1.49, -.22], *p* = .009) and 35 (*Std. effect* = -.84 [-1.47, - .21], *p* = .009) minutes post-HFS. For the control arm, this was driven pairwise comparisons were significant at baseline (*Std. effect* = -.68 [-1.32, -.05], *p* = .03), and at 5 (*Std. effect* = -.75 [-1.38, -.11], *p* = .02) and 20 (*Std. effect* = -.69 [-1.32, -.05], *p* = .03) minutes post-HFS. Interestingly, the control arm session effect was greater for younger participants (*F* (1, 243) = 6.03, *p* = .015) with the threshold of significance at < 39 years. However, VP latency (Supplementary Table 5) demonstrated no significant main effects for either arm, although a significant session-age interaction (*F* (1, 243) = 8.68, *p* < .001) demonstrated greater fluctuations in HFS arm VP latency for older participants with the threshold of significance at > 42 years.

**Figure 7.**
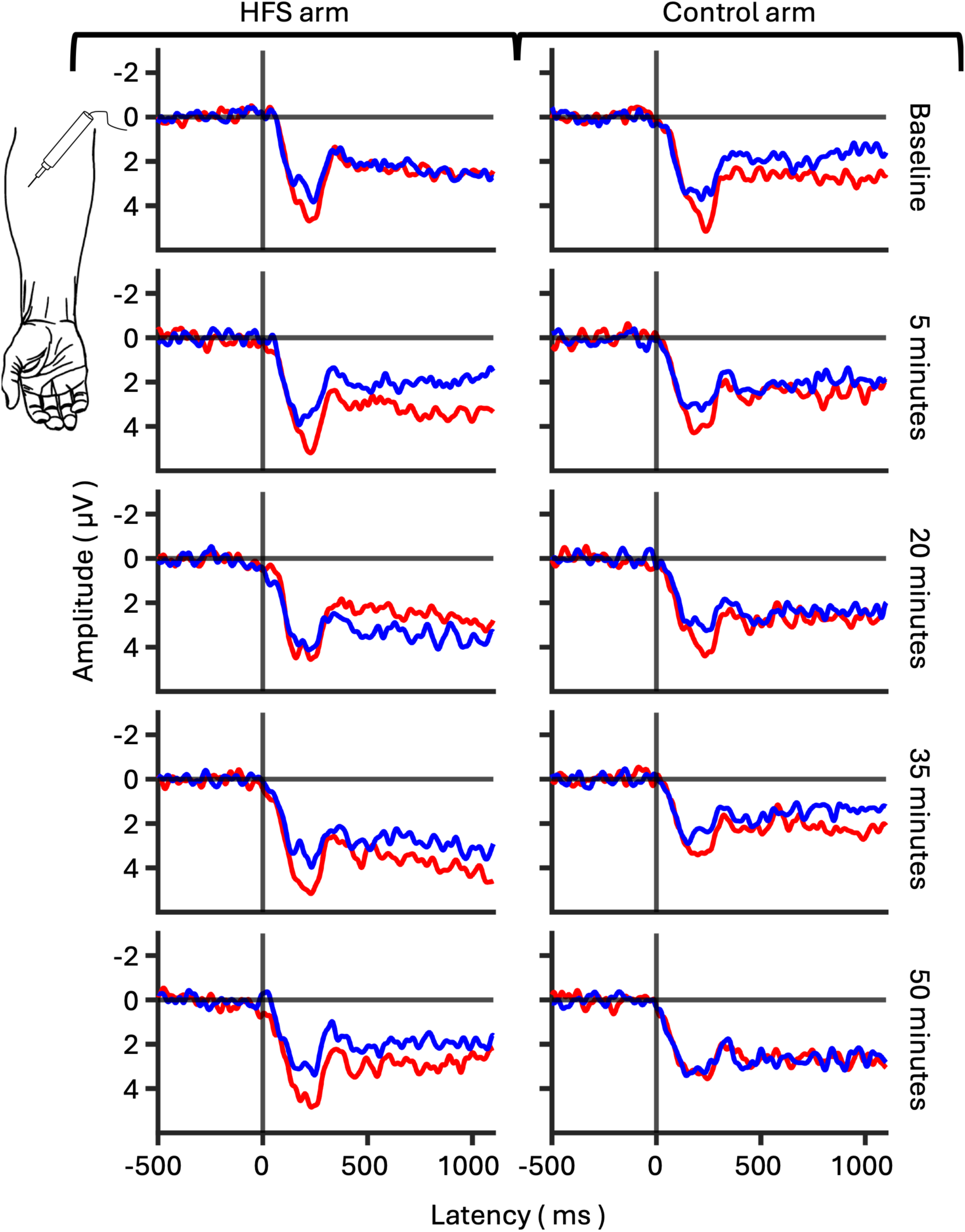
Pinprick-evoked VP results. ERP plots (top) represent the grand-averaged VP across all participants for the HFS (left) and control (right) arms. Statistics were derived from linear mixed-effects modelling and reflect change between sessions VP magnitude for each arm and time point respectively. Dot plots (middle) represent individual VP magnitudes displayed as raw data points alongside group means (red bars), for each time point, session, and arm separately. Statistics were derived from linear mixed-effects modelling and reflect change from baseline VP magnitude for each arm and session respectively. ^ns^ indicates no statistical significance (p > .05); * indicates significance at alpha = 0.05; HFS = High-frequency stimulation; MPS = Mechanical pinprick sensitivity; VP = Vertex potential; ERP = Event related potential.

**Figure 8.**
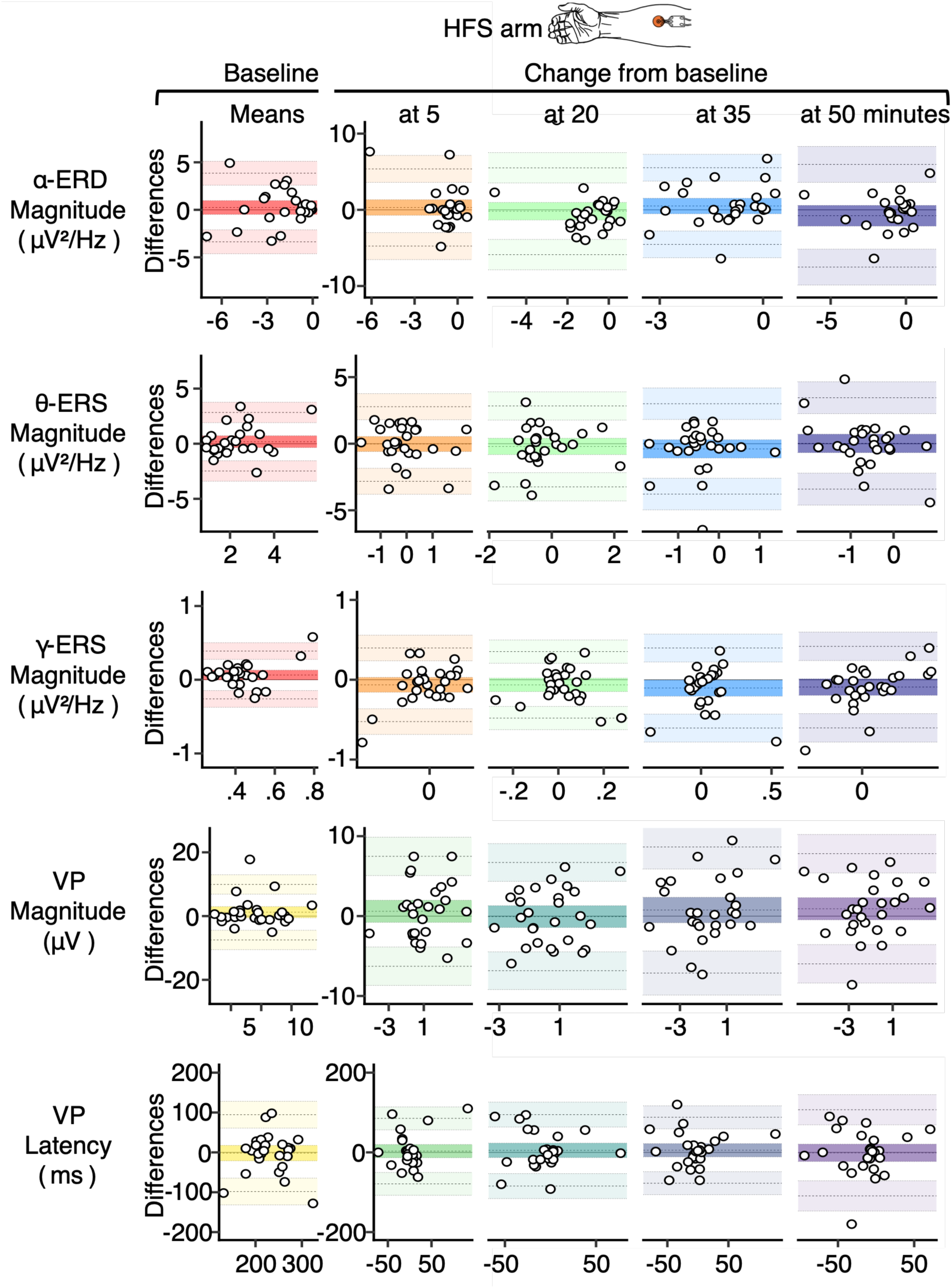
Pinprick-evoked VP ERP plots. ERP plots represent the grand-averaged VP at the vertex electrode (Cz) across all participants for both the HFS (left) and control (right) arms and sessions 1 (red) and 2 (blue). VP = vertex potential. HFS = High-frequency stimulation.

Overall, HFS-induced changes to pinprick-evoked EEG responses were only apparent for α-ERD, which were most substantial at 50 minutes post-HFS during session one. Alternatively, VP magnitude also displayed a significant decrease in amplitude between repeated test sessions which was consistent across arms. However, θ-ERS, VP magnitude, and VP latency demonstrated limited sensitivity to HFS, with varied fluctuations over time and between sessions influenced by inter-individual differences in age and sex.

### Relationships between pinprick-evoked EEG responses and MPS

To investigate the relationship between pinprick-evoked EEG responses and reported pain intensity, mixed-effects modelling was used to characterise changes in MPS over time, between sessions, and across arms as previously (Mugglestone et al., 2026), with the addition of baseline and change from baseline EEG data included as covariates to capture between- and within-individuals variance respectively (Hamaker & Muthén, 2020). Johnson-Neyman intervals are shown in Figure 5c, Figure 6c, and Figure 7c to visualise interactions as applicable.

Across pinprick-evoked EEG responses, change from baseline VP magnitude was the only measure directly associated with MPS. However, HFS-induced changes in MPS over time were related to the magnitude of pinprick-evoked α-ERD at baseline. For all models, significant effects for session (all *F* (1, 513) > 54.23, all *p* < .001), time (all *F* (1, 513) > 7.46, all *p* < .007), and arm (all *F* (1, 513) > 36.48, all *p* < .001) were present, alongside the significant interaction between time and arm (all *F* (1, 513) > 13.99, all *p* < .003), demonstrating the induction of SHA post-HFS as previously reported (Mugglestone et al., 2026).

For α-ERD (), neither baseline magnitude (*F* (1, 535) = .02, *p* = .88) nor change from baseline (*F* (1, 519) = .88, *p* = .35) were directly associated with MPS. However, a significant interaction between time and baseline α-ERD was present (*F* (1, 513) = 4.89, *p* = .027), suggesting that changes in MPS over time were related to inter-individual differences in α-ERD. Subsequently, Johnson-Neyman visualisations revealed that this effect was only present for the HFS arm (Figure 5c) as no significant post-HFS changes were evident for the control arm as previously reported (Mugglestone et al., 2026), demonstrating that the significance of post-HFS changes in MPS was related to greater α-ERD (lower magnitudes) at baseline.

Additionally, whilst pinprick-evoked θ-ERS also demonstrated no direct significant relationships with MPS, significant interactions with session were present for both baseline (*F* (1, 514) = 4.05, *p* = .05) and change from baseline (*F* (1, 514) = 5.4, *p* = .02) magnitudes. Accordingly, Johnson-Neyman visualisations demonstrated that greater θ-ERS was associated with less significant changes in MPS across repeated test sessions (Figure 6c), an effect that was consistent across arms.

In contrast, change from baseline VP magnitude was directly related to MPS (*F* (1, 519) = 4.25, *p* = .04), suggesting that fluctuations in reported pain intensity were reflected by changes in VP magnitude relative to individual-specific baselines (Figure 7c). However, no significant influence of baseline VP magnitude was present (*F* (1, 533) = 2.63, *p* = .11).

However, neither baseline VP latency (*F* (1, 516) = 1.67, *p* = .2) nor change from baseline VP latency (*F* (1, 522) = 2.33, *p* = .13) exhibited any significant influence over MPS.

Overall, α-ERD magnitude at baseline demonstrated a significant influence over subsequent HFS-induced changes to MPS, an effect that was specific to the HFS arm and demonstrates that inter-individual differences in pinprick-evoked α-ERD prior to HFS conditioning may influence the time course of SHA development. However, whilst θ-ERS was associated with changes to MPS across repeated test sessions, only change from baseline VP magnitude was directly related to MPS.

### Test-retest reliability of electrical-evoked responses

To investigate the test-retest reliability of electrical-evoked EEG features ICCs were calculated alongside the Bland-Altman mean biases and LOA presented in Figure 9 and Figure 10. These test-retest reliability measures are reported within Supplementary Table 6, alongside descriptive statistics concerning means, standard deviations, and ranges for baseline and change from baseline values.

**Figure 9.**
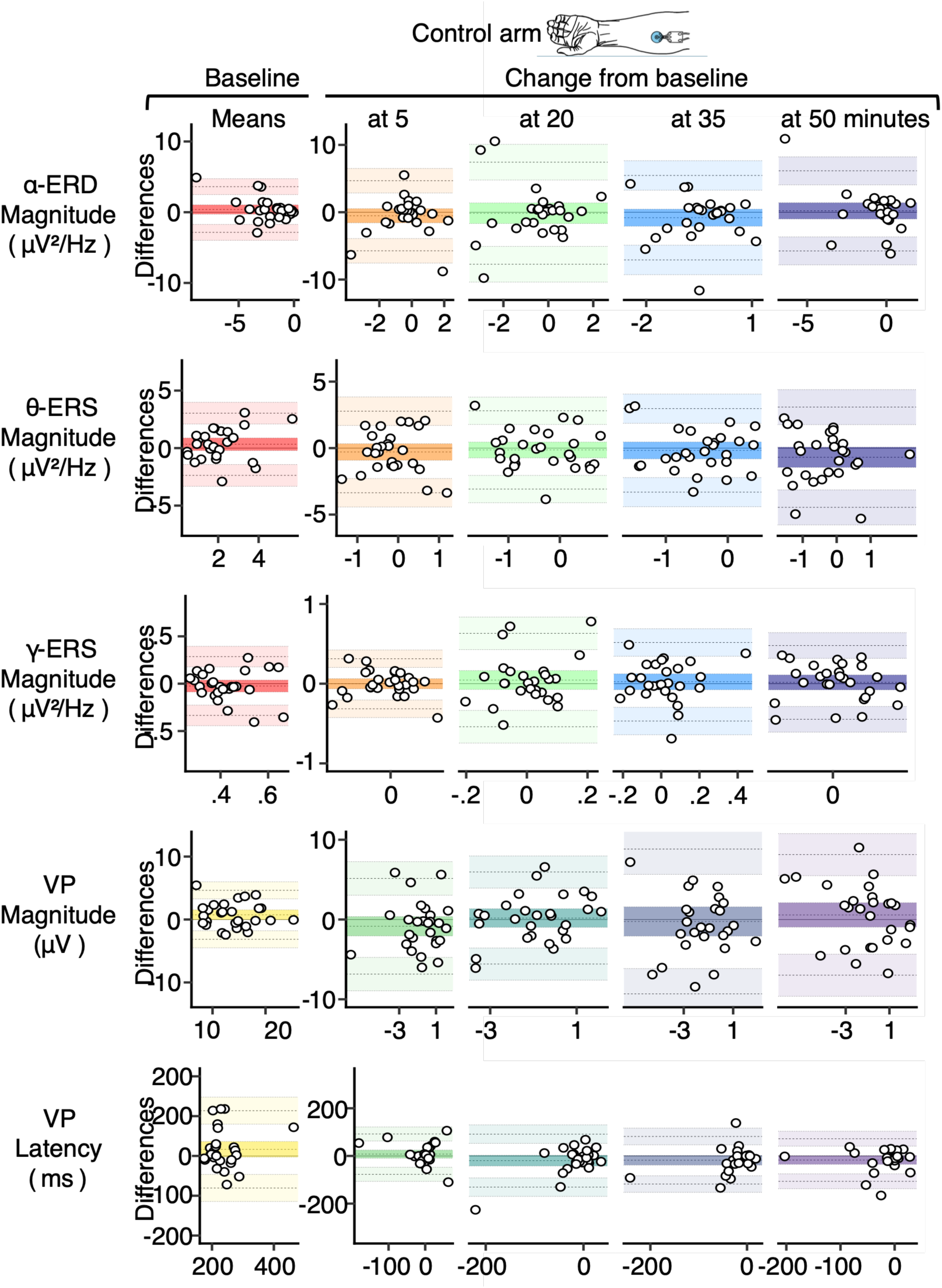
Bland-Altman assessments of agreement for HFS arm electrical-evoked EEG. For each measure, Bland-Altman plots represent the mean bias (middle line), lower (bottom line) and upper (top line) limits of agreement, alongside 95% confidence intervals. Differences between measurements (y-axis) and the mean of paired measurements (x-axis) are displayed for each individual. α-ERD = alpha-band event-related desynchronisation. θ-ERS = theta-band event-related synchronisation. γ-ERS = Gamma-band event-related synchronisation. VP = vertex potential. HFS = High-frequency stimulation.

**Figure 10.**
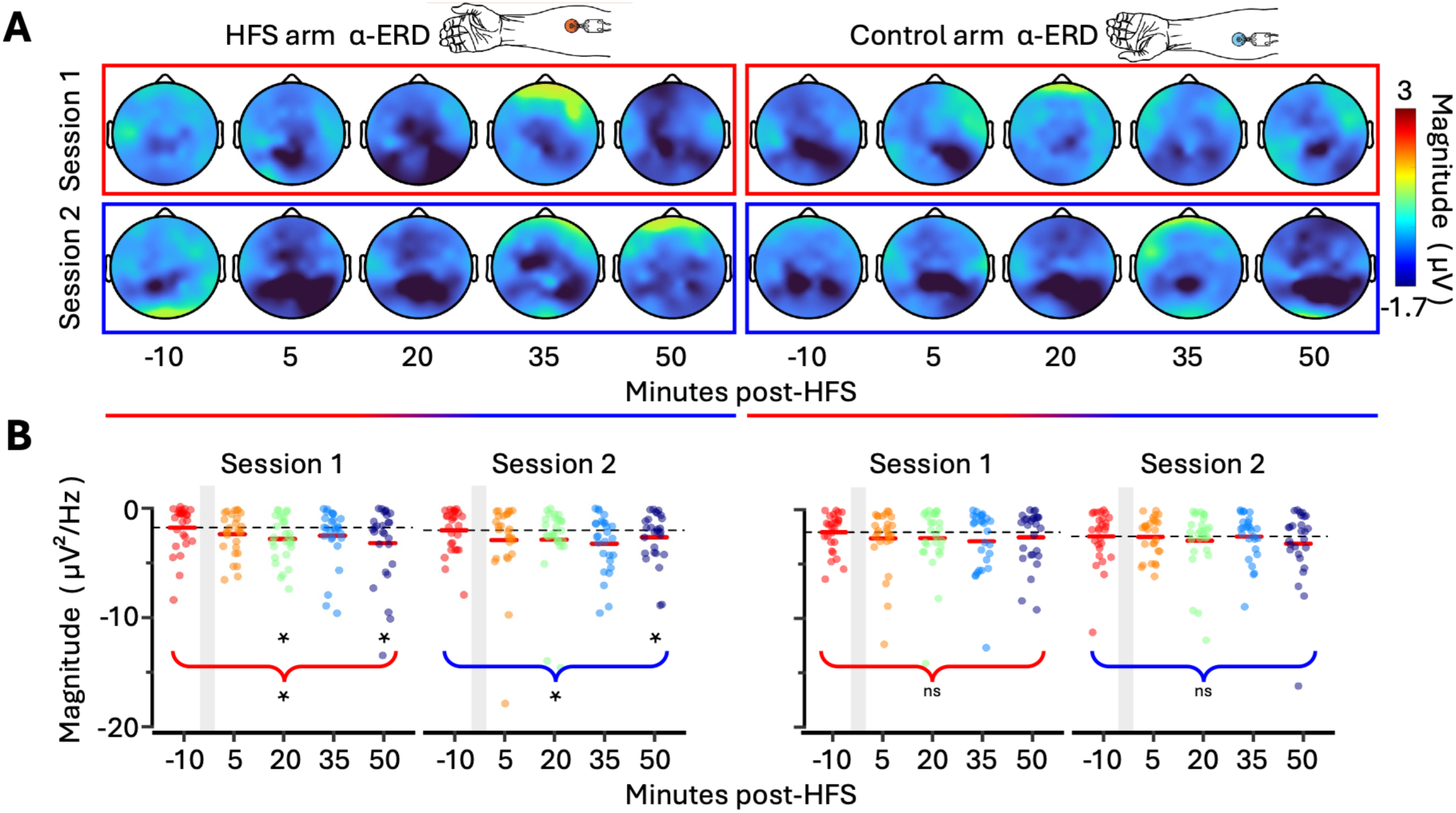
Bland-Altman assessments of agreement for control arm electrical-evoked EEG. For each measure, Bland-Altman plots represent the mean bias (middle line), lower (bottom line) and upper (top line) limits of agreement, alongside 95% confidence intervals. Differences between measurements (y-axis) and the mean of paired measurements (x-axis) are displayed for each individual. α-ERD = alpha-band event-related desynchronisation. θ-ERS = theta-band event-related synchronisation. γ-ERS = Gamma-band event-related synchronisation. VP = vertex potential. HFS = High-frequency stimulation.

At baseline, electrical-evoked ICCs were generally higher than their pinprick-evoked counterparts reported above. However, whilst α-ERD demonstrated moderate reliability (HFS: ICC = .51; control: ICC = .7), θ-ERS demonstrated poor reliability across both arms (HFS: ICC = .45; control: ICC = .48). Alternatively, VP magnitude demonstrated moderate reliability (HFS: ICC = .53; control: ICC = .53), whilst VP latency was moderate-to-poor (HFS: ICC = .51; control: ICC = .4). Electrical-evoked Bland-Altman LOA were also relatively narrower than pinprick-evoked responses, alongside smaller mean biases relative to observed ranges across repeated test sessions.

However, following HFS, electrical-evoked change from baseline ICCs also decreased substantially, with many values approaching zero, accompanied by wider LOA. α-ERD change from baseline ICCs in ascending time order were 0, 0, 0, 0 for the HFS arm and 0, 0, 0, .06 for the control arm. θ-ERS change from baseline ICCs were .26, .09, 0, 0 for the HFS arm and 0, 0, 0, 0 for the control arm. VP magnitude change from baseline ICCs were .32, .13, .23, .39 for the HFS arm and .36, .16, 0, .39 for the control arm. VP latency change from baseline ICCs were .46, .06, .47, .16 for the HFS arm and .52, .26, .2, .2 for the control arm.

Overall, whilst the test-retest reliability of electrical-evoked EEG responses were generally higher than pinprick-evoked EEG responses, none reached good reliability (Koo & Li, 2016), and reliability deteriorated substantially when change from baseline was assessed.

### Post-HFS changes in electrical-evoked EEG responses

To investigate changes in electrical-evoked EEG responses over time, mixed-effects modelling was used. Across electrical-evoked EEG responses, all features demonstrated significant changes, either over time or between sessions. These dynamics were often substantially influenced by individual differences across demographical characteristics.

For α-ERD (Supplementary Table 7), a significant negative time effect (Figure 11) was present for the HFS arm (*F* (4, 243) = 8.71, *p* < .001), driven by significant pairwise comparisons at 5 (*Std. effect* = -.99 [-1.63, -.34], *p* = .011), 20 (*Std. effect* = -1.5 [-2.15, - .84], *p* < .001), and 50 (*Std. effect* = -1.23 [-1.88, -.58], *p* = .001) minutes post-HFS during session 1, and at 5 (*Std. effect* = -.85 [-1.49, -.02], *p* = .037) and 35 (*Std. effect* = -1 [-1.64, -.35], *p* = .011) minutes post-HFS during session 2. Furthermore, although no main effect of session was present, a significant pairwise comparison at 35 minutes post-HFS demonstrated a decrease in electrical-evoked α-ERD (*Std. effect* = -.73 [-1.37, -.09], *p* = .025). Additionally, whilst no significant main effects were present for the control arm, interactions between session and age (*F* (1, 243) = 5.78, *p* = .02), session and sex (*F* (1, 243) = 4.46, *p* = .04), and time and sex (*F* (4, 243) = 2.95, *p* = .02) were significant, suggesting that electrical-evoked α-ERD fluctuated across demographics.

**Figure 11.**
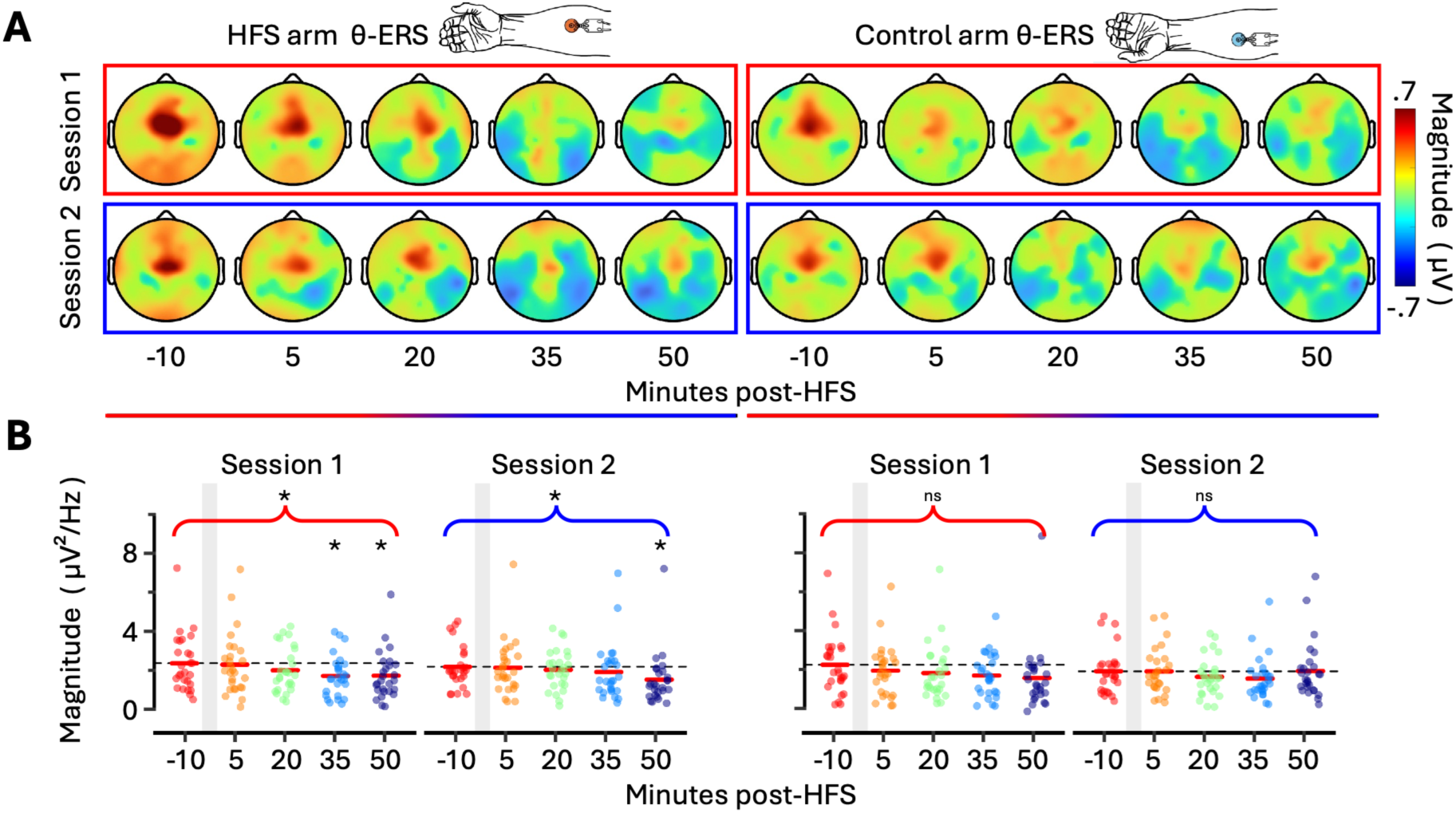
Electrical-evoked α-ERD results. a) ERSP topography plots represent the grand-averaged α-ERD across all participants for the HFS (left) and control (right) arms. Scalp topographies were averaged between 0ms to 500ms to capture the main α-ERD response between 8 Hz to 13 Hz. b) Dot plots represent individual α-ERD magnitudes displayed as raw data points alongside group means (red bars), for each time point, session, and arm separately. Statistics were derived from linear mixed-effects modelling and reflect change from baseline for each arm and session respectively. ^ns^ indicates no statistical significance (p > .05). * indicates significance at alpha = 0.05. HFS = High-frequency stimulation. α-ERD = alpha-band event-related desynchronisation.

Alternatively, for θ-ERS (Supplementary Table 8) significant negative time effects (Figure 12) were present for both the HFS (*F* (4, 243) = 8.2, *p* < .001) and control (*F* (4, 243) = 2.58, *p* = .04) arms. For the HFS arm this was driven by significant pairwise comparisons at 35 (*Std. effect* = -.86 [-1.5, -.23], *p* = .029) and 50 (*Std. effect* = -1 [-1.64, -.36], *p* = .008) minutes post-HFS during session 1, and at 50 (*Std. effect* = -1.22 [-1.86, -.58], *p* < .001) minutes post-HFS during session 2. For the control arm this was driven by a significant pairwise comparison at 50 (*Std. effect* = -.96 [-1.6, -.33], *p* = .012) minutes post-HFS during session 1.

**Figure 12.**
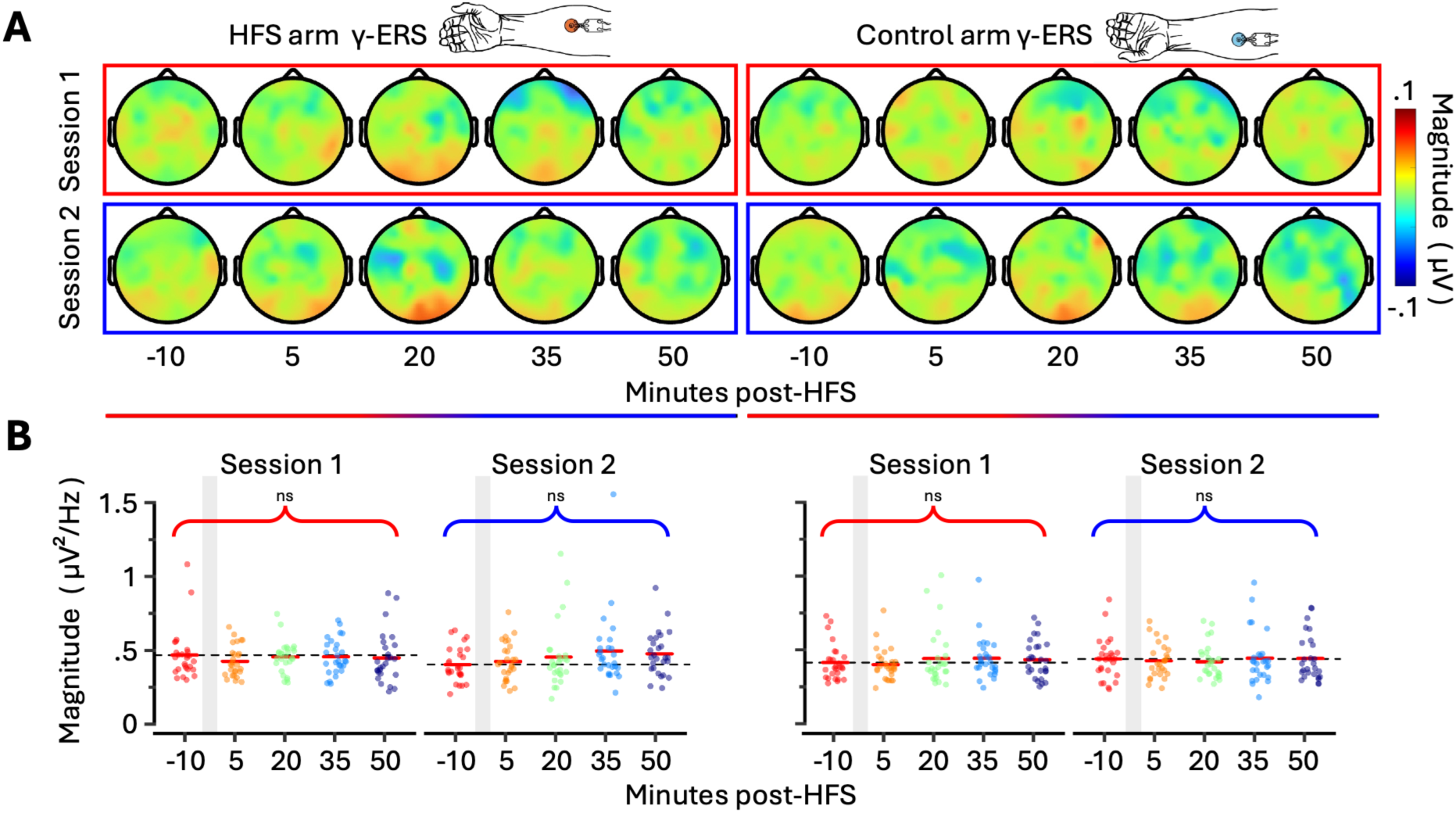
Electrical-evoked θ-ERS results. a) ERSP topography plots (top) represent the grand-averaged θ-ERS across all participants for the HFS (left) and control (right) arms. Scalp topographies were averaged between 0ms to 500ms to capture the main θ-ERD response between 4 Hz to 7 Hz. b) Dot plots (bottom) represent individual θ-ERS magnitudes displayed as raw data points alongside group means (red bars), for each time point, session, and arm separately. Statistics were derived from linear mixed-effects modelling and reflect change from baseline for each arm and session respectively. ^ns^ indicates no statistical significance (p > .05). * indicates significance at alpha = 0.05. HFS = High-frequency stimulation. θ-ERS = theta-band event-related synchronisation.

In addition, γ-ERS (Supplementary Table 9) demonstrated no significant main effects (Figure 13) were present for either arm, although a significant session-sex interaction (*F* (1, 243) = 10.41, *p* = .001) demonstrated greater fluctuations in γ-ERS between sessions for female participants (*F* (1, 38) = 5.87, *p* = .02) than for male participants (*F* (1, 38) = 4.35, *p* = .04).

**Figure 13.**
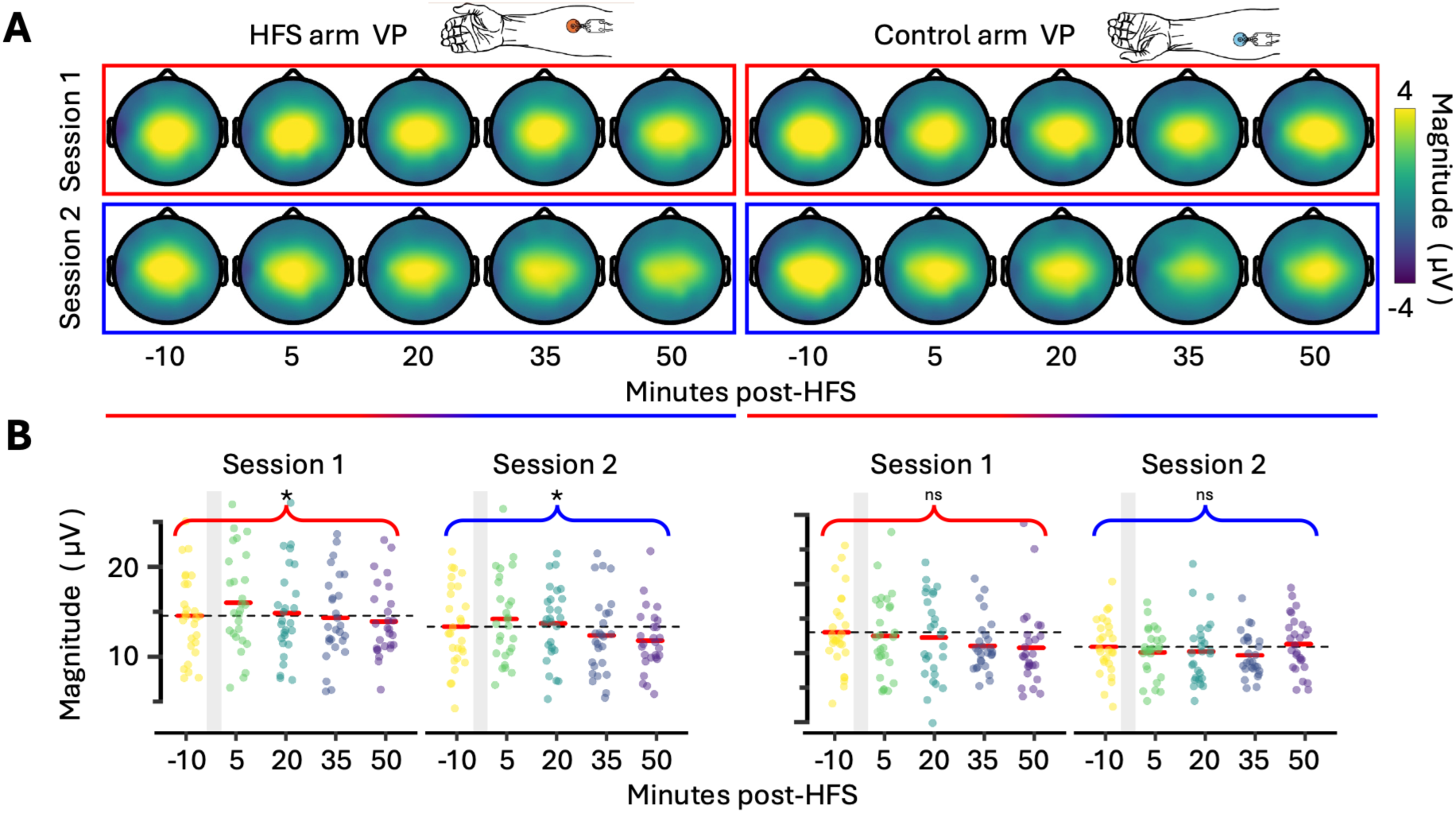
Electrical-evoked γ-ERS results. a) ERSP topography plots (top) represent the grand-averaged γ-ERS across all participants for the HFS (left) and control (right) arms. Scalp topographies were averaged between 0ms to 500ms to capture the main θ-ERD response between 50 Hz to 100 Hz. b) Dot plots (bottom) represent individual γ-ERS magnitudes displayed as raw data points alongside group means (red bars), for each time point, session, and arm separately. Statistics were derived from linear mixed-effects modelling and reflect change from baseline for each arm and session respectively. ^ns^ indicates no statistical significance (p > .05); * indicates significance at alpha = 0.05. HFS = High-frequency stimulation. γ-ERS = theta-band event-related synchronisation.

**Figure 14.**
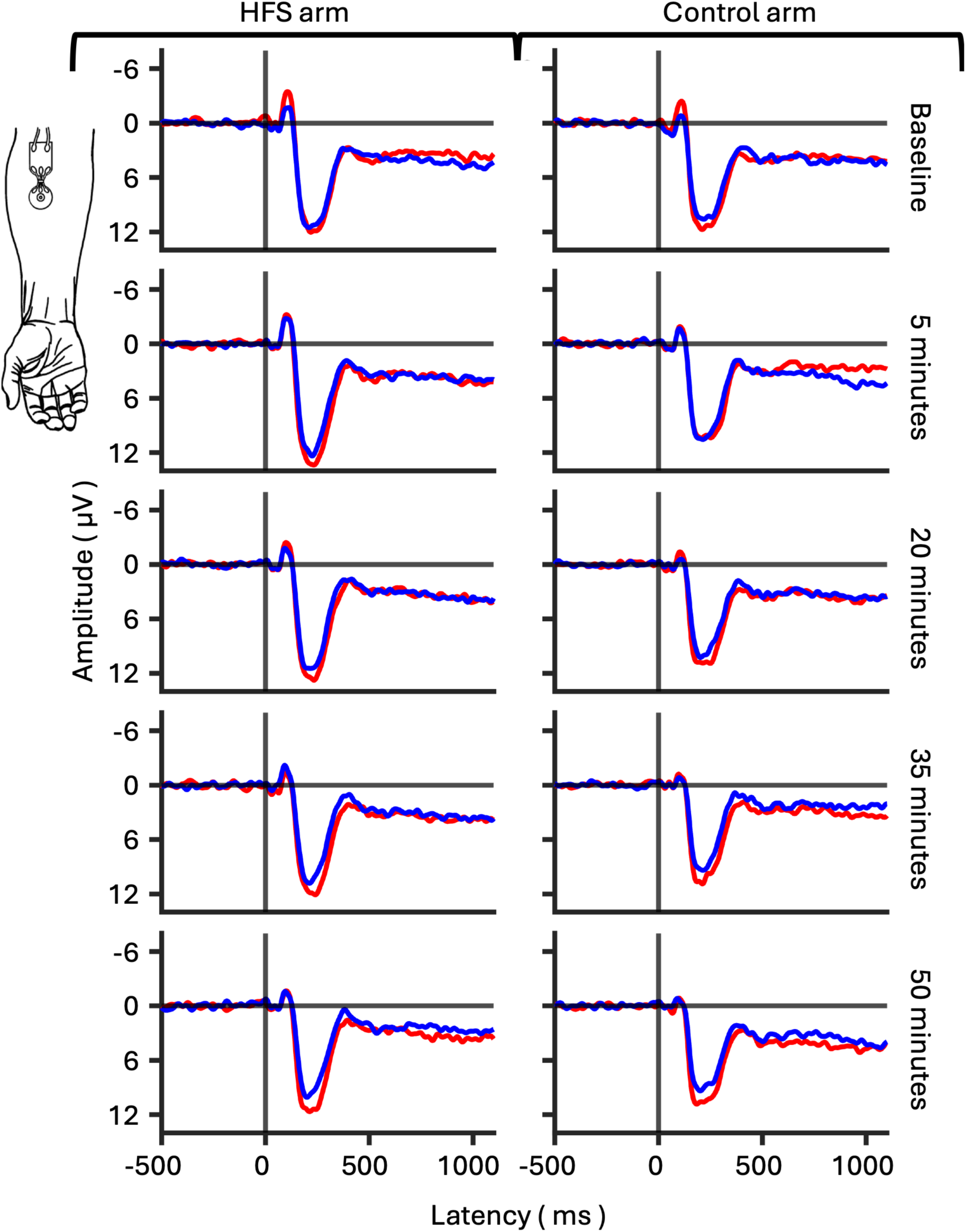
Electrical-evoked VP results. a) ERP plots represent the grand-averaged VP across all participants for the HFS (left) and control (right) arms. Statistics were derived from linear mixed-effects modelling and reflect change between sessions VP magnitude for each arm and time point respectively. b) Dot plots (bottom) represent individual VP magnitudes displayed as raw data points alongside group means (red bars), for each time point, session, and arm separately. Statistics were derived from linear mixed-effects modelling and reflect change from baseline VP magnitude for each arm and session respectively. ^ns^ indicates no statistical significance (p > .05). * indicates significance at alpha = 0.05. HFS = High-frequency stimulation. VP = Vertex potential. ERP = Event related potential.

**Figure 15.** Electrical-evoked VP ERP plots. ERP plots represent the grand-averaged VP at the vertex electrode (Cz) across all participants for both the HFS (left) and control (right) arms and sessions 1 (red) and 2 (blue). VP = vertex potential. HFS = High-frequency stimulation.

In contrast, VP magnitude (Supplementary Table 10 ) exhibited a significant negative session effect for both the HFS (*F* (1, 243) = 30.9, *p* < .001) and control (*F* (1, 243) = 14.34, *p* < .001) arms. For the HFS arm this was driven by multiple significant between-sessions pairwise comparisons at 5 (*Std. effect* = -.93 [-1.57, -.3], *p* = .004), 35 (*Std. effect* = -.79, [- 1.43, -.16], *p* = .015), and 50 (*Std. effect* = -.84 [-1.48, -.21], *p* = .010) minutes post-HFS. For the control arm this was driven by a significant between-sessions pairwise comparison at 50 minutes post-HFS (*Std. effect* = -.73 [-1.36, -.09], *p* = .03). Interestingly, a significant interaction between session and sex for the HFS arm (*F* (1, 243) = 4.12, *p* = .04) indicates that only female participants exhibited a significant change between sessions (*F* (4, 47) = 5.06, *p* = .002), whilst male participants did not (*F* (4, 47) = .79, *p* = .54). Similarly, a significant sex effect (*F* (1, 27) = 7.2, *p* = .01) was also evident for the control arm as female participants consistently exhibited larger VP amplitudes, although the interaction between session and sex was not significant (*F* (1, 243) = 2.37, *p* = .13). Furthermore, a significant time effect (*F* (4, 243) = 3.94, *p* = .004) was also present for the HFS arm, although no change from baseline pairwise comparisons were significant.

Alternatively, VP latency (Supplementary Table 11) fluctuated over time for both the HFS (*F* (4, 243) = 2.89, *p* = .02) and control arms (*F* (4, 243) = 2.42, *p* = .05) in a manner that was substantially influenced by demographical factors for the HFS arm as a significant interaction between age and time was evident (*F* (4, 243) = 3.61, *p* = .01). Furthermore, a significant interaction between age and session (*F* (1, 243) = 6.65, *p* = .01) was present for the HFS arm, indicating that older participants showed greater changes between sessions with the threshold of significance at > 45 years.

Overall, electrical-evoked EEG responses varied widely across time and between sessions in a manner that was often characterised by individual differences in demographical factors.

### Relationships between electrical-evoked EEG responses and EPP

To investigate the relationship between electrical-evoked EEG responses and reported pain intensity, mixed-effects modelling was used to characterise changes in EPP as previously (Mugglestone et al., 2026), with the addition of baseline and change from baseline EEG data included as covariates.

Across electrical-evoked EEG responses, VP magnitude was directly associated with EPP, reflecting both intra- and inter-individual variations through significant baseline and change from baseline effects. Alternatively, the influence of baseline ERSP measures trended towards significance. For all models, a significant effect of session (all *F* (1, 514) > 77.88, all *p* < .001) was present, although no significant effect of time or arm was evident, demonstrating the lack of homotopic sensitivity post-HFS as previously reported (Mugglestone et al., 2026).

For α-ERD, no significant influence of either baseline (*F* (1, 536) = 3.69, *p* = .055) change from baseline (*F* (1, 521) = .01, *p* = .93) was evident. Furthermore, θ-ERS also demonstrated no significant main effects for either baseline (*F* (1, 531) = 3.76, *p* = .053) or change from baseline (*F* (1, 519) = 2.05, *p* = .15). However, a significant interaction between session and baseline θ-ERS was present (*F* (1, 514) = 24.49, *p* < .001), suggesting that changes in EPP between sessions were reflected by inter-individual differences in θ-ERS magnitude.

In addition, both baseline (*F* (1, 522) = 19.6, *p* < .001) and change from baseline (*F* (1, 514) = 3.99, *p* = .046) γ-ERS were significantly related to EPP, suggesting that γ-ERS reflected intra- and inter-individual changes in reported pain intensity between repeated test sessions and over time. However, a significant interaction between γ-ERS change from baseline and arm (*F* (1, 514) = 7.04, *p* = .008) was also evident, potentially presenting a lateralised difference in the accuracy of γ-ERS for reflecting EPP, as the influence of change from baseline γ-ERS on EPP was only significant for the control arm (*β* = .55, *p* = .002) and not the HFS arm (*β* = .06, *p* = .7). Furthermore, a significant interaction between session and baseline γ-ERS magnitude was evident (*F* (1, 514) = 29.06, *p* < .001), demonstrating that inter-individual variations in EPP between sessions were characterised by altered γ-ERS magnitudes.

Alternatively, there were significant relationships between electrical-evoked VP magnitude and EPP both for baseline (*F* (1, 530) = 17.46, *p* < .001) and change from baseline (*F* (1, 516) = 6.41, *p* = .01), suggesting that fluctuations in reported pain intensity between repeated test sessions and over time were reflected by changes in VP magnitude. Furthermore, a significant interaction between session and baseline VP magnitude was evident (*F* (1, 514) = 10.21, *p* = .001), demonstrating that inter-individual variations in EPP between sessions were characterised by altered VP magnitudes. Finally, no significant effects of VP latency were evident for baseline (*F* (1, 522) = 1.07, *p* = .3) or change from baseline (*F* (1, 517) = 0, *p* =.96).

Overall, VP magnitude demonstrated the strongest relationship with EPP, capturing both inter- and intra-individual variations through baseline and change from baseline effects respectively. In contrast, both baseline α-ERD and θ-ERS only trended towards a significant relationship with EPP. However, whilst these relationships may suggest that ERP and ERSP measures can capture inter- and intra-individual differences in reported pain intensity, given the lack of behavioural homotopic sensitivity development, their sensitivity to hyperalgesic changes cannot be fully characterised.

## Discussion

The present study investigated the temporal dynamics of HFS-induced changes to evoked EEG across repeated test sessions. Pinprick-evoked responses were of primary interest given the behavioural development of SHA as previously reported (Mugglestone et al., 2026). Strengthened α-ERD indicated altered somatosensory processing post-HFS, and baseline α-ERD was related to individual differences in the subsequent development of SHA. Additionally, across both electrical and pinprick stimuli, general sensory-related cortical activity assessed through θ-ERS captured differences in reported pain intensity between sessions, whilst individual variations were reflected by VP magnitude, potentially as a product of extralemniscal activity. However, neither VP magnitude or latency, nor θ-ERS, were sensitive to the HFS-induced changes in MPS that indicated of SHA development. This discrepancy may reflect methodological limitations of the handheld pinprick device and warrants further investigation. Furthermore, γ-ERS responses were only evident for electrical stimuli, although they were effective for characterising EPP. All evoked EEG data demonstrated poor-to-moderate test-retest reliability at baseline, with lateralised differences often present. Furthermore, extremely poor reliability was evident when post-HFS change-from-baseline was assessed. Therefore, whilst evoked EEG is useful for clarifying the cortical mechanisms involved in induced sensitivity changes and can provide insight into individual differences in reported pain intensity both within and between sessions, caution is advised when considering the use of these features for interventional testing.

Test arm pinprick-evoked VP magnitude was not significantly altered post-HFS, suggesting that the specificity of nociceptive-evoked VPs is insufficient for characterising HFS-induced SHA (Leone et al., 2025). Previously, significant variations in post-sensitisation VP magnitudes have been demonstrated when an initially sub-painful stimulus was utilised, indicating an allodynic-like effect (Biurrun Manresa et al., 2018; van den Broeke et al., 2020; van den Broeke et al., 2017; Van den Broeke et al., 2016; van den Broeke et al., 2015). Alternatively, higher intensity stimuli necessary to elicit genuinely hyperalgesic responses, such as the 128 mN pinprick used here, often do not exhibit post-HFS alterations (Torta et al., 2020; van den Broeke et al., 2015).

Furthermore, VP magnitude is considered to be driven by extralemniscal activity and may reflect differential rather than absolute stimulus intensity as a product of salience (L Hu et al., 2014; Hu & Iannetti, 2019; Hu et al., 2010; Iannetti et al., 2005; Iannetti et al., 2008; Jin et al., 2018; Lee et al., 2009; Somervail et al., 2025; Somervail et al., 2021; Squintani et al., 2021; Valentini et al., 2012; Wang et al., 2023; Zhang et al., 2021; Zidda et al., 2024). Therefore, these data suggest that VP magnitude may be ineffective for characterising SHA as the extralemniscal response to an already painful stimulus may not be substantially enhanced post-HFS (Leone et al., 2025; Somervail et al., 2021). Indeed, van den Broeke et al. (2015) have demonstrated that changes to pinprick-evoked VP amplitudes following SHA induction can follow a gaussian curve pattern, whereby stimuli on the threshold of painfulness prior to sensitisation show the largest increase in VP amplitude. However, as van den Broeke et al. (2015) used capsaicin to induce SHA rather than HFS, this may be tied to the relative salience of pinprick stimuli compared to the ongoing tonic pain sensations caused by capsaicin application. Therefore, future work is required to replicate this pattern within a HFS experimental paradigm.

For both pinprick and electrical stimuli, change-from-baseline in VP magnitude was related to reported pain intensity across arms, supporting the hypothesis that the VP can reflect intra-individual differences in pain (Tiemann et al., 2026), which may reflect variations in extralemniscal activity (Perovic et al., 2025; Somervail et al., 2025; Somervail et al., 2021). Furthermore, baseline electrical-evoked VP magnitude also exhibited a significant relationship with EPP, indicating that larger inter-individual extralemniscal responses reflected greater reported pain intensity and smaller change between sessions (Lee et al., 2009; Somervail et al., 2025; Somervail et al., 2021; van den Broeke et al., 2015), and that the VP may be sensitive to inter-individual variations in pain perception (Tiemann et al., 2026). However, this effect was absent for pinprick-evoked VP magnitude, which may be explained by the comparatively weaker VP waveform relative to electrical stimulation, likely caused by variations in stimuli durations and force profiles as a handheld mechanical pinprick device was used (Triccas et al., 2022; van den Broeke et al., 2020; van den Broeke et al., 2017). Future work should therefore employ more controlled methods of mechanical stimulation to obtain sharper and more reliable VP waveforms, which may better predict reported pain intensity and HFS-induced MPS (see van den Broeke et al. (2020) for a description of a robot-controlled device). However, multivariate analyses linking VP magnitude to HFS-induced mechanical pinprick sensitivity within such data have yet to be reported. Furthermore, although electrical-evoked VP magnitude presented a more robust waveform, previous work has demonstrated that single-pulse electrical stimuli do not engage the peripheral afferent fibres sensitive to HFS-induced primary hyperalgesia, limiting the utility of this finding (Biurrun Manresa et al., 2018; Leone et al., 2025; Mugglestone et al., 2026; Poulsen et al., 2022; E. van den Broeke et al., 2021; E. N. Van den Broeke et al., 2021).

Alternatively, VP latency did not vary significantly between session or across arms for either nociceptive modality. This suggests that the pattern of post-stimulus cortical engagement was stable, potentially reflecting a lack of change in the timing of extralemniscal engagement across experimental sessions (Lopez-Calderon & Luck, 2014; Somervail et al., 2025). Alternatively, electrical-evoked VP latency decreased over time for both arms, beginning at 20 minutes post-HFS, potentially demonstrating an increased speed of cortical processing. However, this effect was not present for pinprick stimuli, which may be due to the sharper and more reliable electrical-evoked VP waveforms as discussed above.

Unlike the VP, electrical-evoked θ-ERS significantly decreased post-HFS for both arms, denoting a blunted cortical response which may reflect a general habituation effect that was not localised to HFS-specific response due to the impact of repeated non-sensitised stimulation on attentional demands. θ-ERS represents the sum of early-latency ERPs arising from widespread cortical generators present within both the lemniscal and extralemniscal systems (Hu & Iannetti, 2019; Ploner et al., 2017; Zhang et al., 2021). Therefore, this may reflect a general change in lemniscal processing when contrasted with the lack of extralemniscal changes demonstrated by the VP (Hu & Iannetti, 2019; Michail et al., 2016; Zhang et al., 2021). However, this effect was absent for pinprick stimuli, limiting the relevance of θ-ERS for investigating SHA development. Despite no direct relationship between θ-ERS and reported pain intensity, baseline θ-ERS magnitude was significantly related to the between-session change in reported pain intensity for both nociceptive modalities across arms. This suggests that inter-individual responses in θ-ERS magnitude reflected differences in reported pain intensity between sessions. Therefore, despite its non-specificity for HFS-induced SHA, θ-ERS may be useful for characterising inter-individual differences in the stability of pain perception, providing a marker of altered cortical activity related to the reduction in pain intensity present within session 2, which was previously unattributed after considering typical psychophysical factors (Mugglestone et al., 2026).

Alternatively, electrical-evoked γ-ERS responses were somewhat effective for characterising individual differences in EPP, reflecting inter-individual changes to reported pain intensity between sessions for both arms and intra-individual differences over time for the control arm only. This suggests that γ-ERS responses may be most effective for characterising longer term variability in pain perception (Tiemann et al., 2026). However, no robust pinprick-evoked γ-ERS responses were evident, perhaps because γ-ERS is primarily exhibited outside of mechano-nociceptive contexts (Gousset et al., 2023; Linde et al., 2023; Yue et al., 2025). Therefore, these data reinforce the limitations of γ-ERS for depicting MPS SHA (Gousset et al., 2023).

Test-arm α-ERD was strengthened post-HFS for both nociceptive modalities, consistent with increased sensorimotor cortical engagement (Hu et al., 2013; Ploner et al., 2017; Triccas et al., 2022) and pain sensitivity (Harrison et al., 2025). Despite visual variability, post-HFS changes to control-arm α-ERD were not statistically significant, supporting the specificity of test-arm α-ERD reduction to altered cortical processing post-HFS, which may therefore reflect a key mechanism of pain sensitivity development (Harrison et al., 2025; Hu et al., 2013; Ploner et al., 2017), consistent with recent fMRI evidence linking somatosensory cortex activity to acute nociceptive sensitisation (Harrison et al., 2025). However, as altered test-arm α-ERD was present for both pinprick and electrical stimuli regardless of whether behavioural hyperalgesia was evident, this suggests that HFS provoked alterations to the somatotopic localisation of nociceptive information were not specific to a single noxious modality (Ploner et al., 2017).

In addition, whilst the change in α-ERD magnitude did not directly characterise pain intensity, inter-individual variations in pinprick-evoked α-ERD at baseline were related to post-HFS changes in MPS over time, with stronger baseline α-ERD denoting a steeper slope of change. Therefore, these data suggest that the magnitude of pinprick-evoked cortical activity prior to HFS-conditioning can influence the time course of SHA development. This supports the hypothesis that α-ERD does not directly reflect subjective pain intensity but likely signals the spatial integration and feed-back processing of nociceptive information as advocated by Ploner et al. (2017).

However, as α-ERD and θ-ERS both demonstrated reduced magnitudes post-HFS, a general decrease in spectral power must be considered if the assumption that these reflect real changes to specific periodic components is to be justified. Critically, the present study applied SPRiNT (Wilson et al., 2022) to separate the aperiodic 1/f slope (Donoghue et al., 2020) before assessing the magnitude of periodic components. Additionally, a direct subtraction approach was applied, assuming an additive model of EEG (Gyurkovics et al., 2021). Therefore, the authors are confident that the reported ERSP data reflect changes in the scale of oscillatory activity evoked by nociceptive stimulation, and that a general effect is ruled out (Donoghue et al., 2020; Gyurkovics et al., 2021; Li Hu et al., 2014; Rawls & Sponheim, 2025; Wilson et al., 2022). For further justification of these methods see Donoghue et al. (2020), Wilson et al. (2022), and Gyurkovics et al. (2021). Furthermore, α-ERD and θ-ERS were not significantly correlated (r = -.03, t (1078) = -.96, *p* = .34), suggesting that their changes were independent. This is further supported by the present linear mixed-effects modelling which demonstrated that reduced α-ERD magnitude occurred post-HFS for both noxious modalities on the test arm only, whilst reduced θ-ERS magnitude was evident across arms, but only for electrical stimuli, suggesting distinct cortical components.

Whilst the evoked-EEG measures discussed above provide insight into individual variations in the cortical processing of nociceptive stimuli and their specificity for investigating HFS-induced hyperalgesia, all baseline ICC values were below .75, failing to reach good test-retest reliability (Koo & Li, 2016). This is further evidenced by significant between-session variations both pre- and post-HFS, alongside multiple cases in which change-from-baseline test-retest reliability was extremely poor demonstrating greater within-individuals variation than between-individuals. Bland-Altman analyses showed small systematic biases, however, large limits of agreement relative to observed ranges suggest that these evoked EEG measures were generally insufficient for tracking individual responses across sessions. This was further pronounced when change-from-baseline values were assessed. However, these significant between-sessions variabilities mirror the behavioural decrease in reported pain intensity previously reported (Mugglestone et al., 2026), which was evident for both noxious modalities across arms. Therefore, poor test-retest reliability may partly reflect sensitivity to within-individual changes across sessions. Subsequently, these EEG features may be better suited for capturing variability in nociceptive processing than presenting stable biomarkers for use in interventional studies.

In conclusion, these data provide the temporal dynamics of HFS-induced changes to nociceptive-evoked EEG responses across repeated test sessions and demonstrate their utility for understanding variability in HFS-induced changes in pain perception. Whilst pinprick-evoked VP magnitude was not altered by the development of behavioural SHA, individual differences reflected variations in reported pain intensity across noxious modalities and arms. Comparatively, θ-ERS was related to between-session variability in pain perception, whilst baseline α-ERD presented the only significant predictive relationship with subsequent SHA development. Overall, these results suggest that evoked EEG features can be used to investigate individual differences in pain processing and the temporal dynamics of HFS-induced SHA, however, they lack sufficient reliability and precision to serve as robust measures or correlates of SHA across experimental sessions.

## Disclosures

This work is funded through a PhD studentship from the University of Plymouth (School of Psychology) and an Academy of Medical Sciences Springboard Award (Hughes: SBF007\100108). The authors have no conflicts of interest.

## Authorship Contributions

SM: Conceptualisation, funding acquisition, data collection, formal analysis, writing, reviewing and editing.

GG: Conceptualisation, funding acquisition, supervision, methodology, reviewing and editing.

SH: Conceptualisation, funding acquisition, supervision, methodology, reviewing and editing.

## Data availability statement

Data are available upon request.

## Notes

### Competing Interest Statement

The authors have declared no competing interest.

