## Supplementary Materials for "Nociceptive-evoked electroencephalography can predict inter-individual differences in high frequency induced sensitisation of central nociceptive pathways: A test-retest reliability study"

Supplementary Table 1

**Pinprick-evoked EEG descriptive statistics and assessments of reliability and agreement**

| **Key:** | Value (standard deviation) or [95% confidence interval] | | | | |
|---|---|---|---|---|---|
| | **Raw data** | **Change from baseline (minutes)** | | | |
| **Time post-HFS** | Baseline | 5 | 20 | 35 | 50 |
| **Vertex potential magnitude (μV)** | | | | | |
| **HFS arm** | | | | | |
| Session 1 mean | 6.8 (3.5) | 0 (2.9) | -.3 (3) | .3 (2.6) | 0 (2.5) |
| Session 1 range | 2 – 15 | -4.2 – 8 | -6.7 – 6 | -3.9 – 6 | -5.3 – 6 |
| Session 2 mean | 5.4 (3) | .2 (2.3) | .6 (2.3) | .2 (2.6) | 0 (3.3) |
| Session 2 range | -.2 – 10 | -4.5 – 5 | -3.9 – 5 | -3.4 – 8 | -6.4 – 6 |
| ICC | .41 [.04 .7] | .26 [-.1 .6] | .23 [-.2 .6] | .55 [.2 .8] | .25 [-.1 .6] |
| Bias | 1.5 [.1 2.9] | -.2 [-1.5 1.1] | -.9 [-2.2 .4] | .1 [-.89 1.1] | .02 [-1.4 1.5] |
| Upper | 8.4 [6 10.9] | 6.1 [3.9 8.4] | 5.6 [3.3 7.8] | 5 [3.3 6.7] | 7.1 [4.6 9.6] |
| Lower | -5.5 [-8 -3.1] | -6.5 [-8.7 -4.3] | -7.4 [-9.6 -5.1] | -4.8 [-6.5 -3.1] | -7.1 [-9.5 -4.6] |
| **Control arm** | | | | | |
| Session 1 mean | 6.5 (2.8) | -.3 (2.7) | -.4 (2.3) | -1 (2) | -1.1 (2.3) |
| Session 1 range | 1.3 – 13 | -5.3 – 6 | -4.1 – 4 | -4.5 – 4 | -4.9 – 3 |
| Session 2 mean | 5.5 (2) | -.4 (1.6) | -.3 (2.1) | -.6 (2.4) | .2 (2.2) |
| Session 2 range | 1.1 – 10 | -3.6 – 2 | -5.1 – 4 | -6.2 – 5 | -3.0 – 4 |
| ICC | .5 [.2 .7] | .25 [-.1 .6] | .0 [-.4 .4] | .23 [-.2 .6] | .01 [-.4 .4] |
| Bias | 1.1 [.1 2] | .1 [-1 1.2] | -.03 [-1.5 1.4] | -.4 [-1.5 .7] | -1.3 [-2.6 -.1] |
| Upper | 5.9 [4.2 7.6] | 5.5 [3.7 7.4] | 7.1 [4.6 9.5] | 5 [3.2 6.9] | 4.8 [2.7 7] |
| Lower | -3.8 [-5.5 -2.1] | -5.3 [-7.1 -3.4] | -7.1 [-9.6 -4.6] | -5.8 [-7.7 -3.9] | -7.5 [-9.6 -5.3] |

Supplementary Table 1 continued

**Pinprick-evoked EEG descriptive statistics and assessments of reliability and agreement**

| **Vertex potential latency** (ms) | | | | | |
|---|---|---|---|---|---|
| **HFS arm** | | | | | |
| Session 1 mean | 253.9 (116.7) | -4.2 (129.7) | -16.1 (124.9) | -20.8 (118.3) | 7.3 (136.7) |
| Session 1 range | 100 – 498 | -248 – 284 | -250 – 344 | -306 – 320 | -238 – 328 |
| Session 2 mean | 260.5 (119.6) | -22.7 (137.7) | 27.6 (172.1) | 10.8 (108) | -13.4 (144.8) |
| Session 2 range | 112 – 486 | -358 – 310 | -254 – 374 | -298 – 244 | -304 – 302 |
| ICC | .16 [-.2 .5] | 0 [-.4 .4] | .1 [-.3 .4] | 0 [-.4 .4] | 0 [-.4 .4] |
| Bias | -6.7 [-68 55] | 18.4 [-62 99] | -43.7 [-127 40] | -31.6 [-103 40] | 20.7 [-67.3 108] |
| Upper | 298.8 [192 405] | 417.6 [278 557] | 370.8 [226 515] | 325.1 [200 449] | 456.6 [304 608] |
| Lower | -312.1 [-418 -205] | -380.7 [-520 -241] | -458.3 [-603 -313] | -388.4 [-513 -263] | -415.3 [-567 -263] |
| **Control arm** | | | | | |
| Session 1 mean | 241.3 (99.3) | -20.7 (129.5) | 24.5 (118.7) | 7.9 (129.2) | -11.4 (120.9) |
| Session 1 range | 78 – 464 | -308 – 360 | -206 – 380 | -312 – 282 | -284 – 292 |
| Session 2 mean | 249.4 (111.8) | .4 (168.9) | 18.1 (153.5) | -5.9 (121.7) | 15.5 (159.7) |
| Session 2 range | 116 – 496 | -280 – 362 | -290 – 384 | -314 – 172 | -338 – 294 |
| ICC | .29 [-.1 .6] | 0 [-.4 .4] | 0 [-.4 .4] | .1 [-.3 .4] | .15 [-.2 .5] |
| Bias | -8.1 [-62.6 46.5] | -21.2 [-117 75] | 6.4 [-70 83] | 13.8 [-56 83] | -26.9 [-99 45] |
| Upper | 262.1 [167 356] | 455.8 [289 622] | 388.1 [254 521] | 360.9 [239 482] | 332.6 [207 458] |
| Lower | -278.2 [-372 -183] | -498.2 [-664 -331] | -375.3 [-508 -242] | -333.4 [-454 -212] | -386.4 [-512 -260] |

Supplementary Table 1 continued

**Pinprick-evoked EEG descriptive statistics and assessments of reliability and agreement**

| **α-ERD magnitude** (μV²/Hz) | | | | | |
|---|---|---|---|---|---|
| **HFS arm** | | | | | |
| Session 1 mean | -1.3 (1.1) | -.3 (.7) | -.7 (1.7) | -.4 (1.1) | -.8 (1.5) |
| Session 1 range | -3.8 – -.1 | -2 – 1 | -6 – 1 | -3.1 – 2 | -5.7 – 1 |
| Session 2 mean | -1.8 (1.5) | -.2 (1.8) | -.4 (2.2) | -.4 (1.3) | -.3 (2.1) |
| Session 2 range | -5.8 – -.1 | -5.6 – 3 | -9.1 – 3 | -3.6 – 2 | -7.2 – 4 |
| ICC | .58 [.3 .8] | 0 [-.4 .4] | 0 [-.4 .4] | 0 [-.4 .4] | .26 [-.1 .6] |
| Bias | .5 [0 .9] | -.07 [-.9 .8] | -.3 [-1.5 .8] | -0 [-.8 .8] | -.5 [-1.4 .4] |
| Upper | 2.8 [2 3.6] | 4 [2.6 5.4] | 5.2 [3.3 7.2] | 3.7 [2.4 5] | 3.9 [2.4 5.4] |
| Lower | -1.8 [-2.7 -1] | -4.1 [-5.5 -2.7] | -5.9 [-7.8 -3.9] | -3.7 [-5 -2.4] | -4.9 [-6.4 -3.4] |
| **Control arm** | | | | | |
| Session 1 mean | -1.3 (1.1) | -.2 (0.9) | -1 (2.7) | -.9 (1.6) | -.9 (2.3) |
| Session 1 range | -4.9 – -.1 | -2.8 – 2 | -9.9 – 5 | -5.7 – 1 | -7.1 – 5 |
| Session 2 mean | -1.9 (2.3) | .3 (1.6) | -.1 (2.2) | 0 (2.4) | -.1 (1.6) |
| Session 2 range | -9.4 – -.1 | -2.6 – 4 | -6.6 – 6 | -6.8 – 6 | -5.3 – 4 |
| ICC | .28 [-.1 .6] | .03 [-.4 .4] | 0 [-.4 .4] | .04 [-.3 .4] | 0 [-.4 .4] |
| Bias | .6 [-.2 1.5] | -.5 [-1.2 .3] | -.9 [-2.4 .6] | -.9 [-2 .3] | -.8 [-1.9 .3] |
| Upper | 4.8 [3.4 6.3] | 3 [1.8 4.3] | 6.4 [3.8 8.9] | 4.6 [2.7 6.6] | 4.6 [2.7 6.5] |
| Lower | -3.5 [-5 -2.1] | -4 [-5.2 -2.7] | -8.2 [-10.7 -5.6] | -6.4 [-8.3 -4.4] | -6.3 [-8.2 -4.4] |

Supplementary Table 1 continued

**Pinprick-evoked EEG descriptive statistics and assessments of reliability and agreement**

| **θ-ERS magnitude** ($\mu V^2/Hz$) | | | | | |
|---|---|---|---|---|---|
| **HFS arm** | | | | | |
| Session 1 mean | 1.4<br>(.9) | -.1<br>(.7) | -.1<br>(.8) | -.4<br>(.7) | -.3<br>(.9) |
| Session 1 range | .1 – 5 | -2. – 1 | -2.1 – 1 | -1.8 – 1 | -2.4 – 1 |
| Session 2 mean | 1<br>(.8) | .2<br>(.9) | .3<br>(1) | .3<br>(1.1) | .2<br>(1) |
| Session 2 range | -.1 – 3 | -2.2 – 2 | -2 – 3 | -2.1 – 2 | -1.7 – 2 |
| ICC | .59<br>[.3 .8] | .22<br>[-.2 .6] | .41<br>[.1 .7] | .25<br>[-.1 .6] | .06<br>[-.3 .4] |
| Bias | .4<br>[.1 .7] | -.3<br>[-.7 .1] | -.4<br>[-.8 0] | -.7<br>[-1.1 -.3] | -.6<br>[-1.1 -.1] |
| Upper | 2<br>[1.4 2.5] | 1.7<br>[1 2.4] | 1.6<br>[.9 2.2] | 1.5<br>[.7 2.3] | 2<br>[1.1 2.9] |
| Lower | -1.2<br>[-1.8 -.7] | -2.3<br>[-3.1 -1.6] | -2.4<br>[-3 -1.7] | -2.9<br>[-3.7 -2.1] | -3.1<br>[-4 -2.2] |
| **Control arm** | | | | | |
| Session 1 mean | 1.2<br>(.8) | -.1<br>(.8) | .2<br>(1.3) | -.2<br>(.7) | -.1<br>(.9) |
| Session 1 range | -.1 – 3 | -1.9 – 1 | -1.4 – 5 | -1.6 – 1 | -1.4 – 2 |
| Session 2 mean | .9<br>(.6) | 0<br>(.6) | .1<br>(1) | .2<br>(.8) | .4<br>(1.5) |
| Session 2 range | .1 – 2 | -1.4 – 1 | -1.5 – 3 | -1.5 – 2 | -1.3 – 7 |
| ICC | .57<br>[.3 .8] | .19<br>[-.2 .5] | .46<br>[.1 .7] | .15<br>[-.2 .5] | .33<br>[-.1 .6] |
| Bias | .3<br>[0 .5] | -.1<br>[-.5 .3] | .1<br>[-.4 .6] | -.3<br>[-.7 .1] | -.5<br>[-1.1 0] |
| Upper | 1.5<br>[1.1 2] | 1.7<br>[1.1 2.3] | 2.5<br>[1.6 3.3] | 1.6<br>[.9 2.3] | 2.2<br>[1.3 3.2] |
| Lower | -1<br>[-1.4 -.5] | -1.9<br>[-2.5 -1.3] | -2.3<br>[-3.2 -1.5] | -2.3<br>[-3 -1.6] | -3.3<br>[-4.3 -2.3] |

*Supplementary Table 1 Pinprick-evoked EEG descriptive statistics and assessments of reliability and agreement*

Mean values, standard deviations, and ranges were calculated separately for each session, time point, arm, and stimulation modality. Change from baseline scores were calculated as the absolute difference from baseline by subtracting the baseline values for each EEG response from their post-HFS counterparts. ERD = event-related desynchronisation. ERS = event-related synchronisation. HFS = High-frequency stimulation. N = 27, (female = 19).

Supplementary Table 2

**Pinprick-evoked α-ERD magnitude results**

| | | | | |
|---|---|---|---|---|
| **a) Model assessment: HFS arm** | | | | |
| Magnitude ~ time * session * (age + sex) + (1 \| subject) | | | | |
| Conditional $R^2$ = .66 | | Marginal $R^2$ = .06 | | |
| **Fixed effects and interactions** | | | | |
| Predictor | | *F* (DF) | *η2* [95% CI] | *p* |
| Time | | 4.73 (4, 243) | 0.07 [0.02, 1] | 0.001 |
| Session | | 8.17 (1, 243) | 0.03 [0.01, 1] | 0.005 |
| Age | | 0.07 (1, 27) | <.01 [0, 1] | 0.80 |
| Sex | | 0 (1, 27) | <.01 [0, 1] | 0.94 |
| Time: session | | 0.23 (4, 243) | <.01 [0, 1] | 0.92 |
| Time: age | | 0.46 (4, 243) | <.01 [0, 1] | 0.76 |
| Time: sex | | 1.32 (4, 243) | 0.02 [0, 1] | 0.26 |
| Session: age | | 2.83 (1, 243) | 0.01 [0, 1] | 0.09 |
| Session: sex | | 0.6 (1, 243) | <.01 [0, 1] | 0.44 |
| Time: session: age | | 1.56 (4, 243) | 0.03 [0, 1] | 0.18 |
| Time: session: sex | | 0.93 (4, 243) | 0.02 [0, 1] | 0.45 |
| **Between session comparisons: Session 2 – Session 1** | | | | |
| Arm | Minutes post-HFS | Estimate [95% CI] | Std. effect [95% CI] | *P* |
| HFS | -10 | -0.16 [-0.53, 0.21] | -0.28 [-0.91, 0.35] | 0.38 |
| HFS | 5 | -0.33 [-0.7, 0.04] | -0.56 [-1.2, 0.07] | 0.08 |
| HFS | 20 | -0.19 [-0.56, 0.18] | -0.32 [-0.95, 0.32] | 0.32 |
| HFS | 35 | -0.3 [-0.67, 0.07] | -0.51 [-1.14, 0.13] | 0.11 |
| HFS | 50 | -0.14 [-0.51, 0.23] | -0.24 [-0.87, 0.39] | 0.45 |
| **Session 1 comparisons to baseline: Post-HFS – Baseline** | | | | |
| Arm | Minutes post-HFS | Estimate [95% CI] | Std. effect [95% CI] | P |
| HFS | 5 | -0.12 [-0.58, 0.35] | -0.2 [-0.83, 0.43] | 0.88 |
| HFS | 20 | -0.42 [-0.88, 0.05] | -0.71 [-1.35, -0.08] | 0.09 |
| HFS | 35 | -0.25 [-0.72, 0.22] | -0.43 [-1.06, 0.21] | 0.47 |
| HFS | 50 | -0.48 [-0.95, -0.02] | -0.82 [-1.46, -0.19] | 0.04 |
| **Session 2 comparisons to baseline: Post-HFS – Baseline** | | | | |
| Arm | Minutes post-HFS | Estimate [95% CI] | Std. effect [95% CI] | *P* |
| HFS | 5 | -0.28 [-0.75, 0.18] | -0.48 [-1.12, 0.15] | 0.37 |
| HFS | 20 | -0.44 [-0.91, 0.03] | -0.75 [-1.39, -0.11] | 0.07 |
| HFS | 35 | -0.38 [-0.85, 0.08] | -0.66 [-1.29, -0.02] | 0.14 |
| HFS | 50 | -0.46 [-0.92, 0.01] | -0.78 [-1.42, -0.15] | 0.06 |

Supplementary Table 2 continued

**Pinprick-evoked α-ERD magnitude results**

| | | | | |
|---|---|---|---|---|
| **b) Model assessment: Control arm** | | | | |
| Magnitude ~ time * session * (age + sex) + (1 \| subject) | | | | |
| Conditional $R^2$ = .58 | | Marginal $R^2$ = .07 | | |
| **Fixed effects and interactions** | | | | |
| Predictor | | *F* (DF) | *η2* [95% CI] | *P* |
| Time | | 2.03 (4, 243) | 0.03 [0, 1] | 0.09 |
| Session | | 0.34 (1, 243) | <.01 [0, 1] | 0.56 |
| Age | | 0.12 (1, 27) | <.01 [0, 1] | 0.73 |
| Sex | | 1.5 (1, 27) | 0.05 [0, 1] | 0.23 |
| Time: session | | 0.73 (4, 243) | 0.01 [0, 1] | 0.57 |
| Time: age | | 0.79 (4, 243) | 0.01 [0, 1] | 0.53 |
| Time: sex | | 1.39 (4, 243) | 0.02 [0, 1] | 0.24 |
| Session: age | | 5.73 (1, 243) | 0.02 [0, 1] | 0.02 |
| Session: sex | | 0.02 (1, 243) | <.01 [0, 1] | 0.89 |
| Time: session: age | | 0.34 (4, 243) | <.01 [0, 1] | 0.85 |
| Time: session: sex | | 0.47 (4, 243) | <.01 [0, 1] | 0.76 |
| **Between session comparisons: Session 2 – Session 1** | | | | |
| Arm | Minutes post-HFS | Estimate [95% CI] | Std. effect [95% CI] | *P* |
| Control | -10 | -0.2 [-0.61, 0.21] | -0.31 [-0.94, 0.32] | 0.33 |
| Control | 5 | -0.15 [-0.56, 0.26] | -0.23 [-0.86, 0.4] | 0.48 |
| Control | 20 | -0.04 [-0.45, 0.37] | -0.07 [-0.7, 0.56] | 0.84 |
| Control | 35 | 0.23 [-0.18, 0.64] | 0.35 [-0.28, 0.98] | 0.28 |
| Control | 50 | -0.09 [-0.5, 0.32] | -0.14 [-0.77, 0.49] | 0.67 |
| **Session 1 comparisons to baseline: Post-HFS – Baseline** | | | | |
| Arm | Minutes post-HFS | Estimate [95% CI] | Std. effect [95% CI] | *P* |
| Control | 5 | 0.01 [-0.51, 0.53] | 0.02 [-0.62, 0.65] | 1 |
| Control | 20 | -0.34 [-0.85, 0.18] | -0.52 [-1.15, 0.11] | 0.31 |
| Control | 35 | -0.41 [-0.92, 0.11] | -0.62 [-1.26, 0.01] | 0.17 |
| Control | 50 | -0.29 [-0.81, 0.22] | -0.45 [-1.09, 0.18] | 0.42 |
| **Session 2 comparisons to baseline: Post-HFS – Baseline** | | | | |
| Arm | Minutes post-HFS | Estimate [95% CI] | Std. effect [95% CI] | *P* |
| Control | 5 | 0.06 [-0.45, 0.58] | 0.1 [-0.53, 0.73] | 0.98 |
| Control | 20 | -0.18 [-0.69, 0.34] | -0.27 [-0.91, 0.36] | 0.77 |
| Control | 35 | 0.02 [-0.49, 0.54] | 0.03 [-0.6, 0.67] | 1 |
| Control | 50 | -0.18 [-0.7, 0.33] | -0.28 [-0.91, 0.35] | 0.76 |

Supplementary Table 2 continued

**Pinprick-evoked α-ERD magnitude results**

| | | | |
|---|---|---|---|
| **c) Model assessment: Magnitude influence on pain ratings** | | | |
| Pain rating ~ time * session * arm * (magnitude change + magnitude baseline) + (1 \| subject) | | | |
| Conditional $R^2$ = .80 | Marginal $R^2$ = .07 | | |
| **Fixed effects and interactions** | | | |
| Predictor | *F* (DF) | *η2* [95% CI] | *P* |
| Time | 7.76 (1, 513.18) | 0.01 [0, 1] | 0.006 |
| Session | 59.28 (1, 515.47) | 0.1 [0.07, 1] | <.001 |
| Arm | 42.61 (1, 513.23) | 0.08 [0.04, 1] | <.001 |
| Magnitude change | 0.88 (1, 518.52) | <.01 [0, 1] | 0.35 |
| Magnitude baseline | 0.02 (1, 534.74) | <.01 [0, 1] | 0.88 |
| Time: session | 0.72 (1, 513.06) | <.01 [0, 1] | 0.4 |
| Time: arm | 13.99 (1, 513) | 0.03 [0.01, 1] | <.001 |
| Session: arm | 0.63 (1, 513.11) | <.01 [0, 1] | 0.42 |
| Time: magnitude change | 1.97 (1, 513.41) | <.01 [0, 1] | 0.16 |
| Time: magnitude baseline | 4.89 (1, 513.03) | <.01 [0, 1] | 0.03 |
| Session: magnitude change | 1.19 (1, 514.82) | <.01 [0, 1] | 0.28 |
| Session: magnitude baseline | 2.74 (1, 519.25) | <.01 [0, 1] | 0.1 |
| Arm: magnitude change | 1.57 (1, 513.72) | <.01 [0, 1] | 0.21 |
| Arm: magnitude baseline | 2.39 (1, 515.39) | <.01 [0, 1] | 0.12 |
| Time: session: arm | 0 (1, 513.01) | <.01 [0, 1] | 0.99 |
| Time: session: magnitude change | 0.72 (1, 513.33) | <.01 [0, 1] | 0.4 |
| Time: session: magnitude baseline | 0 (1, 513.01) | <.01 [0, 1] | 0.96 |
| Time: arm: magnitude change | 0.46 (1, 513.48) | <.01 [0, 1] | 0.5 |
| Time: arm: magnitude baseline | 1.14 (1, 513.02) | <.01 [0, 1] | 0.29 |
| Session: arm: magnitude change | 2.35 (1, 514.46) | <.01 [0, 1] | 0.13 |
| Session: arm: magnitude baseline | 0.69 (1, 515.16) | <.01 [0, 1] | 0.41 |
| Time: session: arm: magnitude change | 2.14 (1, 513.58) | <.01 [0, 1] | 0.14 |
| Time: session: arm: magnitude baseline | 0.12 (1, 513.05) | <.01 [0, 1] | 0.73 |

*Supplementary Table 2 Pinprick-evoked α-ERD mixed-effects modelling*

The linear mixed effects modelling results for α-ERD data are displayed for the a) HFS arm, b) control arm, and c) their relationship with MPS. HFS = High-frequency stimulation. α-ERD = alpha-band event-related desynchronisation. N = 27, (female = 19).

Supplementary Table 3

**Pinprick-evoked θ-ERS magnitude results**

| **a) Model assessment: HFS arm** | | | | |
|---|---|---|---|---|
| Magnitude ~ time * session * (age + sex) + (1 \| subject) | | | | |
| Conditional $R^2$ = .35 | | .09 | | |
| **Fixed effects and interactions** | | | | |
| Predictor | | *F* (DF) | *η2* [95% CI] | *p* |
| Time | | 0.59 (4, 243) | <.01 [0, 1] | 0.67 |
| Session | | 0.27 (1, 243) | <.01 [0, 1] | 0.61 |
| Age | | 0.06 (1, 27) | <.01 [0, 1] | 0.80 |
| Sex | | 0.53 (1, 27) | 0.02 [0, 1] | 0.47 |
| Time: session | | 1.91 (4, 243) | 0.03 [0, 1] | 0.11 |
| Time: age | | 0.69 (4, 243) | 0.01 [0, 1] | 0.60 |
| Time: sex | | 2.63 (4, 243) | 0.04 [0, 1] | 0.04 |
| Session: age | | 0.63 (1, 243) | <.01 [0, 1] | 0.43 |
| Session: sex | | 1.88 (1, 243) | <.01 [0, 1] | 0.17 |
| Time: session: age | | 0.53 (4, 243) | <.01 [0, 1] | 0.71 |
| Time: session: sex | | 1.63 (4, 243) | 0.03 [0, 1] | 0.17 |
| **Between session comparisons: Session 2 – Session 1** | | | | |
| Arm | Minutes post-HFS | Estimate [95% CI] | Std. effect [95% CI] | *P* |
| HFS | -10 | -0.36 [-0.86, 0.14] | -0.45 [-1.07, 0.18] | 0.162 |
| HFS | 5 | -0.11 [-0.61, 0.39] | -0.14 [-0.76, 0.49] | 0.669 |
| HFS | 20 | -0.04 [-0.54, 0.47] | -0.05 [-0.67, 0.58] | 0.886 |
| HFS | 35 | 0.32 [-0.18, 0.82] | 0.4 [-0.23, 1.03] | 0.208 |
| HFS | 50 | 0.46 [-0.04, 0.96] | 0.57 [-0.06, 1.2] | 0.074 |
| **Session 1 comparisons to baseline: Post-HFS – Baseline** | | | | |
| Arm | Minutes post-HFS | Estimate [95% CI] | Std. effect [95% CI] | P |
| HFS | 5 | -0.18 [-0.81, 0.45] | -0.22 [-0.85, 0.4] | 0.848 |
| HFS | 20 | -0.25 [-0.88, 0.38] | -0.32 [-0.94, 0.31] | 0.685 |
| HFS | 35 | -0.47 [-1.1, 0.16] | -0.59 [-1.22, 0.04] | 0.201 |
| HFS | 50 | -0.65 [-1.28, -0.02] | -0.81 [-1.44, -0.18] | 0.04 |
| **Session 2 comparisons to baseline: Post-HFS – Baseline** | | | | |
| Arm | Minutes post-HFS | Estimate [95% CI] | Std. effect [95% CI] | *P* |
| HFS | 5 | 0.07 [-0.56, 0.7] | 0.09 [-0.54, 0.71] | 0.984 |
| HFS | 20 | 0.07 [-0.56, 0.7] | 0.08 [-0.54, 0.71] | 0.985 |
| HFS | 35 | 0.21 [-0.42, 0.84] | 0.26 [-0.37, 0.88] | 0.794 |
| HFS | 50 | 0.16 [-0.47, 0.79] | 0.2 [-0.42, 0.83] | 0.878 |

Supplementary Table 3 continued

**Pinprick-evoked θ-ERS magnitude results**

| **b) Model assessment: Control arm** | | | | |
|---|---|---|---|---|
| Magnitude ~ time * session * (age + sex) + (1 \| subject) | | | | |
| Conditional $R^2$ = .39 | | Marginal $R^2$ = .12 | | |
| **Fixed effects and interactions** | | | | |
| Predictor | | *F* (DF) | *η2* [95% CI] | *P* |
| Time | | 0.44 (4, 243) | <.01 [0, 1] | 0.78 |
| Session | | 1.61 (1, 243) | <.01 [0, 1] | 0.20 |
| Age | | 0 (1, 27) | <.01 [0, 1] | 0.99 |
| Sex | | 3.9 (1, 27) | 0.13 [0, 1] | 0.06 |
| Time: session | | 1.96 (4, 243) | 0.03 [0, 1] | 0.10 |
| Time: age | | 2.03 (4, 243) | 0.03 [0, 1] | 0.09 |
| Time: sex | | 0.93 (4, 243) | 0.02 [0, 1] | 0.45 |
| Session: age | | 0.14 (1, 243) | <.01 [0, 1] | 0.71 |
| Session: sex | | 0.28 (1, 243) | <.01 [0, 1] | 0.60 |
| Time: session: age | | 0.91 (4, 243) | 0.01 [0, 1] | 0.46 |
| Time: session: sex | | 1.08 (4, 243) | 0.02 [0, 1] | 0.36 |
| **Between session comparisons: Session 2 – Session 1** | | | | |
| Arm | Minutes post-HFS | Estimate [95% CI] | Std. effect [95% CI] | *P* |
| Control | -10 | -0.4 [-0.89, 0.09] | -0.52 [-1.14, 0.11] | 0.106 |
| Control | 5 | -0.4 [-0.88, 0.09] | -0.51 [-1.14, 0.12] | 0.111 |
| Control | 20 | -0.29 [-0.78, 0.19] | -0.38 [-1.01, 0.25] | 0.233 |
| Control | 35 | 0.3 [-0.19, 0.78] | 0.38 [-0.24, 1.01] | 0.230 |
| Control | 50 | 0.13 [-0.35, 0.62] | 0.17 [-0.45, 0.8] | 0.588 |
| **Session 1 comparisons to baseline: Post-HFS – Baseline** | | | | |
| Arm | Minutes post-HFS | Estimate [95% CI] | Std. effect [95% CI] | *P* |
| Control | 5 | -0.03 [-0.64, 0.58] | -0.04 [-0.67, 0.58] | 0.997 |
| Control | 20 | -0.02 [-0.63, 0.59] | -0.02 [-0.65, 0.61] | 0.999 |
| Control | 35 | -0.44 [-1.05, 0.17] | -0.57 [-1.19, 0.06] | 0.232 |
| Control | 50 | -0.15 [-0.76, 0.46] | -0.2 [-0.82, 0.43] | 0.889 |
| **Session 2 comparisons to baseline: Post-HFS – Baseline** | | | | |
| Arm | Minutes post-HFS | Estimate [95% CI] | Std. effect [95% CI] | *P* |
| Control | 5 | -0.03 [-0.64, 0.58] | -0.03 [-0.66, 0.59] | 0.998 |
| Control | 20 | 0.09 [-0.52, 0.7] | 0.12 [-0.51, 0.74] | 0.967 |
| Control | 35 | 0.26 [-0.35, 0.87] | 0.33 [-0.29, 0.96] | 0.653 |
| Control | 50 | 0.38 [-0.23, 0.99] | 0.49 [-0.14, 1.12] | 0.344 |

Supplementary Table 3 continued

**Pinprick-evoked θ-ERS magnitude results**

| **c) Model assessment: Magnitude influence on pain ratings** | | | |
|---|---|---|---|
| Pain rating ~ time * session * arm * (magnitude change + magnitude baseline) + (1 \| subject) | | | |
| Conditional $R^2$ = .78 | Marginal $R^2$ = .07 | | |
| **Fixed effects and interactions** | | | |
| Predictor | *F* (DF) | *η2* [95% CI] | *P* |
| Time | 7.49 (1, 512.98) | 0.01 [0, 1] | 0.006 |
| Session | 67.89 (1, 515.32) | 0.12 [0.08, 1] | <.001 |
| Arm | 45.65 (1, 513.55) | 0.08 [0.05, 1] | <.001 |
| Magnitude change | 0.1 (1, 517.3) | <.01 [0, 1] | 0.758 |
| Magnitude baseline | 1.42 (1, 534.97) | <.01 [0, 1] | 0.234 |
| Time: session | 0.31 (1, 513.08) | <.01 [0, 1] | 0.579 |
| Time: arm | 15.11 (1, 512.98) | 0.03 [0.01, 1] | <.001 |
| Session: arm | 1.24 (1, 513.41) | <.01 [0, 1] | 0.266 |
| Time: magnitude change | 0.95 (1, 513.57) | <.01 [0, 1] | 0.331 |
| Time: magnitude baseline | 1.92 (1, 513.3) | <.01 [0, 1] | 0.166 |
| Session: magnitude change | 5.4 (1, 513.67) | 0.01 [0, 1] | 0.021 |
| Session: magnitude baseline | 4.05 (1, 514.89) | <.01 [0, 1] | 0.045 |
| Arm: magnitude change | 0.7 (1, 514.92) | <.01 [0, 1] | 0.402 |
| Arm: magnitude baseline | 1.05 (1, 515.66) | <.01 [0, 1] | 0.307 |
| Time: session: arm | 0.45 (1, 513) | <.01 [0, 1] | 0.504 |
| Time: session: magnitude change | 0.41 (1, 513.63) | <.01 [0, 1] | 0.524 |
| Time: session: magnitude baseline | 0.46 (1, 513.07) | <.01 [0, 1] | 0.496 |
| Time: arm: magnitude change | 0.07 (1, 513.69) | <.01 [0, 1] | 0.787 |
| Time: arm: magnitude baseline | 1.06 (1, 513.23) | <.01 [0, 1] | 0.305 |
| Session: arm: magnitude change | 0.26 (1, 514.02) | <.01 [0, 1] | 0.611 |
| Session: arm: magnitude baseline | 2.37 (1, 514.36) | <.01 [0, 1] | 0.124 |
| Time: session: arm: magnitude change | 0.7 (1, 513.87) | <.01 [0, 1] | 0.404 |
| Time: session: arm: magnitude baseline | 0.26 (1, 513.08) | <.01 [0, 1] | 0.609 |

*Supplementary Table 3 Pinprick-evoked θ-ERS mixed-effects modelling*

The linear mixed effects modelling results for θ-ERS data are displayed for the a) HFS arm, b) control arm, and c) their relationship with MPS. HFS = High-frequency stimulation. θ-ERS = threta-band event-related synchronisation. N = 27, (female = 19).

Supplementary Table 4

**Pinprick-evoked VP magnitude results**

| **a) Model assessment: HFS arm** | | | | |
|---|---|---|---|---|
| Magnitude ~ time * session * (age + sex) + (1 \| subject) | | | | |
| Conditional $R^2$ = .52 | | Marginal $R^2$ = .09 | | |
| **Fixed effects and interactions** | | | | |
| Predictor | | *F* (DF) | *η2* [95% CI] | *p* |
| Time | | 0.16 (4, 243) | <.01 [0, 1] | 0.958 |
| Session | | 25.29 (1, 243) | 0.09 [0.04, 1] | <.001 |
| Age | | 0 (1, 27) | <.01 [0, 1] | 0.975 |
| Sex | | 0.99 (1, 27) | 0.04 [0, 1] | 0.33 |
| Time: session | | 0.68 (4, 243) | 0.01 [0, 1] | 0.608 |
| Time: age | | 0.71 (4, 243) | 0.01 [0, 1] | 0.585 |
| Time: sex | | 0.92 (4, 243) | 0.01 [0, 1] | 0.453 |
| Session: age | | 0.07 (1, 243) | <.01 [0, 1] | 0.786 |
| Session: sex | | 1.82 (1, 243) | <.01 [0, 1] | 0.178 |
| Time: session: age | | 0.18 (4, 243) | <.01 [0, 1] | 0.95 |
| Time: session: sex | | (4, 243) | 0.02 [0, 1] | 0.406 |
| **Between session comparisons: Session 2 – Session 1** | | | | |
| Arm | Minutes post-HFS | Estimate [95% CI] | Std. effect [95% CI] | *P* |
| HFS | -10 | -1.68 [-3.01, -0.35] | -0.79 [-1.43, -0.16] | 0.014 |
| HFS | 5 | -1.8 [-3.13, -0.47] | -0.85 [-1.48, -0.22] | 0.009 |
| HFS | 20 | -0.57 [-1.9, 0.76] | -0.27 [-0.9, 0.36] | 0.400 |
| HFS | 35 | -1.78 [-3.11, -0.45] | -0.84 [-1.47, -0.21] | 0.009 |
| HFS | 50 | -1.29 [-2.62, 0.04] | -0.61 [-1.24, 0.02] | 0.057 |
| **Session 1 comparisons to baseline: Post-HFS – Baseline** | | | | |
| Arm | Minutes post-HFS | Estimate [95% CI] | Std. effect [95% CI] | P |
| HFS | 5 | 0.08 [-1.59, 1.75] | 0.04 [-0.59, 0.67] | 0.998 |
| HFS | 20 | -0.59 [-2.26, 1.08] | -0.28 [-0.91, 0.35] | 0.758 |
| HFS | 35 | 0 [-1.67, 1.67] | 0 [-0.63, 0.63] | 1.000 |
| HFS | 50 | -0.49 [-2.16, 1.19] | -0.23 [-0.86, 0.4] | 0.840 |
| **Session 2 comparisons to baseline: Post-HFS – Baseline** | | | | |
| Arm | Minutes post-HFS | Estimate [95% CI] | Std. effect [95% CI] | *P* |
| HFS | 5 | -0.04 [-1.72, 1.63] | -0.02 [-0.65, 0.61] | 0.999 |
| HFS | 20 | 0.52 [-1.15, 2.19] | 0.25 [-0.38, 0.88] | 0.815 |
| HFS | 35 | -0.1 [-1.77, 1.57] | -0.05 [-0.68, 0.58] | 0.996 |
| HFS | 50 | -0.1 [-1.77, 1.57] | -0.05 [-0.68, 0.58] | 0.996 |

Supplementary Table 4 continued

**Pinprick-evoked VP magnitude results**

| **b) Model assessment: Control arm** | | | | |
|---|---|---|---|---|
| Magnitude ~ time * session * (age + sex) + (1 \| subject) | | | | |
| Conditional $R^2$ = .52 | | Marginal $R^2$ = .12 | | |
| **Fixed effects and interactions** | | | | |
| Predictor | | *F* (DF) | *η2* [95% CI] | *P* |
| Time | | 1.14 (4, 243) | 0.02 [0, 1] | 0.338 |
| Session | | 14.96 (1, 243) | 0.06 [0.02, 1] | <.001 |
| Age | | 0.42 (1, 27) | 0.02 [0, 1] | 0.521 |
| Sex | | 1.75 (1, 27) | 0.06 [0, 1] | 0.198 |
| Time: session | | 1.24 (4, 243) | 0.02 [0, 1] | 0.295 |
| Time: age | | 1.12 (4, 243) | 0.02 [0, 1] | 0.349 |
| Time: sex | | 0.84 (4, 243) | 0.01 [0, 1] | 0.502 |
| Session: age | | 6.03 (1, 243) | 0.02 [0, 1] | 0.015 |
| Session: sex | | 1.75 (1, 243) | <.01 [0, 1] | 0.187 |
| Time: session: age | | 0.63 (4, 243) | 0.01 [0, 1] | 0.641 |
| Time: session: sex | | 0.08 (4, 243) | <.01 [0, 1] | 0.989 |
| **Between session comparisons: Session 2 – Session 1** | | | | |
| Arm | Minutes post-HFS | Estimate [95% CI] | Std. effect [95% CI] | *P* |
| Control | -10 | -1.13 [-2.17, -0.09] | -0.68 [-1.32, -0.05] | 0.033 |
| Control | 5 | -1.24 [-2.28, -0.2] | -0.75 [-1.38, -0.11] | 0.020 |
| Control | 20 | -1.14 [-2.18, -0.1] | -0.69 [-1.32, -0.05] | 0.033 |
| Control | 35 | -0.87 [-1.91, 0.17] | -0.53 [-1.16, 0.1] | 0.099 |
| Control | 50 | 0.1 [-0.94, 1.14] | 0.06 [-0.57, 0.69] | 0.850 |
| **Session 1 comparisons to baseline: Post-HFS – Baseline** | | | | |
| Arm | Minutes post-HFS | Estimate [95% CI] | Std. effect [95% CI] | *P* |
| Control | 5 | -0.25 [-1.56, 1.06] | -0.15 [-0.78, 0.48] | 0.939 |
| Control | 20 | -0.14 [-1.45, 1.17] | -0.09 [-0.71, 0.54] | 0.984 |
| Control | 35 | -0.84 [-2.15, 0.47] | -0.51 [-1.14, 0.12] | 0.323 |
| Control | 50 | -0.89 [-2.2, 0.42] | -0.54 [-1.17, 0.09] | 0.275 |
| **Session 2 comparisons to baseline: Post-HFS – Baseline** | | | | |
| Arm | Minutes post-HFS | Estimate [95% CI] | Std. effect [95% CI] | *P* |
| Control | 5 | -0.35 [-1.66, 0.96] | -0.21 [-0.84, 0.42] | 0.865 |
| Control | 20 | -0.14 [-1.45, 1.16] | -0.09 [-0.72, 0.54] | 0.983 |
| Control | 35 | -0.58 [-1.88, 0.73] | -0.35 [-0.98, 0.28] | 0.624 |
| Control | 50 | 0.34 [-0.96, 1.65] | 0.21 [-0.42, 0.84] | 0.872 |

Supplementary Table 4 continued

**Pinprick-evoked VP magnitude results**

| **c) Model assessment: Magnitude influence on pain ratings** | | | |
|---|---|---|---|
| Pain rating ~ time * session * arm * (magnitude change + magnitude baseline) + (1 \| subject) | | | |
| Conditional $R^2$ = .78 | Marginal $R^2$ = .07 | | |
| **Fixed effects and interactions** | | | |
| Predictor | *F* (DF) | *η2* [95% CI] | *P* |
| Time | 9.92 (1, 513) | 0.02 [0, 1] | 0.002 |
| Session | 54.21 (1, 516) | 0.1 [0.06, 1] | <.001 |
| Arm | 36.48 (1, 513) | 0.07 [0.04, 1] | <.001 |
| Magnitude change | 4.25 (1, 519) | <.01 [0, 1] | 0.04 |
| Magnitude baseline | 2.63 (1, 533) | <.01 [0, 1] | 0.106 |
| Time: session | 1.32 (1, 513) | <.01 [0, 1] | 0.251 |
| Time: arm | 14.21 (1, 513) | 0.03 [0.01, 1] | <.001 |
| Session: arm | 0.58 (1, 513) | <.01 [0, 1] | 0.446 |
| Time: magnitude change | 0.04 (1, 514) | <.01 [0, 1] | 0.84 |
| Time: magnitude baseline | 0.02 (1, 514) | <.01 [0, 1] | 0.884 |
| Session: magnitude change | 0.75 (1, 514) | <.01 [0, 1] | 0.385 |
| Session: magnitude baseline | 0.61 (1, 517) | <.01 [0, 1] | 0.435 |
| Arm: magnitude change | 0.79 (1, 514) | <.01 [0, 1] | 0.373 |
| Arm: magnitude baseline | 0.15 (1, 516) | <.01 [0, 1] | 0.697 |
| Time: session: arm | 0.62 (1, 513) | <.01 [0, 1] | 0.431 |
| Time: session: magnitude change | 0.32 (1, 514) | <.01 [0, 1] | 0.574 |
| Time: session: magnitude baseline | 0.53 (1, 513) | <.01 [0, 1] | 0.466 |
| Time: arm: magnitude change | 2.43 (1, 513) | <.01 [0, 1] | 0.12 |
| Time: arm: magnitude baseline | 0.13 (1, 513) | <.01 [0, 1] | 0.716 |
| Session: arm: magnitude change | 0.19 (1, 514) | <.01 [0, 1] | 0.664 |
| Session: arm: magnitude baseline | 0.84 (1, 514) | <.01 [0, 1] | 0.359 |
| Time: session: arm: magnitude change | 0.24 (1, 513) | <.01 [0, 1] | 0.621 |
| Time: session: arm: magnitude baseline | 0.46 (1, 513) | <.01 [0, 1] | 0.497 |

*Supplementary Table 4 Pinprick-evoked VP magnitude mixed-effects modelling*

The linear mixed effects modelling results for VP magnitude data are displayed for the a) HFS arm, b) control arm, and c) their relationship with MPS. HFS = High-frequency stimulation. VP = Vertex potential. N = 27, (female = 19).

Supplementary Table 5

**Pinprick-evoked VP latency results**

| a) Model assessment: HFS arm | | | | |
|---|---|---|---|---|
| Latency ~ time * session * (age + sex) + (1 \| subject) | | | | |
| Conditional $R^2$ = .34 | | Marginal $R^2$ = .13 | | |
| **Fixed effects and interactions** | | | | |
| Predictor | | *F* (DF) | *η2* [95% CI] | *p* |
| Time | | 0.55 (4, 243) | <.01 [0, 1] | .70 |
| Session | | 0.44 (1, 243) | <.01 [0, 1] | .51 |
| Age | | 3.15 (1, 27) | 0.1 [0, 1] | .09 |
| Sex | | 0.31 (1, 27) | 0.01 [0, 1] | .58 |
| Time: session | | 1.62 (4, 243) | 0.03 [0, 1] | .17 |
| Time: age | | 0.43 (4, 243) | <.01 [0, 1] | .79 |
| Time: sex | | 1.59 (4, 243) | 0.03 [0, 1] | .18 |
| Session: age | | 8.68 (1, 243) | 0.03 [0.01, 1] | .004 |
| Session: sex | | 0.38 (1, 243) | <.01 [0, 1] | .54 |
| Time: session: age | | 1.59 (4, 243) | 0.03 [0, 1] | .18 |
| Time: session: sex | | 0.69 (4, 243) | 0.01 [0, 1] | .60 |
| **Between session comparisons: Session 2 – Session 1** | | | | |
| Arm | Minutes post-HFS | Estimate [95% CI] | Std. effect [95% CI] | *P* |
| HFS | -10 | 0.01 [-0.49, 0.52] | 0.02 [-0.61, 0.64] | .954 |
| HFS | 5 | -0.09 [-0.59, 0.42] | -0.11 [-0.74, 0.51] | .728 |
| HFS | 20 | 0.46 [-0.05, 0.96] | 0.57 [-0.06, 1.2] | .074 |
| HFS | 35 | 0.29 [-0.22, 0.79] | 0.36 [-0.27, 0.98] | .262 |
| HFS | 50 | -0.32 [-0.82, 0.19] | -0.39 [-1.02, 0.23] | .217 |
| **Session 1 comparisons to baseline: Post-HFS – Baseline** | | | | |
| Arm | Minutes post-HFS | Estimate [95% CI] | Std. effect [95% CI] | P |
| HFS | 5 | 0.03 [-0.61, 0.66] | 0.03 [-0.59, 0.66] | .999 |
| HFS | 20 | -0.12 [-0.76, 0.51] | -0.15 [-0.78, 0.47] | .936 |
| HFS | 35 | -0.29 [-0.92, 0.34] | -0.36 [-0.98, 0.27] | .602 |
| HFS | 50 | 0.13 [-0.5, 0.76] | 0.16 [-0.47, 0.78] | .931 |
| **Session 2 comparisons to baseline: Post-HFS – Baseline** | | | | |
| Arm | Minutes post-HFS | Estimate [95% CI] | Std. effect [95% CI] | *P* |
| HFS | 5 | -0.08 [-0.71, 0.55] | -0.1 [-0.72, 0.53] | .978 |
| HFS | 20 | 0.32 [-0.31, 0.95] | 0.4 [-0.23, 1.02] | .523 |
| HFS | 35 | -0.02 [-0.65, 0.62] | -0.02 [-0.64, 0.61] | 1 |
| HFS | 50 | -0.2 [-0.84, 0.43] | -0.25 [-0.88, 0.37] | .803 |

Supplementary Table 5 continued

**Pinprick-evoked VP latency results**

| | | | | |
|---|---|---|---|---|
| **b) Model assessment: Control arm** | | | | |
| Latency ~ time * session * (age + sex) + (1 \| subject) | | | | |
| Conditional $R^2$ = .25 | | Marginal $R^2$ = .09 | | |
| **Fixed effects and interactions** | | | | |
| Predictor | | *F* (DF) | *η2* [95% CI] | *P* |
| Time | | 1.39 (4, 243) | 0.02 [0, 1] | .24 |
| Session | | 0.34 (1, 243) | <.01 [0, 1] | .56 |
| Age | | 0.61 (1, 27) | 0.02 [0, 1] | .44 |
| Sex | | 0.02 (1, 27) | <.01 [0, 1] | .88 |
| Time: session | | 1.27 (4, 243) | 0.02 [0, 1] | .28 |
| Time: age | | 0.83 (4, 243) | 0.01 [0, 1] | .50 |
| Time: sex | | 0.33 (4, 243) | <.01 [0, 1] | .86 |
| Session: age | | 2.3 (1, 243) | <.01 [0, 1] | .13 |
| Session: sex | | 0.02 (1, 243) | <.01 [0, 1] | .88 |
| Time: session: age | | 1.47 (4, 243) | 0.02 [0, 1] | .21 |
| Time: session: sex | | 1.45 (4, 243) | 0.02 [0, 1] | .22 |
| **Between session comparisons: Session 2 – Session 1** | | | | |
| Arm | Minutes post-HFS | Estimate [95% CI] | Std. effect [95% CI] | *P* |
| Control | -10 | 0.06 [-0.48, 0.6] | 0.07 [-0.55, 0.7] | 0.823 |
| Control | 5 | 0.38 [-0.15, 0.92] | 0.45 [-0.18, 1.07] | 0.161 |
| Control | 20 | -0.14 [-0.68, 0.4] | -0.16 [-0.79, 0.46] | 0.609 |
| Control | 35 | -0.29 [-0.83, 0.25] | -0.34 [-0.96, 0.29] | 0.288 |
| Control | 50 | 0.32 [-0.22, 0.86] | 0.37 [-0.25, 1] | 0.241 |
| **Session 1 comparisons to baseline: Post-HFS – Baseline** | | | | |
| Arm | Minutes post-HFS | Estimate [95% CI] | Std. effect [95% CI] | *P* |
| Control | 5 | -0.34 [-1.02, 0.33] | -0.4 [-1.03, 0.22] | 0.515 |
| Control | 20 | 0.32 [-0.35, 1] | 0.38 [-0.25, 1] | 0.564 |
| Control | 35 | 0.18 [-0.49, 0.85] | 0.21 [-0.42, 0.83] | 0.871 |
| Control | 50 | -0.22 [-0.9, 0.45] | -0.26 [-0.89, 0.37] | 0.790 |
| **Session 2 comparisons to baseline: Post-HFS – Baseline** | | | | |
| Arm | Minutes post-HFS | Estimate [95% CI] | Std. effect [95% CI] | *P* |
| Control | 5 | -0.02 [-0.69, 0.65] | -0.03 [-0.65, 0.6] | 0.999 |
| Control | 20 | 0.12 [-0.55, 0.8] | 0.14 [-0.48, 0.77] | 0.946 |
| Control | 35 | -0.17 [-0.84, 0.5] | -0.2 [-0.82, 0.43] | 0.883 |
| Control | 50 | 0.04 [-0.64, 0.71] | 0.04 [-0.58, 0.67] | 0.997 |

Supplementary Table 5 continued

**Pinprick-evoked VP latency results**

| **c) Model assessment: Latency influence on pain ratings** | | | |
|---|---|---|---|
| Pain rating ~ time * session * arm * (latency change + latency baseline) + (1 \| subject) | | | |
| Conditional $R^2$ = .78 | Marginal $R^2$ = .07 | | |
| **Fixed effects and interactions** | | | |
| Predictor | *F* (DF) | *η2* [95% CI] | *P* |
| Time | 7.46 (1, 513) | 0.01 [0, 1] | 0.007 |
| Session | 70.45 (1, 513) | 0.12 [0.08, 1] | <.001 |
| Arm | 47.06 (1, 513) | 0.08 [0.05, 1] | <.001 |
| Latency change | 1.67 (1, 516) | <.01 [0, 1] | 0.196 |
| Latency baseline | 2.33 (1, 522) | <.01 [0, 1] | 0.128 |
| Time: session | 0.85 (1, 513) | <.01 [0, 1] | 0.357 |
| Time: arm | 17.92 (1, 513) | 0.03 [0.01, 1] | <.001 |
| Session: arm | 0.49 (1, 513) | <.01 [0, 1] | 0.485 |
| Time: latency change | 2.38 (1, 513) | <.01 [0, 1] | 0.123 |
| Time: latency baseline | 0.07 (1, 513) | <.01 [0, 1] | 0.795 |
| Session: latency change | 0.73 (1, 514) | <.01 [0, 1] | 0.393 |
| Session: latency baseline | 1.17 (1, 518) | <.01 [0, 1] | 0.279 |
| Arm: latency change | 0.32 (1, 514) | <.01 [0, 1] | 0.573 |
| Arm: latency baseline | 1.31 (1, 516) | <.01 [0, 1] | 0.253 |
| Time: session: arm | 0.23 (1, 513) | <.01 [0, 1] | 0.628 |
| Time: session: latency change | 0.34 (1, 513) | <.01 [0, 1] | 0.559 |
| Time: session: latency baseline | 0.36 (1, 513) | <.01 [0, 1] | 0.55 |
| Time: arm: latency change | 0.53 (1, 513) | <.01 [0, 1] | 0.468 |
| Time: arm: latency baseline | 3.77 (1, 513) | <.01 [0, 1] | 0.053 |
| Session: arm: latency change | 3.38 (1, 514) | <.01 [0, 1] | 0.067 |
| Session: arm: latency baseline | 0.02 (1, 515) | <.01 [0, 1] | 0.902 |
| Time: session: arm: latency change | 0.04 (1, 513) | <.01 [0, 1] | 0.838 |
| Time: session: arm: latency baseline | 0.66 (1, 513) | <.01 [0, 1] | 0.417 |

*Supplementary Table 5 Pinprick-evoked VP latency mixed-effects modelling*

The linear mixed effects modelling results for VP latency data are displayed for the a) HFS arm, b) control arm, and c) their relationship with MPS. HFS = High-frequency stimulation. VP = Vertex potential. N = 27, (female = 19).

Supplementary Table 6

**Electrical-evoked EEG descriptive statistics and assessments of reliability and agreement**

| **Key:** | Value (standard deviation) or [95% confidence interval] | | | | |
|---|---|---|---|---|---|
| | **Raw data** | **Change from baseline (minutes)** | | | |
| **Time post-HFS** | Baseline | 5 | 20 | 35 | 50 |
| **Vertex potential magnitude (μV)** | | | | | |
| **HFS arm** | | | | | |
| Session 1 mean | 14.6 (4.6) | 1.4 (3.1) | .3 (2.8) | -.2 (3.6) | -.6 (3.3) |
| Session 1 range | 7.6 – 25 | -6.3 – 8 | -5.2 – 8 | -6.3 – 9 | -7 – 6 |
| Session 2 mean | 13.3 (4.6) | .8 (2.9) | .4 (2.4) | -1 (2.8) | -1.6 (3.1) |
| Session 2 range | 4.2 – 22 | -4.1 – 8 | -3.4 – 5 | -6.7 – 4 | -9.9 – 4 |
| ICC | .53 [.2 .8] | .32 [-.1 .6] | .13 [-.3 .5] | .23 [-.2 .6] | .39 [0 .7] |
| Bias | 1.2 [-.6 3] | .6 [-.8 2] | -.07 [-1.4 1.3] | .8 [-.8 2.4] | .9 [-.5 2.3] |
| Upper | 9.9 [6.9 13] | 7.5 [5.1 9.9] | 6.7 [4.3 9.1] | 8.6 [5.9 11.4] | 7.8 [5.4 10.2] |
| Lower | 7.5 [-10.5 -4.5] | -6.3 [-8.7 -3.9] | -6.8 [-9.2 -4.5] | 7.1 [-9.9 -4.4] | -6 [-8.4 -3.6] |
| **Control arm** | | | | | |
| Session 1 mean | 14.5 (4.8) | -1 (2.8) | -.7 (2.6) | -1.9 (2.8) | -.9 (2.9) |
| Session 1 range | 7.6 – 25 | -10.5 – 4 | -6.7 – 3 | -9 – 2 | -7.6 – 3 |
| Session 2 mean | 13 (5.1) | -.2 (2.6) | -.9 (1.8) | -1.6 (3.6) | -1.5 (3.9) |
| Session 2 range | 1.5 – 25 | -6.4 – 4 | -3.9 – 2 | -12.4 – 4 | -11.1 – 4 |
| ICC | .53 [.2 .8] | .36 [0 .6] | .16 [-.2 .5] | 0 [-.4 .4] | .39 [0 .7] |
| Bias | 1.5 [-.1 3.1] | -.9 [-2.1 .4] | .2 [-1 1.4] | -.3 [-2.1 1.6] | .6 [-1 2.1] |
| Upper | 9.3 [6.6 12] | 5.2 [3.1 7.2] | 6 [3.9 8] | 8.8 [5.7 12] | 8.1 [5.5 10.8] |
| Lower | -6.3 [-9 -3.6] | -6.9 [-8.9 -4.8] | -5.6 [-7.6 -3.6] | -9.3 [-12.5 -6.2] | -7 [-9.6 -4.3] |

Supplementary Table 6 continued

**Electrical-evoked EEG descriptive statistics and assessments of reliability and agreement**

| **Vertex potential latency** (ms) | | | | | |
|---|---|---|---|---|---|
| **HFS arm** | | | | | |
| Session 1 mean | 230.1 (46) | 2.2 (45) | -5 (33) | -2.9 (43) | -15.8 (43) |
| Session 1 range | 80 – 308 | -72 – 188 | -94 – 76 | -82 – 150 | -122 – 100 |
| Session 2 mean | 232 (49.1) | -.9 (34) | -10.1 (36) | -8.2 (41) | -14.7 (42) |
| Session 2 range | 174 – 388 | -88 – 78 | -106 – 78 | -108 – 104 | -106 – 58 |
| ICC | .51 [.2 .7] | .46 [.1 .7] | .06 [-.3 .4] | .47 [.1 .7] | .16 [-.2 .5] |
| Bias | -1.9 [-21 17] | 3.1 [-13 19] | 5.1 [-13 23] | 5.3 [-11 22] | -1.1 [-22 20] |
| Upper | 94.4 [60 128] | 85.3 [56 114] | 94.8 [63 126] | 88 [59 116] | 107 [69 144] |
| Lower | -98.1 [-131 -64] | -79 [-107 -50] | -84.5 [-115 -53] | -77.3 [-106 -48] | -109.2 [-146 -71] |
| **Control arm** | | | | | |
| Session 1 mean | 243.3 (62) | -2.1 (45) | -26.4 (70) | -32.7 (62) | -27.5 (51) |
| Session 1 range | 172 – 500 | -154 – 112 | -334 – 44 | -288 – 46 | -204 – 30 |
| Session 2 mean | 226.7 (58) | -9.6 (55) | -6.7 (34) | -14.5 (44) | -10.4 (50) |
| Session 2 range | 146 – 428 | -208 – 118 | -108 – 48 | -196 – 32 | -202 – 64 |
| ICC | .4 [0 .7] | .52 [.2 .8] | .26 [-.1 .6] | .2 [-.2 .5] | .2 [-.2 .5] |
| Bias | 16.6 [-3 36] | 7.6 [-9 24] | -19.7 [-42 2] | -18.2 [-38 2] | -17.1 [-35 1] |
| Upper | 113.7 [79 147] | 92.2 [62 121] | 91.6 [52 130] | 81.7 [46 116] | 72.2 [41 103] |
| Lower | -80.5 [-114 -46] | -77 [-106 -47] | -131 [-169 -92] | -118 [-152 -83] | -106.5 [-137 -75] |

Supplementary Table 6 continued

**Electrical-evoked EEG descriptive statistics and assessments of reliability and agreement**

| **α-ERD magnitude** (μV²/Hz) | | | | | |
|---|---|---|---|---|---|
| **HFS arm** | | | | | |
| Session 1 mean | -1.8 (2.1) | -.6 (1.4) | -1 (1.6) | -.7 (1.8) | -1.4 (2.7) |
| Session 1 range | -8.4 – -.1 | -3.6 – 3 | -4.4 – 3 | -4.9 – 3 | -10.5 – 4 |
| Session 2 mean | -2 (2) | -.9 (2.1) | -.9 (2.1) | -1.2 (1.5) | -.6 (2) |
| Session 2 range | -7.9 – -.1 | -9.9 – 1 | -8.4 – 1 | -4.4 – 2 | -7.7 – 4 |
| ICC | .61 [.3 .8] | 0 [-.4 .4] | 0 [-.4 .4] | 0 [-.4 .4] | 0 [-.4 .4] |
| Bias | .2 [-.5 1] | .3 [-.7 1.3] | -.2 [-1.3 1] | .5 [-.5 1.5] | -.8 [-2.1 .6] |
| Upper | 3.9 [2.6 5.1] | 5.4 [3.6 7.1] | 5.5 [3.5 7.5] | 5.5 [3.8 7.3] | 6 [3.6 8.3] |
| Lower | -3.4 [-4.6 -2.1] | -4.8 [-6.5 -3] | -5.9 [-7.9 -4] | -4.5 [-6.3 -2.8] | -7.5 [-9.9 -5.2] |
| **Control arm** | | | | | |
| Session 1 mean | -2.1 (1.7) | -.6 (1.9) | -.5 (2.4) | -.8 (1.8) | -.5 (1.6) |
| Session 1 range | -6.4 – -.1 | -6.9 – 2 | -7.8 – 4 | -6.3 – 1 | -5.9 – 2 |
| Session 2 mean | -2.4 (2.4) | 0 (1.8) | -.4 (2.4) | 0 (1.8) | -.7 (2.7) |
| Session 2 range | -11.3 – -.1 | -3.2 – 6 | -7.7 – 3 | -4.6 – 5 | -11.9 – 3 |
| ICC | .7 [.4 .9] | 0 [-.4 .4] | 0 [-.4 .4] | 0 [-.4 .4] | .06 [-.3 .4] |
| Bias | .4 [-.3 1] | -.5 [-1.6 .5] | -.2 [-1.7 1.4] | -.8 [-2.1 .5] | .2 [-1 1.4] |
| Upper | 3.6 [2.5 4.7] | 4.7 [2.9 6.5] | 7.4 [4.8 10.1] | 5.5 [3.3 7.7] | 6.2 [4.1 8.2] |
| Lower | -2.9 [-4 -1.7] | -5.7 [-7.6 -3.9] | -7.7 [-10.4 -5.1] | -7.1 [-9.3 -4.9] | -5.8 [-7.8 -3.7] |

Supplementary Table 6 continued

**Electrical-evoked EEG descriptive statistics and assessments of reliability and agreement**

| **θ-ERS magnitude** (μV²/Hz) | | | | | |
|---|---|---|---|---|---|
| **HFS arm** | | | | | |
| Session 1 mean | 2.4 (1.5) | -.1 (1.1) | -.4 (1.2) | -.7 (1.1) | -.6 (.9) |
| Session 1 range | .5 – 7 | -2.4 – 3 | -3.4 – 2 | -3.6 – 1 | -2.3 – 1 |
| Session 2 mean | 2.2 (1.1) | -0 (1.2) | -.2 (1) | -.3 (1) | -.7 (1.3) |
| Session 2 range | .8 – 5 | -2.1 – 3 | -2.4 – 3 | -1.7 – 3 | -3.6 – 3 |
| ICC | .45 [.1 .7] | .26 [-.1 .6] | .09 [-.3 .5] | 0 [-.4 .4] | 0 [-.4 .4] |
| Bias | .2 [-.4 .7] | -.02 [-.6 .5] | -.2 [-.8 .4] | -.4 [-1.1 .3] | .02 [-.7 .7] |
| Upper | 2.8 [1.9 3.8] | 2.8 [1.8 3.8] | 2.8 [1.8 3.9] | 3 [1.8 4.2] | 3.5 [2.3 4.7] |
| Lower | -2.5 [-3.4 -1.56] | -2.8 [-3.8 -1.8] | -3.2 [-4.3 -2.2] | -3.8 [-5 -2.6] | -3.4 [-4.6 -2.2] |
| **Control arm** | | | | | |
| Session 1 mean | 2.2 (1.5) | -.3 (.9) | -.4 (.9) | -.5 (.9) | -.7 (1.2) |
| Session 1 range | .2 – 7 | -2.6 – 2 | -2.2 – 1 | -2.2 – 1 | -3.7 – 2 |
| Session 2 mean | 1.9 (1.2) | -0 (1) | -.3 (1.1) | -.4 (1) | 0 (1.3) |
| Session 2 range | .4 – 5 | -1.7 – 3 | -3.3 – 2 | -3.1 – 1 | -2.7 – 3 |
| ICC | .48 [.1 .7] | 0 [-.4 .4] | 0 [-.4 .4] | 0 [-.4 .4] | 0 [-.4 .4] |
| Bias | .3 [-.2 .9] | -.3 [-.9 .3] | -.1 [-.7 .5] | -.2 [-.8 .5] | -.7 [-1.5 .1] |
| Upper | 3 [2.1 4] | 2.8 [1.7 3.8] | 2.8 [1.8 3.8] | 3 [1.9 4.1] | 3.1 [1.8 4.4] |
| Lower | -2.4 [-3.3 -1.4] | -3.4 [-4.4 -2.3] | -3.1 [-4.1 -2.1] | -3.3 [-4.4 -2.2] | -4.5 [-5.8 -3.1] |

Supplementary Table 6 continued

**Electrical-evoked EEG descriptive statistics and assessments of reliability and agreement**

| **γ-ERS magnitude** (μV$^2$/Hz) | | | | | |
|---|---|---|---|---|---|
| **HFS arm** | | | | | |
| Session 1 mean | .5<br>(.2) | -0<br>(.2) | -0<br>(.1) | -0<br>(.2) | -0<br>(.2) |
| Session 1 range | .3 - 1 | -.6 - 0 | -.4 - 0 | -.7 - 0 | -.6 - 0 |
| Session 2 mean | .4<br>(.1) | 0<br>(.1) | .1<br>(.2) | .1<br>(.2) | .1<br>(.1) |
| Session 2 range | .2 - 1 | -.2 - 0 | -.2 - 1 | -.1 - 1 | -.1 - 0 |
| ICC | .35<br>[-.03 .64] | 0<br>[-.37 .37] | 0<br>[-.37 .37] | .11<br>[-.28 .46] | 0<br>[-.37 .37] |
| Bias | .06<br>[0 .13] | -.06<br>[-.16 .03] | -.06<br>[-.15 .02] | -.1<br>[-.21 0] | -.09<br>[-.2 .01] |
| Upper | .39<br>[.28 .5] | .40<br>[.24 .56] | .35<br>[.21 .5] | .40<br>[.22 .57] | .42<br>[.24 .6] |
| Lower | -.26<br>[-.37 -.15] | -.52<br>[-.69 -.36] | -.48<br>[-.63 -.33] | -.61<br>[-.78 -.43] | -.6<br>[-.78 -.42] |
| **Control arm** | | | | | |
| Session 1 mean | .4<br>(.1) | 0<br>(.1) | 0<br>(.2) | 0<br>(.2) | 0<br>(.1) |
| Session 1 range | .3 - 1 | -.3 - 0 | -.3 - 1 | -.3 - 1 | -.4 - 0 |
| Session 2 mean | .4<br>(.1) | 0<br>(.1) | 0<br>(.2) | 0<br>(.2) | 0<br>(.2) |
| Session 2 range | 0.2 - 1 | -0.3 - 0 | -0.4 - 0 | -0.4 - 0 | -.3 - 0 |
| ICC | .28<br>[-.11 .59] | 0<br>[-.37 .37] | 0<br>[-.37 .37] | .1<br>[-.28 .46] | 0<br>[-0.37 0.37] |
| Bias | -.02<br>[-.09 .04] | 0<br>[-.06 .06] | .05<br>[-.07 .16] | .02<br>[-.08 .12] | .02<br>[-.08 .11] |
| Upper | .29<br>[.18 .39] | .31<br>[.20 .42] | .63<br>[.43 .84] | .52<br>[.34 .69] | .48<br>[.32 .64] |
| Lower | -0.34<br>[-.44 -.23] | -.32<br>[-.42 -.21] | -.54<br>[-.74 -.34] | -.47<br>[-.64 -.3] | -.45<br>[-.61 -.28] |

*Supplementary Table 6 Electrical-evoked EEG descriptive statistics and assessments of reliability and agreement*

Mean values, standard deviations, and ranges were calculated separately for each session, time point, arm, and stimulation modality. Change from baseline scores were calculated as the absolute difference from baseline by subtracting the baseline values for each EEG response from their post-HFS counterparts. ERD = event-related desynchronisation. ERS = event-related synchronisation. HFS = High-frequency stimulation. N = 27, (female = 19).

Supplementary Table 7

**Electrical-evoked α-ERD magnitude results**

| **a) Model assessment: HFS arm**<br>Magnitude ~ time * session * (age + sex) + (1 \| subject) | | | | |
|---|---|---|---|---|
| Conditional $R^2$ = .78 | | Marginal $R^2$ = .07 | | |
| **Fixed effects and interactions** | | | | |
| Predictor | | *F* (DF) | *η2* [95% CI] | *p* |
| Time | | 8.71 (4, 243) | 0.13 [0.06, 1] | <.001 |
| Session | | 2.82 (1, 243) | 0.01 [0, 1] | 0.094 |
| Age | | 0.04 (1, 27) | <.01 [0, 1] | 0.851 |
| Sex | | 0.41 (1, 27) | 0.01 [0, 1] | 0.527 |
| Time: session | | 1.94 (4, 243) | 0.03 [0, 1] | 0.104 |
| Time: age | | 0.51 (4, 243) | <.01 [0, 1] | 0.727 |
| Time: sex | | 1.87 (4, 243) | 0.03 [0, 1] | 0.116 |
| Session: age | | 1.22 (1, 243) | <.01 [0, 1] | 0.271 |
| Session: sex | | 0.36 (1, 243) | <.01 [0, 1] | 0.55 |
| Time: session: age | | 0.65 (4, 243) | 0.01 [0, 1] | 0.626 |
| Time: session: sex | | 0.35 (4, 243) | <.01 [0, 1] | 0.841 |
| **Between session comparisons: Session 2 – Session 1** | | | | |
| Arm | Minutes post-HFS | Estimate [95% CI] | Std. effect [95% CI] | *P* |
| HFS | -10 | -0.22 [-0.52, 0.08] | -0.46 [-1.1, 0.18] | 0.152 |
| HFS | 5 | -0.15 [-0.45, 0.15] | -0.32 [-0.96, 0.32] | 0.319 |
| HFS | 20 | 0.14 [-0.16, 0.44] | 0.3 [-0.34, 0.93] | 0.354 |
| HFS | 35 | -0.34 [-0.64, -0.04] | -0.73 [-1.37, -0.09] | 0.025 |
| HFS | 50 | 0.04 [-0.26, 0.34] | 0.09 [-0.54, 0.73] | 0.772 |
| **Session 1 comparisons to baseline: Post-HFS – Baseline** | | | | |
| Arm | Minutes post-HFS | Estimate [95% CI] | Std. effect [95% CI] | P |
| HFS | 5 | -0.46 [-0.84, -0.09] | -0.99 [-1.63, -0.34] | 0.011 |
| HFS | 20 | -0.7 [-1.08, -0.32] | -1.5 [-2.15, -0.84] | <.001 |
| HFS | 35 | -0.34 [-0.72, 0.04] | -0.73 [-1.37, -0.09] | 0.088 |
| HFS | 50 | -0.58 [-0.96, -0.2] | -1.23 [-1.88, -0.58] | 0.001 |
| **Session 2 comparisons to baseline: Post-HFS – Baseline** | | | | |
| Arm | Minutes post-HFS | Estimate [95% CI] | Std. effect [95% CI] | *P* |
| HFS | 5 | -0.4 [-0.77, -0.02] | -0.85 [-1.49, -0.2] | 0.037 |
| HFS | 20 | -0.35 [-0.72, 0.03] | -0.74 [-1.38, -0.1] | 0.08 |
| HFS | 35 | -0.47 [-0.84, -0.09] | -1 [-1.64, -0.35] | 0.011 |
| HFS | 50 | -0.32 [-0.7, 0.06] | -0.68 [-1.32, -0.04] | 0.122 |

Supplementary Table 7 continued

**Electrical-evoked α-ERD magnitude results**

| | | | | |
|---|---|---|---|---|
| **b) Model assessment: Control arm** | | | | |
| Magnitude ~ time * session * (age + sex) + (1 \| subject) | | | | |
| Conditional $R^2$ = .81 | | Marginal $R^2$ = .06 | | |
| **Fixed effects and interactions** | | | | |
| Predictor | | *F* (DF) | *η2* [95% CI] | *P* |
| Time | | 1.12 (4, 243) | 0.02 [0, 1] | 0.35 |
| Session | | 0.25 (1, 243) | <.01 [0, 1] | 0.62 |
| Age | | 0 (1, 27) | <.01 [0, 1] | 0.95 |
| Sex | | 0.91 (1, 27) | 0.03 [0, 1] | 0.35 |
| Time: session | | 1.81 (4, 243) | 0.03 [0, 1] | 0.13 |
| Time: age | | 0.42 (4, 243) | <.01 [0, 1] | 0.80 |
| Time: sex | | 2.95 (4, 243) | 0.05 [0, 1] | 0.02 |
| Session: age | | 5.78 (1, 243) | 0.02 [0, 1] | 0.02 |
| Session: sex | | 4.46 (1, 243) | 0.02 [0, 1] | 0.04 |
| Time: session: age | | 1.95 (4, 243) | 0.03 [0, 1] | 0.10 |
| Time: session: sex | | 1.07 (4, 243) | 0.02 [0, 1] | 0.37 |
| **Between session comparisons: Session 2 – Session 1** | | | | |
| Arm | Minutes post-HFS | Estimate [95% CI] | Std. effect [95% CI] | *P* |
| Control | -10 | -0.04 [-0.32, 0.23] | -0.1 [-0.74, 0.54] | 0.752 |
| Control | 5 | 0.07 [-0.21, 0.34] | 0.16 [-0.48, 0.8] | 0.623 |
| Control | 20 | -0.19 [-0.47, 0.08] | -0.44 [-1.08, 0.2] | 0.168 |
| Control | 35 | 0.21 [-0.06, 0.49] | 0.49 [-0.15, 1.13] | 0.126 |
| Control | 50 | -0.19 [-0.46, 0.09] | -0.44 [-1.08, 0.2] | 0.173 |
| **Session 1 comparisons to baseline: Post-HFS – Baseline** | | | | |
| Arm | Minutes post-HFS | Estimate [95% CI] | Std. effect [95% CI] | *P* |
| Control | 5 | -0.17 [-0.52, 0.18] | -0.4 [-1.04, 0.24] | 0.529 |
| Control | 20 | 0.02 [-0.32, 0.37] | 0.06 [-0.58, 0.7] | 0.994 |
| Control | 35 | -0.25 [-0.59, 0.1] | -0.57 [-1.21, 0.07] | 0.235 |
| Control | 50 | -0.1 [-0.45, 0.24] | -0.24 [-0.88, 0.4] | 0.824 |
| **Session 2 comparisons to baseline: Post-HFS – Baseline** | | | | |
| Arm | Minutes post-HFS | Estimate [95% CI] | Std. effect [95% CI] | *P* |
| Control | 5 | -0.06 [-0.41, 0.29] | -0.14 [-0.78, 0.5] | 0.949 |
| Control | 20 | -0.12 [-0.47, 0.22] | -0.29 [-0.93, 0.35] | 0.744 |
| Control | 35 | 0.01 [-0.34, 0.36] | 0.02 [-0.61, 0.66] | 0.999 |
| Control | 50 | -0.25 [-0.6, 0.1] | -0.58 [-1.22, 0.06] | 0.225 |

Supplementary Table 7 continued

**Electrical-evoked α-ERD magnitude results**

| **c) Model assessment: Magnitude influence on pain ratings** | | | |
|---|---|---|---|
| Pain rating ~ time * session * arm * (magnitude change + magnitude baseline) + (1 \| subject) | | | |
| Conditional $R^2$ = .88 | Marginal $R^2$ = .04 | | |
| **Fixed effects and interactions** | | | |
| Predictor | *F* (DF) | *η2* [95% CI] | *P* |
| Time | 4.8 (1, 513.29) | <.01 [0, 1] | 0.29 |
| Session | 112.91 (1, 513.65) | 0.18 [0.13, 1] | <.001 |
| Arm | 0.47 (1, 513.37) | <.01 [0, 1] | 0.494 |
| Magnitude change | 0.01 (1, 520.86) | <.01 [0, 1] | 0.927 |
| Magnitude baseline | 3.69 (1, 535.8) | <.01 [0, 1] | 0.055 |
| Time: session | 2.03 (1, 513.02) | <.01 [0, 1] | 0.155 |
| Time: arm | 0.12 (1, 513.01) | <.01 [0, 1] | 0.732 |
| Session: arm | 0 (1, 513) | <.01 [0, 1] | 0.986 |
| Time: magnitude change | 0.46 (1, 513.19) | <.01 [0, 1] | 0.499 |
| Time: magnitude baseline | 0.42 (1, 512.99) | <.01 [0, 1] | 0.516 |
| Session: magnitude change | 1.82 (1, 513.71) | <.01 [0, 1] | 0.178 |
| Session: magnitude baseline | 0.66 (1, 514.31) | <.01 [0, 1] | 0.416 |
| Arm: magnitude change | 0.33 (1, 513.54) | <.01 [0, 1] | 0.565 |
| Arm: magnitude baseline | 0.13 (1, 513.76) | <.01 [0, 1] | 0.716 |
| Time: session: arm | 0.46 (1, 513.05) | <.01 [0, 1] | 0.498 |
| Time: session: magnitude change | 3.5 (1, 513.24) | <.01 [0, 1] | 0.062 |
| Time: session: magnitude baseline | 0.66 (1, 513.07) | <.01 [0, 1] | 0.417 |
| Time: arm: magnitude change | 0.01 (1, 513.13) | <.01 [0, 1] | 0.933 |
| Time: arm: magnitude baseline | 0.72 (1, 513.02) | <.01 [0, 1] | 0.396 |
| Session: arm: magnitude change | 4.67 (1, 514.21) | <.01 [0, 1] | 0.31 |
| Session: arm: magnitude baseline | 1.6 (1, 513.91) | <.01 [0, 1] | 0.207 |
| Time: session: arm: magnitude change | 0.22 (1, 513.75) | <.01 [0, 1] | 0.639 |
| Time: session: arm: magnitude baseline | 0.58 (1, 513) | <.01 [0, 1] | 0.447 |

*Supplementary Table 7 Electrical-evoked α-ERD mixed-effects modelling*

The linear mixed effects modelling results for α-ERD data are displayed for the a) HFS arm, b) control arm, and c) their relationship with MPS. HFS = High-frequency stimulation. α-ERD = alpha-band event-related desynchronisation. N = 27, (female = 19).

Supplementary Table 8

**Electrical-evoked θ-ERS magnitude results**

| **a) Model assessment: HFS arm** | | | | |
|---|---|---|---|---|
| Magnitude ~ time * session * (age + sex) + (1 \| subject) | | | | |
| Conditional $R^2$ = .62 | | Marginal $R^2$ = .09 | | |
| **Fixed effects and interactions** | | | | |
| Predictor | | *F* (DF) | *η2* [95% CI] | *p* |
| Time | | 8.2 (4, 243) | 0.12 [0.05, 1] | <.001 |
| Session | | 0.25 (1, 243) | <.01 [0, 1] | 0.615 |
| Age | | 0.07 (1, 27) | <.01 [0, 1] | 0.799 |
| Sex | | 0.58 (1, 27) | 0.02 [0, 1] | 0.452 |
| Time: session | | 0.59 (4, 243) | <.01 [0, 1] | 0.673 |
| Time: age | | 0.57 (4, 243) | <.01 [0, 1] | 0.681 |
| Time: sex | | 0.92 (4, 243) | 0.01 [0, 1] | 0.453 |
| Session: age | | 0.35 (1, 243) | <.01 [0, 1] | 0.556 |
| Session: sex | | 0.4 (1, 243) | <.01 [0, 1] | 0.527 |
| Time: session: age | | 1.18 (4, 243) | 0.02 [0, 1] | 0.318 |
| Time: session: sex | | 0.35 (4, 243) | <.01 [0, 1] | 0.847 |
| **Between session comparisons: Session 2 – Session 1** | | | | |
| Arm | Minutes post-HFS | Estimate [95% CI] | Std. effect [95% CI] | *P* |
| HFS | -10 | -0.1 [-0.49, 0.29] | -0.17 [-0.8, 0.46] | 0.598 |
| HFS | 5 | -0.05 [-0.44, 0.34] | -0.08 [-0.72, 0.55] | 0.792 |
| HFS | 20 | 0.1 [-0.29, 0.49] | 0.17 [-0.46, 0.8] | 0.595 |
| HFS | 35 | 0.08 [-0.31, 0.47] | 0.13 [-0.5, 0.77] | 0.675 |
| HFS | 50 | -0.24 [-0.63, 0.15] | -0.39 [-1.02, 0.24] | 0.225 |
| **Session 1 comparisons to baseline: Post-HFS – Baseline** | | | | |
| Arm | Minutes post-HFS | Estimate [95% CI] | Std. effect [95% CI] | P |
| HFS | 5 | -0.22 [-0.71, 0.27] | -0.35 [-0.99, 0.28] | 0.614 |
| HFS | 20 | -0.31 [-0.8, 0.18] | -0.5 [-1.13, 0.13] | 0.332 |
| HFS | 35 | -0.53 [-1.02, -0.04] | -0.86 [-1.5, -0.23] | 0.029 |
| HFS | 50 | -0.62 [-1.11, -0.13] | -1 [-1.64, -0.36] | 0.008 |
| **Session 2 comparisons to baseline: Post-HFS – Baseline** | | | | |
| Arm | Minutes post-HFS | Estimate [95% CI] | Std. effect [95% CI] | *P* |
| HFS | 5 | -0.17 [-0.66, 0.32] | -0.27 [-0.9, 0.36] | 0.774 |
| HFS | 20 | -0.1 [-0.59, 0.39] | -0.16 [-0.8, 0.47] | 0.925 |
| HFS | 35 | -0.35 [-0.84, 0.14] | -0.56 [-1.2, 0.07] | 0.241 |
| HFS | 50 | -0.75 [-1.24, -0.26] | -1.22 [-1.86, -0.58] | <.001 |

Supplementary Table 8 continued

**Electrical-evoked θ-ERS magnitude results**

| | | | | |
|---|---|---|---|---|
| **b) Model assessment: Control arm** | | | | |
| Magnitude ~ time * session * (age + sex) + (1 \| subject) | | | | |
| Conditional $R^2$ = .57 | | Marginal $R^2$ = .06 | | |
| **Fixed effects and interactions** | | | | |
| Predictor | | *F* (DF) | *η2* [95% CI] | *P* |
| Time | | 2.58 (4, 243) | 0.04 [0, 1] | 0.04 |
| Session | | 0.53 (1, 243) | <.01 [0, 1] | 0.47 |
| Age | | 0.27 (1, 27) | <.01 [0, 1] | 0.61 |
| Sex | | 0.24 (1, 27) | <.01 [0, 1] | 0.63 |
| Time: session | | 1.16 (4, 243) | 0.02 [0, 1] | 0.33 |
| Time: age | | 0.73 (4, 243) | 0.01 [0, 1] | 0.58 |
| Time: sex | | 0.21 (4, 243) | <.01 [0, 1] | 0.93 |
| Session: age | | 3.5 (1, 243) | 0.01 [0, 1] | 0.06 |
| Session: sex | | 3.05 (1, 243) | 0.01 [0, 1] | 0.08 |
| Time: session: age | | 0.36 (4, 243) | <.01 [0, 1] | 0.84 |
| Time: session: sex | | 0.18 (4, 243) | <.01 [0, 1] | 0.95 |
| **Between session comparisons: Session 2 – Session 1** | | | | |
| Arm | Minutes post-HFS | Estimate [95% CI] | Std. effect [95% CI] | *P* |
| Control | -10 | -0.15 [-0.57, 0.26] | -0.23 [-0.86, 0.4] | 0.468 |
| Control | 5 | 0.13 [-0.28, 0.55] | 0.2 [-0.43, 0.84] | 0.521 |
| Control | 20 | -0.03 [-0.44, 0.38] | -0.05 [-0.68, 0.58] | 0.882 |
| Control | 35 | -0.03 [-0.44, 0.39] | -0.04 [-0.67, 0.59] | 0.891 |
| Control | 50 | 0.4 [-0.02, 0.81] | 0.6 [-0.03, 1.24] | 0.061 |
| **Session 1 comparisons to baseline: Post-HFS – Baseline** | | | | |
| Arm | Minutes post-HFS | Estimate [95% CI] | Std. effect [95% CI] | *P* |
| Control | 5 | -0.3 [-0.82, 0.22] | -0.46 [-1.1, 0.17] | 0.398 |
| Control | 20 | -0.35 [-0.87, 0.17] | -0.53 [-1.16, 0.1] | 0.284 |
| Control | 35 | -0.44 [-0.96, 0.08] | -0.67 [-1.3, -0.04] | 0.123 |
| Control | 50 | -0.63 [-1.15, -0.11] | -0.96 [-1.6, -0.33] | 0.012 |
| **Session 2 comparisons to baseline: Post-HFS – Baseline** | | | | |
| Arm | Minutes post-HFS | Estimate [95% CI] | Std. effect [95% CI] | *P* |
| Control | 5 | -0.02 [-0.54, 0.5] | -0.03 [-0.66, 0.6] | 0.999 |
| Control | 20 | -0.23 [-0.75, 0.29] | -0.35 [-0.98, 0.28] | 0.625 |
| Control | 35 | -0.32 [-0.84, 0.2] | -0.48 [-1.11, 0.15] | 0.365 |
| Control | 50 | -0.09 [-0.61, 0.44] | -0.13 [-0.76, 0.5] | 0.957 |

Supplementary Table 8 continued

**Electrical-evoked θ-ERS magnitude results**

| **c) Model assessment: Magnitude influence on pain ratings** | | | |
|---|---|---|---|
| Pain rating ~ time * session * arm * (magnitude change + magnitude baseline) + (1 \| subject) | | | |
| Conditional $R^2$ = .88 | Marginal $R^2$ = .39 | | |
| **Fixed effects and interactions** | | | |
| Predictor | *F* (DF) | *η2* [95% CI] | *P* |
| Time | 3.52 (1, 513.36) | <.01 [0, 1] | 0.061 |
| Session | 101.79 (1, 513.35) | 0.17 [0.12, 1] | <.001 |
| Arm | 0.03 (1, 513.39) | <.01 [0, 1] | 0.867 |
| Magnitude change | 2.05 (1, 518.77) | <.01 [0, 1] | 0.152 |
| Magnitude baseline | 3.76 (1, 530.93) | <.01 [0, 1] | 0.053 |
| Time: session | 3.71 (1, 513.03) | <.01 [0, 1] | 0.055 |
| Time: arm | 0.67 (1, 513.06) | <.01 [0, 1] | 0.414 |
| Session: arm | 0.16 (1, 513.1) | <.01 [0, 1] | 0.692 |
| Time: magnitude change | 0.09 (1, 513.17) | <.01 [0, 1] | 0.758 |
| Time: magnitude baseline | 0.68 (1, 513.22) | <.01 [0, 1] | 0.41 |
| Session: magnitude change | 1.71 (1, 514.65) | <.01 [0, 1] | 0.191 |
| Session: magnitude baseline | 24.49 (1, 514.61) | 0.05 [0.02, 1] | <.001 |
| Arm: magnitude change | 0 (1, 513.52) | <.01 [0, 1] | 0.963 |
| Arm: magnitude baseline | 1.4 (1, 513.36) | <.01 [0, 1] | 0.237 |
| Time: session: arm | 0.49 (1, 513.07) | <.01 [0, 1] | 0.483 |
| Time: session: magnitude change | 0.05 (1, 513.17) | <.01 [0, 1] | 0.824 |
| Time: session: magnitude baseline | 0.98 (1, 513.03) | <.01 [0, 1] | 0.323 |
| Time: arm: magnitude change | 0.51 (1, 513.18) | <.01 [0, 1] | 0.477 |
| Time: arm: magnitude baseline | 0.44 (1, 513.02) | <.01 [0, 1] | 0.508 |
| Session: arm: magnitude change | 0.03 (1, 513.91) | <.01 [0, 1] | 0.857 |
| Session: arm: magnitude baseline | 0.37 (1, 513.55) | <.01 [0, 1] | 0.546 |
| Time: session: arm: magnitude change | 0.14 (1, 513.18) | <.01 [0, 1] | 0.71 |
| Time: session: arm: magnitude baseline | 0 (1, 513.04) | <.01 [0, 1] | 0.973 |

*Supplementary Table 8 Electrical-evoked θ-ERS mixed-effects modelling*

The linear mixed effects modelling results for θ-ERS data are displayed for the a) HFS arm, b) control arm, and c) their relationship with MPS. HFS = High-frequency stimulation. θ-ERS = theta-band event-related synchronisation. N = 27, (female = 19).

Supplementary Table 9

**Electrical-evoked γ-ERS magnitude results**

| | | | | |
|---|---|---|---|---|
| **a) Model assessment: HFS arm** | | | | |
| Magnitude ~ time * session * (age + sex) + (1 \| subject) | | | | |
| Conditional $R^2$ = .46 | | Marginal $R^2$ = .08 | | |
| **Fixed effects and interactions** | | | | |
| Predictor | | *F* (DF) | *η2* [95% CI] | *p* |
| Time | | 1.86 (4, 243) | .03 [0, 1] | .12 |
| Session | | .22 (1, 243) | <.01 [0, 1] | .64 |
| Age | | .11 (1, 27) | <.01 [0, 1] | .74 |
| Sex | | .3 (1, 27) | .01 [0, 1] | .59 |
| Time: session | | 1.48 (4, 243) | .02 [0, 1] | .21 |
| Time: age | | .83 (4, 243) | .01 [0, 1] | .50 |
| Time: sex | | 1.09 (4, 243) | .02 [0, 1] | .36 |
| Session: age | | .29 (1, 243) | <.01 [0, 1] | .59 |
| Session: sex | | 10.41 (1, 243) | .04 [.01, 1] | .001 |
| Time: session: age | | .19 (4, 243) | <.01 [0, 1] | .94 |
| Time: session: sex | | .22 (4, 243) | <.01 [0, 1] | .93 |
| **Between session comparisons: Session 2 – Session 1** | | | | |
| Arm | Minutes post-HFS | Estimate [95% CI] | Std. effect [95% CI] | *P* |
| HFS | -10 | -.24 [-.7, .22] | -.33 [-.96, .3] | .298 |
| HFS | 5 | .1 [-.36, .56] | .14 [-.49, .76] | .671 |
| HFS | 20 | -.18 [-.64, .28] | -.25 [-.88, .38] | .429 |
| HFS | 35 | .15 [-.31, .6] | .2 [-.43, .83] | .532 |
| HFS | 50 | .41 [-.05, .87] | .57 [-.06, 1.2] | .077 |
| **Session 1 comparisons to baseline: Post-HFS – Baseline** | | | | |
| Arm | Minutes post-HFS | Estimate [95% CI] | Std. effect [95% CI] | P |
| HFS | 5 | -.26 [-.84, .32] | -.35 [-.98, .27] | .610 |
| HFS | 20 | .01 [-.57, .59] | .01 [-.62, .64] | 1 |
| HFS | 35 | .06 [-.51, .64] | .09 [-.54, .71] | .984 |
| HFS | 50 | -.1 [-.68, .47] | -.14 [-.77, .49] | .949 |
| **Session 2 comparisons to baseline: Post-HFS – Baseline** | | | | |
| Arm | Minutes post-HFS | Estimate [95% CI] | Std. effect [95% CI] | *P* |
| HFS | 5 | .08 [-.49, .66] | .11 [-.52, .74] | .970 |
| HFS | 20 | .07 [-.51, .64] | .09 [-.54, .72] | .981 |
| HFS | 35 | .45 [-.13, 1.03] | .62 [-.01, 1.25] | .172 |
| HFS | 50 | .55 [-.02, 1.13] | .76 [.13, 1.39] | .064 |

Supplementary Table 9 continued

**Electrical-evoked γ-ERS magnitude results**

| **b) Model assessment: Control arm** | | | | |
|---|---|---|---|---|
| Magnitude ~ time * session * (age + sex) + (1 \| subject) | | | | |
| Conditional $R^2$ = .34 | | Marginal $R^2$ = .03 | | |
| **Fixed effects and interactions** | | | | |
| Predictor | | *F* (DF) | *η2* [95% CI] | *P* |
| Time | | .69 (4, 243) | .01 [0, 1] | .60 |
| Session | | .17 (1, 243) | <.01 [0, 1] | .68 |
| Age | | (1, 27) | <.01 [0, 1] | 1 |
| Sex | | .05 (1, 27) | <.01 [0, 1] | .82 |
| Time: session | | .34 (4, 243) | <.01 [0, 1] | .85 |
| Time: age | | .31 (4, 243) | <.01 [0, 1] | .87 |
| Time: sex | | .89 (4, 243) | .01 [0, 1] | .47 |
| Session: age | | (1, 243) | <.01 [0, 1] | .98 |
| Session: sex | | .11 (1, 243) | <.01 [0, 1] | .74 |
| Time: session: age | | .49 (4, 243) | <.01 [0, 1] | .74 |
| Time: session: sex | | .61 (4, 243) | <.01 [0, 1] | .66 |
| **Between session comparisons: Session 2 – Session 1** | | | | |
| Arm | Minutes post-HFS | Estimate [95% CI] | Std. effect [95% CI] | *P* |
| Control | -10 | .04 [-.47, .55] | .05 [-.57, .68] | .869 |
| Control | 5 | .22 [-.29, .73] | .27 [-.36, .89] | .400 |
| Control | 20 | .08 [-.43, .59] | .1 [-.53, .72] | .763 |
| Control | 35 | -.18 [-.69, .33] | -.22 [-.85, .41] | .491 |
| Control | 50 | .06 [-.45, .57] | .08 [-.55, .7] | .806 |
| **Session 1 comparisons to baseline: Post-HFS – Baseline** | | | | |
| Arm | Minutes post-HFS | Estimate [95% CI] | Std. effect [95% CI] | *P* |
| Control | 5 | -.18 [-.82, .46] | -.22 [-.85, .4] | .852 |
| Control | 20 | .06 [-.58, .7] | .07 [-.55, .7] | .988 |
| Control | 35 | .29 [-.35, .93] | .36 [-.27, .98] | .609 |
| Control | 50 | .06 [-.58, .7] | .07 [-.56, .7] | .990 |
| **Session 2 comparisons to baseline: Post-HFS – Baseline** | | | | |
| Arm | Minutes post-HFS | Estimate [95% CI] | Std. effect [95% CI] | *P* |
| Control | 5 | -.01 [-.64, .63] | -.01 [-.63, .62] | 1 |
| Control | 20 | .1 [-.54, .73] | .12 [-.51, .74] | .965 |
| Control | 35 | .07 [-.57, .71] | .08 [-.54, .71] | .985 |
| Control | 50 | .08 [-.56, .72] | .1 [-.53, .72] | .979 |

Supplementary Table 9 continued

**Electrical-evoked γ-ERS magnitude results**

| **c) Model assessment: Magnitude influence on pain ratings** | | | |
|---|---|---|---|
| Pain rating ~ time * session * arm * (magnitude change + magnitude baseline) + (1 \| subject) | | | |
| Conditional $R^2$ = .9 | Marginal $R^2$ = .05 | | |
| **Fixed effects and interactions** | | | |
| Predictor | *F* (DF) | *η2* [95% CI] | *P* |
| Time | 4.04 (1, 512.85) | <.01 [0, 1] | .051 |
| Session | 128.62 (1, 512.98) | 0.2 [0.15, 1] | <.001 |
| Arm | 3.75 (1, 512.94) | <.01 [0, 1] | .053 |
| Magnitude change | 3.99 (1, 514.48) | <.01 [0, 1] | .046 |
| Magnitude baseline | 19.58 (1, 522.34) | 0.04 [0.01, 1] | <.001 |
| Time: session | 2.74 (1, 512.84) | <.01 [0, 1] | .098 |
| Time: arm | .4 (1, 512.84) | <.01 [0, 1] | .526 |
| Session: arm | 1.47 (1, 513.44) | <.01 [0, 1] | .226 |
| Time: magnitude change | 1.5 (1, 513) | <.01 [0, 1] | .222 |
| Time: magnitude baseline | 2.63 (1, 512.99) | <.01 [0, 1] | .105 |
| Session: magnitude change | .11 (1, 513.08) | <.01 [0, 1] | .738 |
| Session: magnitude baseline | 29.06 (1, 513.99) | 0.05 [0.03, 1] | <.001 |
| Arm: magnitude change | 7.04 (1, 513.17) | 0.01 [0, 1] | .008 |
| Arm: magnitude baseline | 3.01 (1, 513.26) | <.01 [0, 1] | .083 |
| Time: session: arm | .37 (1, 512.84) | <.01 [0, 1] | .544 |
| Time: session: magnitude change | 2.97 (1, 513.01) | <.01 [0, 1] | .086 |
| Time: session: magnitude baseline | 3.25 (1, 512.88) | <.01 [0, 1] | .072 |
| Time: arm: magnitude change | 2.31 (1, 512.95) | <.01 [0, 1] | .13 |
| Time: arm: magnitude baseline | 3.69 (1, 512.88) | <.01 [0, 1] | .055 |
| Session: arm: magnitude change | .15 (1, 513.5) | <.01 [0, 1] | .703 |
| Session: arm: magnitude baseline | .39 (1, 513.78) | <.01 [0, 1] | .535 |
| Time: session: arm: magnitude change | .13 (1, 513.16) | <.01 [0, 1] | .722 |
| Time: session: arm: magnitude baseline | .07 (1, 512.94) | <.01 [0, 1] | .798 |

*Supplementary Table 9 Electrical-evoked γ-ERS mixed-effects modelling*

The linear mixed effects modelling results for γ-ERS data are displayed for the a) HFS arm, b) control arm, and c) their relationship with MPS. HFS = High-frequency stimulation. γ-ERS = gamma-band event-related synchronisation. N = 27, (female = 19).

Supplementary Table 10

**Electrical-evoked VP magnitude results**

| | | | | |
|---|---|---|---|---|
| **a) Model assessment: HFS arm** | | | | |
| Magnitude ~ time * session * (age + sex) + (1 \| subject) | | | | |
| Conditional $R^2$ = .68 | | Marginal $R^2$ = .19 | | |
| **Fixed effects and interactions** | | | | |
| Predictor | | *F* (DF) | *η2* [95% CI] | *p* |
| Time | | 3.94 (4, 243) | 0.06 [0.01, 1] | 0.004 |
| Session | | 30.9 (1, 243) | 0.11 [0.06, 1] | <.001 |
| Age | | 0.94 (1, 27) | 0.03 [0, 1] | 0.341 |
| Sex | | 3.87 (1, 27) | 0.13 [0, 1] | 0.06 |
| Time: session | | 0.31 (4, 243) | <.01 [0, 1] | 0.873 |
| Time: age | | 0.24 (4, 243) | <.01 [0, 1] | 0.913 |
| Time: sex | | 0.71 (4, 243) | 0.01 [0, 1] | 0.586 |
| Session: age | | 1.1 (1, 243) | <.01 [0, 1] | 0.296 |
| Session: sex | | 4.12 (1, 243) | 0.02 [0, 1] | 0.043 |
| Time: session: age | | 0.37 (4, 243) | <.01 [0, 1] | 0.833 |
| Time: session: sex | | 0.52 (4, 243) | <.01 [0, 1] | 0.723 |
| **Between session comparisons: Session 2 – Session 1** | | | | |
| Arm | Minutes post-HFS | Estimate [95% CI] | Std. effect [95% CI] | *P* |
| HFS | -10 | -1.43 [-3.08, 0.22] | -0.55 [-1.18, 0.09] | 0.089 |
| HFS | 5 | -2.44 [-4.09, -0.79] | -0.93 [-1.57, -0.3] | 0.004 |
| HFS | 20 | -1.56 [-3.21, 0.09] | -0.6 [-1.23, 0.04] | 0.064 |
| HFS | 35 | -2.07 [-3.72, -0.42] | -0.79 [-1.43, -0.16] | 0.015 |
| HFS | 50 | -2.21 [-3.86, -0.55] | -0.84 [-1.48, -0.21] | 0.010 |
| **Session 1 comparisons to baseline: Post-HFS – Baseline** | | | | |
| Arm | Minutes post-HFS | Estimate [95% CI] | Std. effect [95% CI] | P |
| HFS | 5 | 1.42 [-0.66, 3.5] | 0.54 [-0.09, 1.18] | 0.266 |
| HFS | 20 | 0.42 [-1.66, 2.5] | 0.16 [-0.47, 0.79] | 0.929 |
| HFS | 35 | -0.39 [-2.47, 1.69] | -0.15 [-0.78, 0.48] | 0.941 |
| HFS | 50 | -0.59 [-2.67, 1.49] | -0.23 [-0.86, 0.41] | 0.844 |
| **Session 2 comparisons to baseline: Post-HFS – Baseline** | | | | |
| Arm | Minutes post-HFS | Estimate [95% CI] | Std. effect [95% CI] | *P* |
| HFS | 5 | 0.41 [-1.67, 2.49] | 0.16 [-0.48, 0.79] | 0.933 |
| HFS | 20 | 0.29 [-1.79, 2.37] | 0.11 [-0.52, 0.74] | 0.970 |
| HFS | 35 | -1.03 [-3.11, 1.05] | -0.4 [-1.03, 0.24] | 0.530 |
| HFS | 50 | -1.37 [-3.45, 0.71] | -0.53 [-1.16, 0.11] | 0.295 |

Supplementary Table 10 continued

**Electrical-evoked VP magnitude results**

| | | | | |
|---|---|---|---|---|
| **b) Model assessment: Control arm** | | | | |
| Magnitude ~ time * session * (age + sex) + (1 \| subject) | | | | |
| Conditional $R^2$ = .71 | | Marginal $R^2$ = .23 | | |
| **Fixed effects and interactions** | | | | |
| Predictor | | *F* (DF) | *η2* [95% CI] | *P* |
| Time | | 2.21 (4, 243) | 0.04 [0, 1] | 0.069 |
| Session | | 14.34 (1, 243) | 0.06 [0.02, 1] | <.001 |
| Age | | 0.44 (1, 27) | 0.02 [0, 1] | 0.511 |
| Sex | | 7.2 (1, 27) | 0.21 [0.03, 1] | 0.012 |
| Time: session | | 0.27 (4, 243) | <.01 [0, 1] | 0.896 |
| Time: age | | 0.18 (4, 243) | <.01 [0, 1] | 0.946 |
| Time: sex | | 1.84 (4, 243) | 0.03 [0, 1] | 0.122 |
| Session: age | | 0.05 (1, 243) | <.01 [0, 1] | 0.823 |
| Session: sex | | 2.37 (1, 243) | <.01 [0, 1] | 0.125 |
| Time: session: age | | 0.95 (4, 243) | 0.02 [0, 1] | 0.434 |
| Time: session: sex | | 0.69 (4, 243) | 0.01 [0, 1] | 0.597 |
| **Between session comparisons: Session 2 – Session 1** | | | | |
| Arm | Minutes post-HFS | Estimate [95% CI] | Std. effect [95% CI] | *P* |
| Control | -10 | -1.39 [-2.92, 0.14] | -0.58 [-1.21, 0.06] | 0.074 |
| Control | 5 | -0.75 [-2.28, 0.78] | -0.31 [-0.94, 0.32] | 0.332 |
| Control | 20 | -1.12 [-2.65, 0.41] | -0.46 [-1.1, 0.17] | 0.148 |
| Control | 35 | -1.1 [-2.62, 0.43] | -0.45 [-1.09, 0.18] | 0.156 |
| Control | 50 | -1.76 [-3.28, -0.23] | -0.73 [-1.36, -0.09] | 0.025 |
| **Session 1 comparisons to baseline: Post-HFS – Baseline** | | | | |
| Arm | Minutes post-HFS | Estimate [95% CI] | Std. effect [95% CI] | *P* |
| Control | 5 | -0.54 [-2.47, 1.38] | -0.22 [-0.86, 0.41] | 0.848 |
| Control | 20 | -0.94 [-2.86, 0.99] | -0.39 [-1.02, 0.25] | 0.544 |
| Control | 35 | -1.5 [-3.42, 0.43] | -0.62 [-1.26, 0.02] | 0.172 |
| Control | 50 | -0.64 [-2.57, 1.28] | -0.27 [-0.9, 0.37] | 0.782 |
| **Session 2 comparisons to baseline: Post-HFS – Baseline** | | | | |
| Arm | Minutes post-HFS | Estimate [95% CI] | Std. effect [95% CI] | *P* |
| Control | 5 | 0.1 [-1.83, 2.02] | 0.04 [-0.59, 0.67] | 0.997 |
| Control | 20 | -0.67 [-2.59, 1.26] | -0.28 [-0.91, 0.36] | 0.762 |
| Control | 35 | -1.2 [-3.13, 0.72] | -0.5 [-1.13, 0.14] | 0.338 |
| Control | 50 | -1.01 [-2.93, 0.92] | -0.42 [-1.05, 0.22] | 0.486 |

Supplementary Table 10 continued

**Electrical-evoked VP magnitude results**

| **c) Model assessment: Magnitude influence on pain ratings** | | | |
|---|---|---|---|
| Pain rating ~ time * session * arm * (magnitude change + magnitude baseline) + (1 \| subject) | | | |
| Conditional $R^2$ = .88 | Marginal $R^2$ = .05 | | |
| **Fixed effects and interactions** | | | |
| Predictor | *F* (DF) | *η2* [95% CI] | *P* |
| Time | 1.46 (1, 513) | <.01 [0, 1] | 0.227 |
| Session | 77.88 (1, 515) | 0.13 [0.09, 1] | <.001 |
| Arm | 0 (1, 513) | <.01 [0, 1] | 0.956 |
| Magnitude change | 6.41 (1,516) | 0.01 [0, 1] | 0.012 |
| Magnitude baseline | 17.46 (1, 530) | 0.03 [0.01, 1] | <.001 |
| Time: session | 1.26 (1, 513) | <.01 [0, 1] | 0.263 |
| Time: arm | 0 (1, 513) | <.01 [0, 1] | 0.982 |
| Session: arm | 0.38 (1, 513) | <.01 [0, 1] | 0.536 |
| Time: magnitude change | 1.64 (1, 513) | <.01 [0, 1] | 0.201 |
| Time: magnitude baseline | 2.55 (1, 513) | <.01 [0, 1] | 0.111 |
| Session: magnitude change | 3.36 (1, 514) | <.01 [0, 1] | 0.068 |
| Session: magnitude baseline | 10.21 (1, 514) | 0.02 [0, 1] | 0.001 |
| Arm: magnitude change | 1.64 (1, 513) | <.01 [0, 1] | 0.201 |
| Arm: magnitude baseline | 0.07 (1, 513) | <.01 [0, 1] | 0.79 |
| Time: session: arm | 0.29 (1, 513) | <.01 [0, 1] | 0.589 |
| Time: session: magnitude change | 0.98 (1, 513) | <.01 [0, 1] | 0.323 |
| Time: session: magnitude baseline | 1.21 (1, 513) | <.01 [0, 1] | 0.272 |
| Time: arm: magnitude change | 0.43 (1, 513) | <.01 [0, 1] | 0.511 |
| Time: arm: magnitude baseline | 0.9 (1, 513) | <.01 [0, 1] | 0.344 |
| Session: arm: magnitude change | 0.45 (1, 513) | <.01 [0, 1] | 0.501 |
| Session: arm: magnitude baseline | 1.13 (1, 513) | <.01 [0, 1] | 0.289 |
| Time: session: arm: magnitude change | 0 (1, 513) | <.01 [0, 1] | 0.992 |
| Time: session: arm: magnitude baseline | 0.65 (1. 513) | <.01 [0, 1] | 0.422 |

*Supplementary Table 10 Electrical-evoked VP magnitude mixed-effects modelling*

The linear mixed effects modelling results for VP magnitude data are displayed for the a) HFS arm, b) control arm, and c) their relationship with MPS. HFS = High-frequency stimulation. VP = Vertex potential. N = 27, (female = 19).

Supplementary Table 11

**Electrical-evoked VP latency results**

| **a) Model assessment: HFS arm** | | | | |
|---|---|---|---|---|
| Latency ~ time * session * (age + sex) + (1 \| subject) | | | | |
| Conditional $R^2$ = .51 | | Marginal $R^2$ = .14 | | |
| **Fixed effects and interactions** | | | | |
| Predictor | | *F* (DF) | *η2* [95% CI] | *p* |
| Time | | 2.89 (4, 243) | 0.05 [0, 1] | 0.02 |
| Session | | 0 (1, 243) | <.01 [0, 1] | 0.99 |
| Age | | 0.1 (1, 243) | <.01 [0, 1] | 0.76 |
| Sex | | 3.31 (1, 27) | 0.11 [0, 1] | 0.08 |
| Time: session | | 0.14 (4, 243) | <.01 [0, 1] | 0.97 |
| Time: age | | 3.61 (4, 243) | 0.06 [0.01, 1] | 0.01 |
| Time: sex | | 1.1 (4, 243) | 0.02 [0, 1] | 0.36 |
| Session: age | | 6.65 (1, 243) | 0.03 [0, 1] | 0.01 |
| Session: sex | | 0.7 (1, 243) | <.01 [0, 1] | 0.40 |
| Time: session: age | | 0.85 (4, 243) | 0.01 [0, 1] | 0.49 |
| Time: session: sex | | 0.53 (4, 243) | <.01 [0, 1] | 0.71 |
| **Between session comparisons: Session 2 – Session 1** | | | | |
| Arm | Minutes post-HFS | Estimate [95% CI] | Std. effect [95% CI] | *P* |
| HFS | -10 | 0.03 [-0.4, 0.47] | 0.05 [-0.58, 0.68] | 0.879 |
| HFS | 5 | -0.05 [-0.49, 0.39] | -0.07 [-0.7, 0.56] | 0.831 |
| HFS | 20 | 0.03 [-0.41, 0.47] | 0.04 [-0.59, 0.67] | 0.901 |
| HFS | 35 | -0.1 [-0.54, 0.34] | -0.15 [-0.78, 0.48] | 0.643 |
| HFS | 50 | 0.09 [-0.34, 0.53] | 0.14 [-0.49, 0.76] | 0.671 |
| **Session 1 comparisons to baseline: Post-HFS – Baseline** | | | | |
| Arm | Minutes post-HFS | Estimate [95% CI] | Std. effect [95% CI] | P |
| HFS | 5 | 0.12 [-0.43, 0.67] | 0.17 [-0.45, 0.8] | 0.915 |
| HFS | 20 | -0.19 [-0.75, 0.36] | -0.28 [-0.91, 0.35] | 0.757 |
| HFS | 35 | -0.02 [-0.57, 0.53] | -0.03 [-0.66, 0.59] | 0.998 |
| HFS | 50 | -0.4 [-0.95, 0.15] | -0.58 [-1.21, 0.05] | 0.217 |
| **Session 2 comparisons to baseline: Post-HFS – Baseline** | | | | |
| Arm | Minutes post-HFS | Estimate [95% CI] | Std. effect [95% CI] | *P* |
| HFS | 5 | 0.04 [-0.51, 0.59] | 0.06 [-0.57, 0.69] | 0.994 |
| HFS | 20 | -0.2 [-0.75, 0.35] | -0.29 [-0.92, 0.34] | 0.741 |
| HFS | 35 | -0.16 [-0.71, 0.39] | -0.23 [-0.86, 0.4] | 0.841 |
| HFS | 50 | -0.34 [-0.89, 0.21] | -0.49 [-1.12, 0.14] | 0.347 |

Supplementary Table 11 continued

**Electrical-evoked VP latency results**

| | | | | |
|---|---|---|---|---|
| **b) Model assessment: Control arm** | | | | |
| Latency ~ time * session * (age + sex) + (1 \| subject) | | | | |
| Conditional $R^2$ = .54 | | Marginal $R^2$ = .11 | | |
| **Fixed effects and interactions** | | | | |
| Predictor | | *F* (DF) | *η2* [95% CI] | *P* |
| Time | | 2.42 (4, 243) | 0.04 [0, 1] | 0.05 |
| Session | | 0.15 (1, 243) | <.01 [0, 1] | 0.70 |
| Age | | 0.01 (1, 27) | <.01 [0, 1] | 0.94 |
| Sex | | 1.09 (1, 27) | 0.04 [0, 1] | 0.30 |
| Time: session | | 1.67 (4, 243) | 0.03 [0, 1] | 0.16 |
| Time: age | | 1.41 (4, 243) | 0.02 [0, 1] | 0.23 |
| Time: sex | | 1.16 (4, 243) | 0.02 [0, 1] | 0.33 |
| Session: age | | 0.08 (1, 243) | <.01 [0, 1] | 0.77 |
| Session: sex | | 7.42 (1, 243) | 0.03 [0, 1] | 0.01 |
| Time: session: age | | 0.71 (4, 243) | 0.01 [0, 1] | 0.59 |
| Time: session: sex | | 0.37 (4, 243) | <.01 [0, 1] | 0.83 |
| **Between session comparisons: Session 2 – Session 1** | | | | |
| Arm | Minutes post-HFS | Estimate [95% CI] | Std. effect [95% CI] | *P* |
| Control | -10 | -0.18 [-0.61, 0.24] | -0.27 [-0.9, 0.36] | 0.393 |
| Control | 5 | -0.42 [-0.84, 0.01] | -0.61 [-1.25, 0.02] | 0.055 |
| Control | 20 | 0.22 [-0.21, 0.64] | 0.32 [-0.31, 0.95] | 0.315 |
| Control | 35 | 0.15 [-0.27, 0.58] | 0.23 [-0.4, 0.86] | 0.477 |
| Control | 50 | 0.05 [-0.37, 0.48] | 0.08 [-0.55, 0.71] | 0.803 |
| **Session 1 comparisons to baseline: Post-HFS – Baseline** | | | | |
| Arm | Minutes post-HFS | Estimate [95% CI] | Std. effect [95% CI] | *P* |
| Control | 5 | 0.02 [-0.52, 0.56] | 0.03 [-0.6, 0.66] | 0.999 |
| Control | 20 | -0.38 [-0.92, 0.16] | -0.56 [-1.19, 0.07] | 0.245 |
| Control | 35 | -0.57 [-1.1, -0.03] | -0.84 [-1.47, -0.2] | 0.035 |
| Control | 50 | -0.42 [-0.95, 0.12] | -0.61 [-1.24, 0.02] | 0.176 |
| **Session 2 comparisons to baseline: Post-HFS – Baseline** | | | | |
| Arm | Minutes post-HFS | Estimate [95% CI] | Std. effect [95% CI] | *P* |
| Control | 5 | -0.21 [-0.75, 0.32] | -0.32 [-0.94, 0.31] | 0.689 |
| Control | 20 | 0.02 [-0.51, 0.56] | 0.03 [-0.59, 0.66] | 0.998 |
| Control | 35 | -0.23 [-0.77, 0.31] | -0.34 [-0.97, 0.29] | 0.644 |
| Control | 50 | -0.18 [-0.71, 0.36] | -0.26 [-0.89, 0.37] | 0.788 |

Supplementary Table 11 continued

**Electrical-evoked VP latency results**

| **c) Model assessment: Latency influence on pain ratings** | | | |
|---|---|---|---|
| Pain rating ~ time * session * arm * (latency change + latency baseline) + (1 \| subject) | | | |
| Conditional $R^2$ = .88 | Marginal $R^2$ = .04 | | |
| **Fixed effects and interactions** | | | |
| Predictor | *F* (DF) | *η2* [95% CI] | *P* |
| Time | 4.35 (1, 513) | <.01 [0, 1] | 0.37 |
| Session | 98.02 (1, 513) | 0.16 [0.12, 1] | <.001 |
| Arm | 0.48 (1, 513) | <.01 [0, 1] | 0.487 |
| Latency change | 0 (1, 516) | <.01 [0, 1] | 0.958 |
| Latency baseline | 1.07 (1, 522) | <.01 [0, 1] | 0.302 |
| Time: session | 3.64 (1, 513) | <.01 [0, 1] | 0.057 |
| Time: arm | 0.58 (1, 513) | <.01 [0, 1] | 0.445 |
| Session: arm | 0.58 (1, 513) | <.01 [0, 1] | 0.448 |
| Time: latency change | 0.02 (1, 514) | <.01 [0, 1] | 0.893 |
| Time: latency baseline | 0.59 (1, 514) | <.01 [0, 1] | 0.443 |
| Session: latency change | 0.17 (1, 514) | <.01 [0, 1] | 0.682 |
| Session: latency baseline | 1.89 (1, 514) | <.01 [0, 1] | 0.17 |
| Arm: latency change | 0.29 (1, 514) | <.01 [0, 1] | 0.588 |
| Arm: latency baseline | 2.61 (1, 514) | <.01 [0, 1] | 0.107 |
| Time: session: arm | 0.02 (1, 513) | <.01 [0, 1] | 0.896 |
| Time: session: latency change | 0.3 (1, 513) | <.01 [0, 1] | 0.585 |
| Time: session: latency baseline | 0.05 (1, 513) | <.01 [0, 1] | 0.823 |
| Time: arm: latency change | 0.01 (1, 513) | <.01 [0, 1] | 0.934 |
| Time: arm: latency baseline | 0.2 (1, 513) | <.01 [0, 1] | 0.656 |
| Session: arm: latency change | 1.09 (1, 514) | <.01 [0, 1] | 0.297 |
| Session: arm: latency baseline | 0.03 (1, 513) | <.01 [0, 1] | 0.867 |
| Time: session: arm: latency change | 1.06 (1, 513) | <.01 [0, 1] | 0.303 |
| Time: session: arm: latency baseline | 1.25 (1, 513) | <.01 [0, 1] | 0.265 |

*Supplementary Table 11 Electrical-evoked VP latency mixed-effects modelling*

The linear mixed effects modelling results for VP latency data are displayed for the a) HFS arm, b) control arm, and c) their relationship with MPS. HFS = High-frequency stimulation. VP = Vertex potential. N = 27, (female = 19).

The below nociceptive-evoked time-frequency plots display grand-averaged data across all participants for both the HFS (top) and control (bottom) arms, and for sessions 1 (left) and 2 (right) separately at each time point. Three types of data are considered: Raw FFT refers to the averaged ERSP data following Fast-Fourier Transform; SPRiNT peak fit refers to the averaged ERSP data following the removal of aperiodic components using Spectral Parameterisation Resolved in Time (Donoghue et al., 2020); Baseline removed refers to averaged ERSP data following the subtraction of the average baseline (-700 ms to -100 ms) for each frequency step (1 Hz) separately.

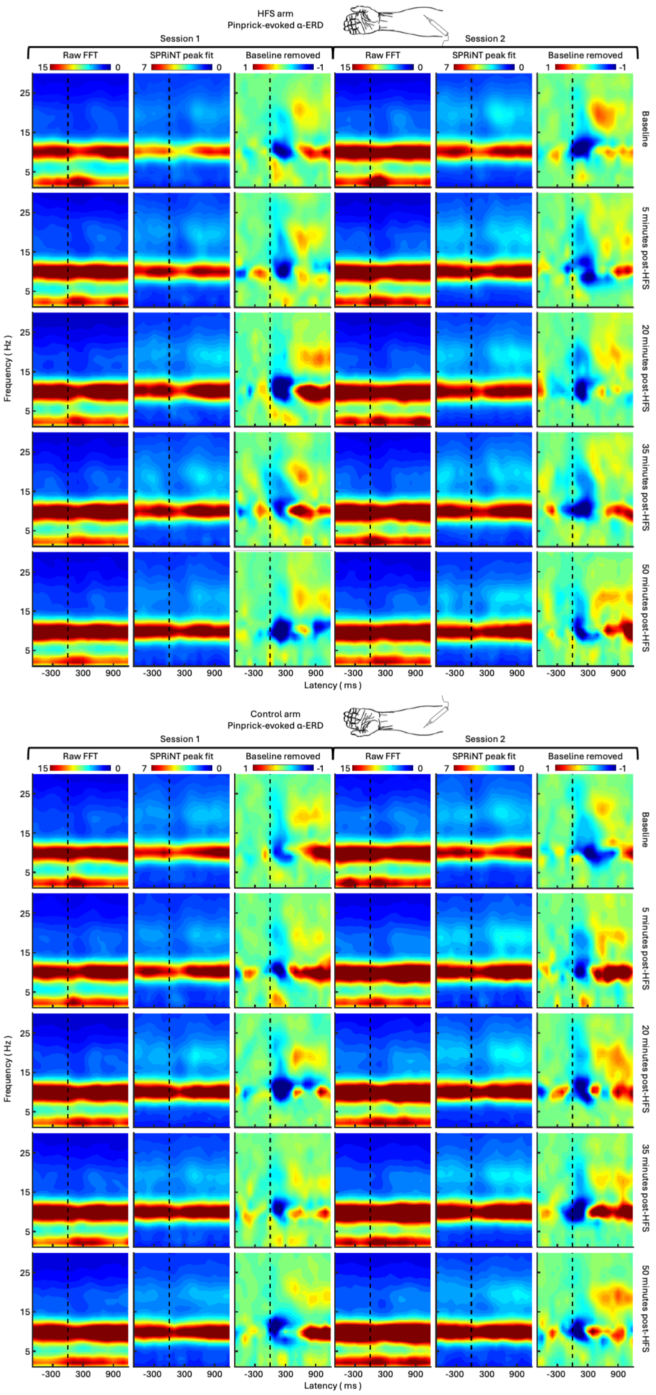


*Supplementary Figure 1 Pinprick-evoked α-ERD time-frequency plots*

ERSP time-frequency plots represent the grand-averaged α-ERD at contralateral central electrodes across all participants for both the HFS (top) and control (bottom) arms and sessions 1 (left) and 2 (right). α-ERD = Alpha-band event-related desynchronisation. HFS = High-frequency stimulation.

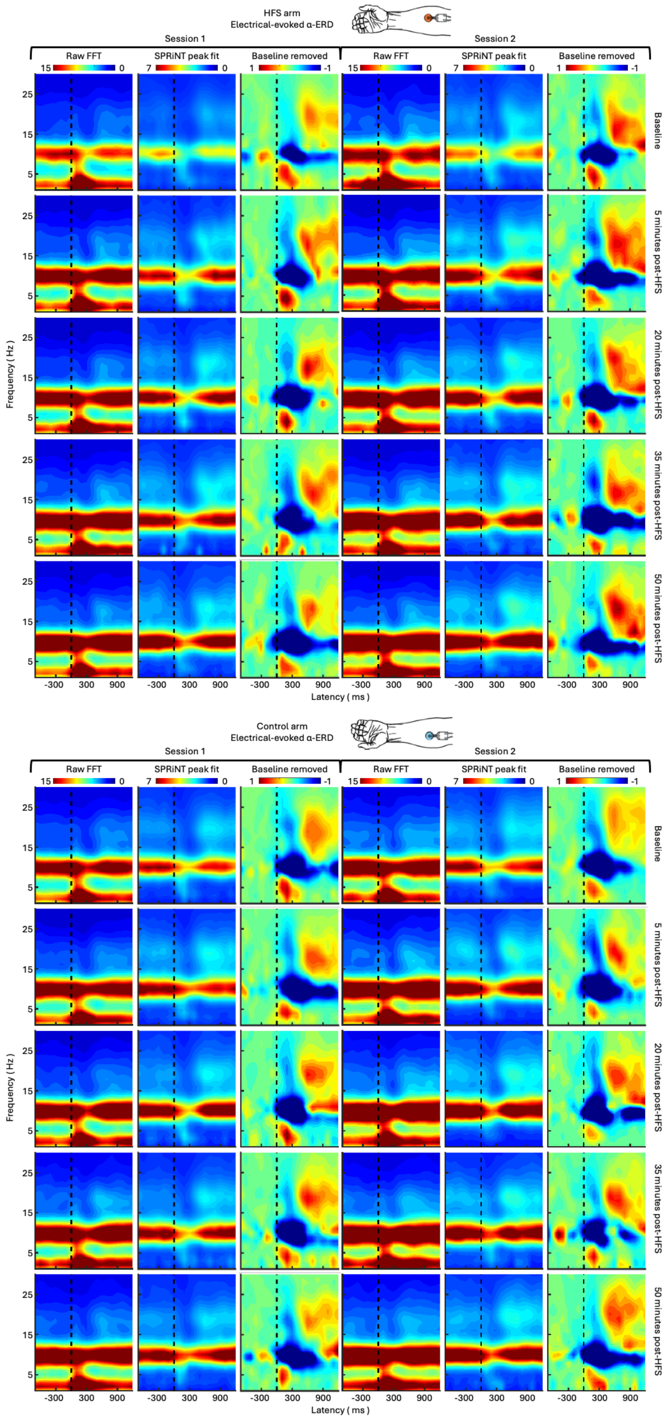


*Supplementary Figure 2 Electrical-evoked α-ERD time-frequency plots*

ERSP time-frequency plots represent the grand-averaged α-ERD at contralateral central electrodes across all participants for both the HFS (top) and control (bottom) arms and sessions 1 (left) and 2 (right). α-ERD = Alpha-band event-related desynchronisation. HFS = High-frequency stimulation.

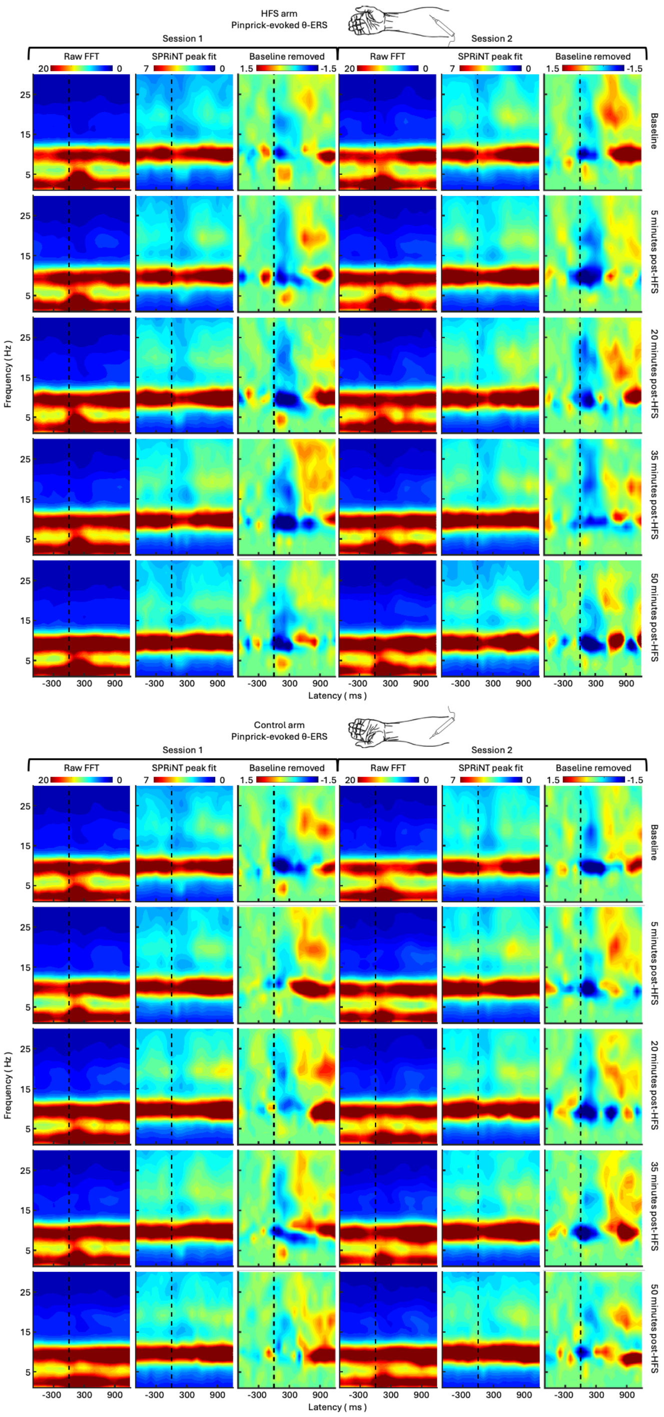


*Supplementary Figure 3 Pinprick-evoked θ-ERS time-frequency plots*

ERSP time-frequency plots represent the grand-averaged θ-ERS at the vertex electrode (Cz) across all participants for both the HFS (top) and control (bottom) arms and sessions 1 (left) and 2 (right). θ-ERS = Theta-band event-related synchronisation. HFS = High-frequency stimulation.

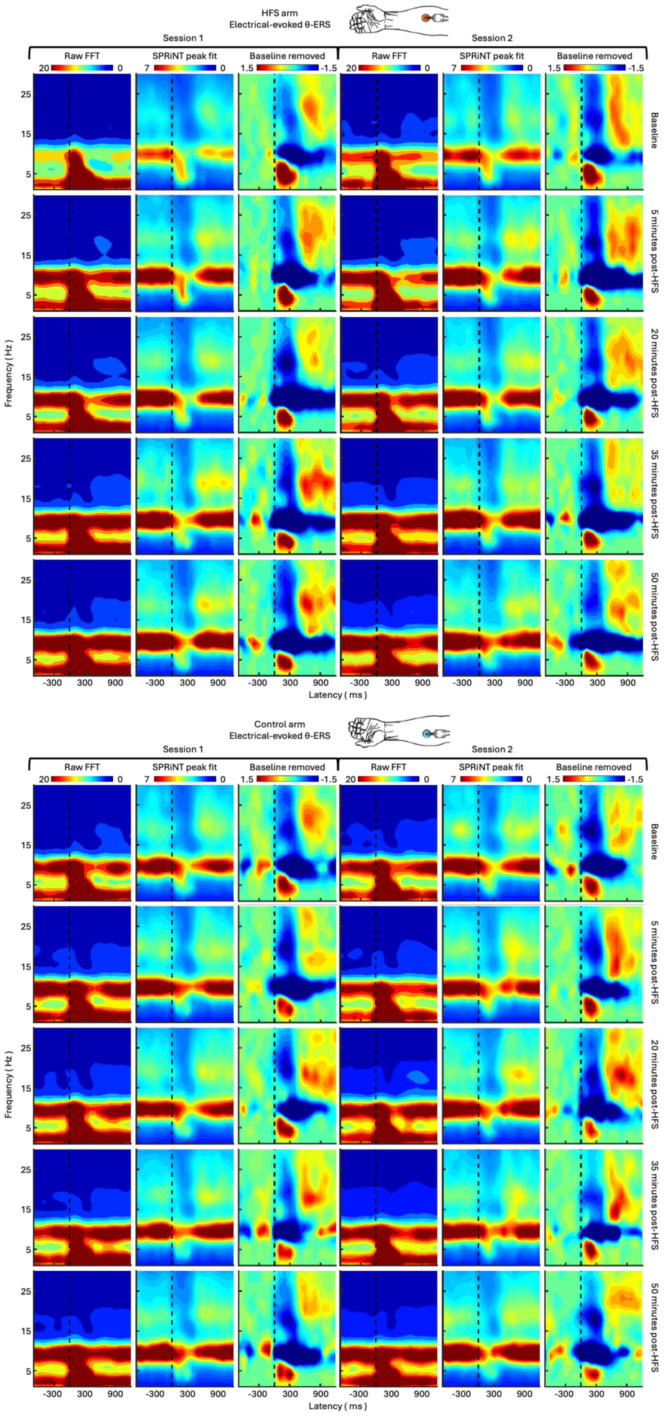


*Supplementary Figure 4 Electrical-evoked θ-ERS time-frequency plots*

ERSP time-frequency plots represent the grand-averaged θ-ERS at the vertex electrode (Cz) across all participants for both the HFS (top) and control (bottom) arms and sessions 1 (left) and 2 (right). θ-ERS = Theta-band event-related synchronisation. HFS = High-frequency stimulation.

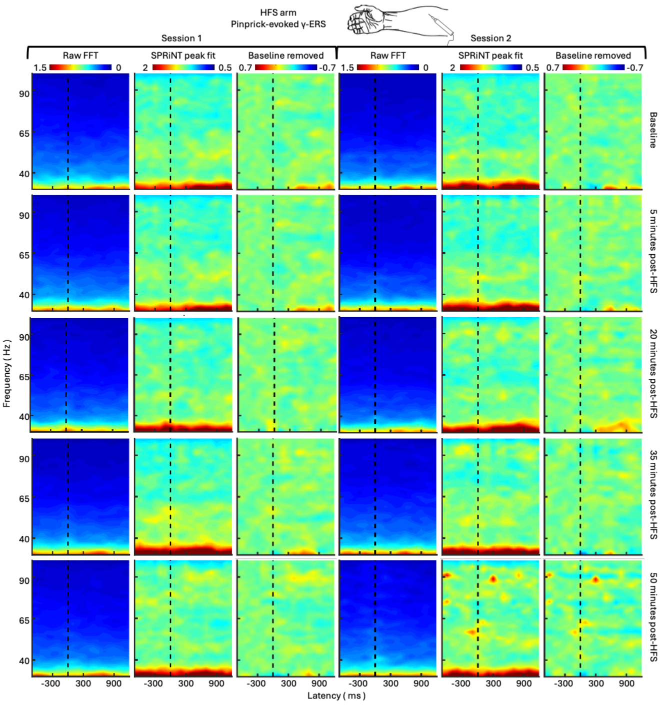


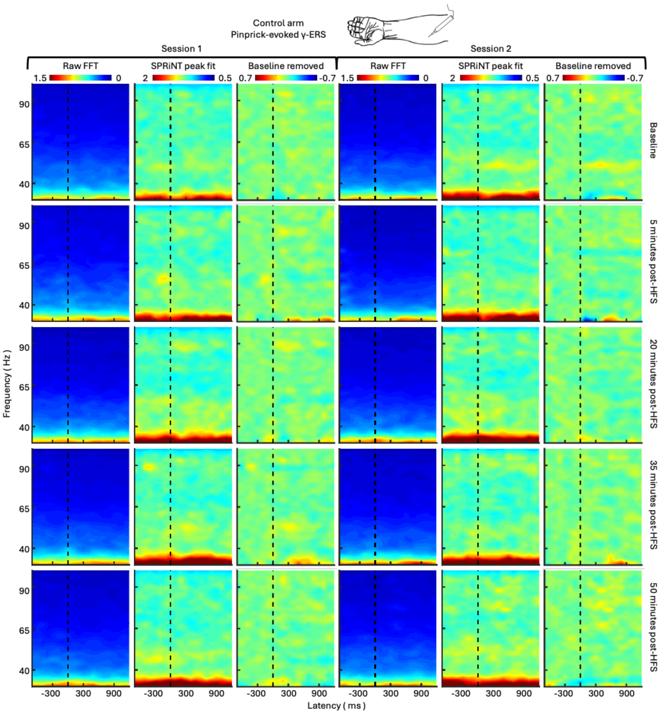


*Supplementary Figure 5 Pinprick-evoked γ-ERS time-frequency plots*

ERSP time-frequency plots represent the grand-averaged γ-ERS at the vertex electrode (Cz) across all participants for both the HFS (top) and control (bottom) arms and sessions 1 (left) and 2 (right). γ-ERS = Gamma-band event-related synchronisation. HFS = High-frequency stimulation.

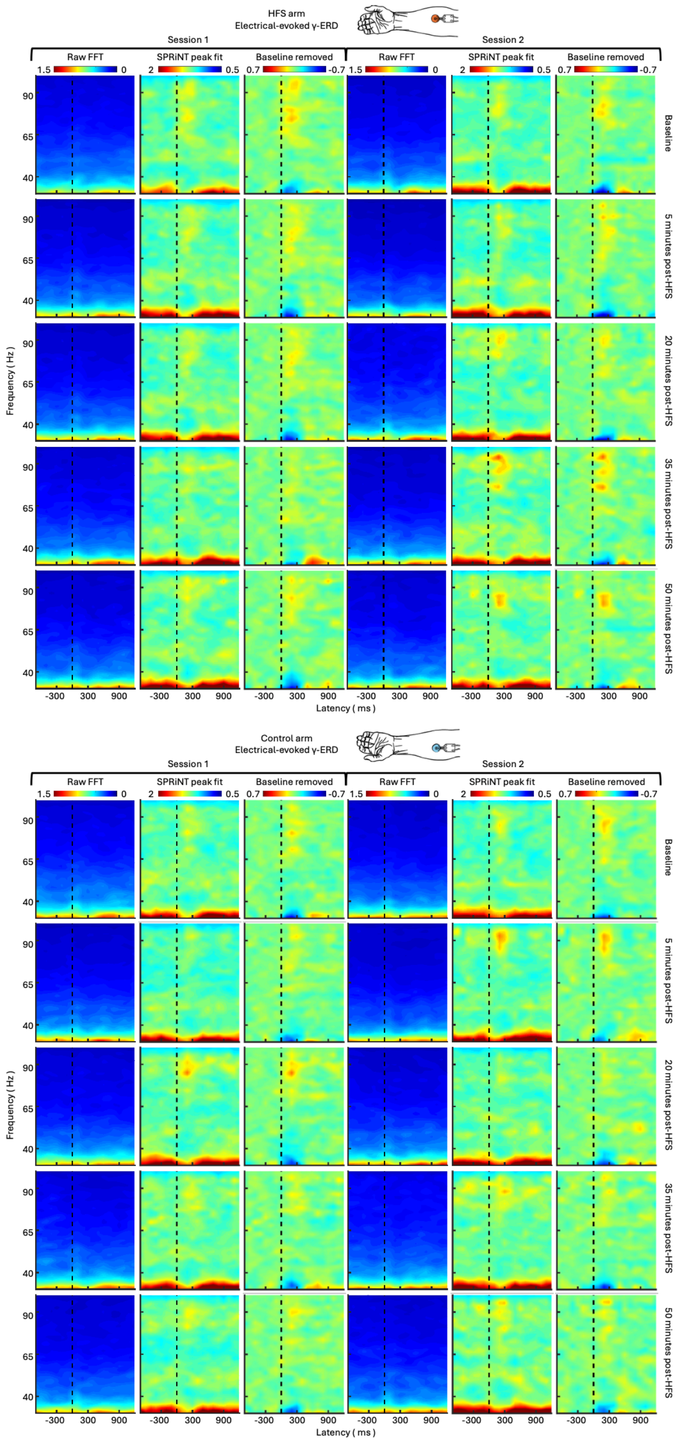


*Supplementary Figure 6 Electrical-evoked γ-ERS time-frequency plots*

ERSP time-frequency plots represent the grand-averaged γ-ERS at the vertex electrode (Cz) across all participants for both the HFS (top) and control (bottom) arms and sessions 1 (left) and 2 (right). γ-ERS = Gamma-band event-related synchronisation. HFS = High-frequency stimulation.
